# Mouse PAW: reverse-translating the FINGER multimodal lifestyle intervention enhances synaptic plasticity and cognition in adult wild type female mice

**DOI:** 10.1101/2024.09.23.614418

**Authors:** Vilma Alanko, Francesca Eroli, Ákos Végvári, Alina Solomon, Tobias Hartmann, Per Nilsson, Miia Kivipelto, Silvia Maioli, Anna Matton

**Author notes:** Correspondence to: Anna Matton and Silvia Maioli Full address: Karolinska vägen 37A, QA32, 171 64 Solna, Sweden Contact.

## Abstract

**Background:** Targeting multiple risk factors through preventive interventions can halt the onset or progression of dementia. The clinical efficacy of this approach was shown in the multimodal FINGER trial, yet knowledge of the underlying biological mechanisms is limited. In the current study, we applied for the first time a multimodal lifestyle intervention in mice to study effects on cognition and associated molecular pathways.

**Methods:** We have established a novel experimental model, the lifestyle intervention PAW (**P**revention of **A**lzheimer’s by a Multimodal Lifestyle Protocol – **W**orking-mechanisms for Memory Gains) in mice. In this model, we combined different modalities of intervention aiming to achieve synergistic effects. The animals were given access to running wheels (voluntary exercise), Fortasyn Connect (a medical food that slows disease progression in early Alzheimer’s disease), and they were subjected to cognitive training in an IntelliCage environment (PAW group). To separately investigate the effects of pharmacological vascular management as a preventive measure, another group of mice was given Atorvastatin and Enalapril (cholesterol and blood pressure lowering pharmaceuticals, respectively) dosed in the diet (Pharma group). Control mice were housed in normal conditions. We included 12 wild type C57BL/6J female mice 6.5 months of age per group. The full intervention lasted for 8 weeks. Blood pressure was measured at baseline and the end of the intervention. All mice underwent a battery of behavioural testing after the eight weeks of intervention. Hippocampi were dissected for proteomics analysis.

**Results:** During the intervention, the PAW group actively participated in the intervention by learning the different cognitive training programs and by using the running wheels. At the end of the intervention, the PAW group showed a lowering of blood pressure to a similar extent as in the Pharma group (systolic 158 mmHg to 139 mmHg, *p* = 0.003, and 163 mmHg to 136 mmHg, *p* = 0.042, respectively). Furthermore, the PAW group displayed a better short-term spatial working memory compared to the control group as assessed by the spontaneous alternation in the Y maze test (66.5% and 56.7%, respectively, *p* = 0.036). Proteomic analysis of the hippocampi and downstream bioinformatic analysis revealed that several pathways were upregulated in the PAW mice including synaptogenesis (Z = 4.27, *p* = 1.26E^–18^) and glutamate binding, activation of AMPA receptors and synaptic plasticity (Z = 3.74, *p* = 3.16E^–19^).

**Conclusions:** Our findings suggest that the PAW model effectively mimics the clinical effects of multimodal lifestyle dementia prevention, and further demonstrates the activation of hippocampal-specific molecular drivers of memory gains.

## Introduction

The global prevalence of Alzheimer’s disease and related dementias (ADRDs) is estimated to increase during the coming decades (Nichols et al., 2022). In addition to the ongoing research and development of drugs to modify and cure ADRDs (Budd Haeberlein et al., 2022; Cummings et al., 2023; Sims et al., 2023; van Dyck et al., 2023), prevention strategies have been considered to have great potential in reducing dementia risk (Livingston et al., 2024; World Health Organization, 2019). ADRDs are considered highly heterogeneous (Boyle et al., 2019; Kapasi et al., 2017; Mattsson-Carlgren et al., 2022) and many lifestyle-related factors influence the risk of cognitive impairment and progression into dementia (Livingston et al., 2024). Out of the fourteen potentially modifiable lifestyle-related risk factors identified by the *Lancet* Commission in 2024 (Livingston et al., 2024), risk factors such as hypertension, high cholesterol levels, physical inactivity, and diabetes can be more easily modifiable on an individual level in comparison to, for example, the other listed risk factors such as traumatic brain injury and air pollution. Still, lifestyle-related risk factors often co-occur (Kivipelto et al., 2006), resulting in complex risk profiles.

Considering the multifactorial nature of ADRDs and their risk factors, a preventive multimodal approach may be required to account for diverse disease mechanisms to simultaneously capture different risk profiles and possibly gain synergistic effects of the domains. In the field of dementia prevention, single-domain lifestyle changes where only one aspect of a lifestyle is modified, such as only increasing exercise or only changing diet, have been widely researched both in clinical and pre-clinical settings with mixed results (Alanko et al., 2022; Arnoldy et al., 2023; Hvid et al., 2021; Ji et al., 2021; Law et al., 2020; Townsend et al., 2023; Yu et al., 2020). The Finnish Geriatric Intervention Study to Prevent Cognitive Impairment and Disability (FINGER) was the first randomised-controlled trial showing that a multimodal lifestyle intervention has the potential to decrease the risk of dementia, resulting in personal health and societal benefits (Lehtisalo et al., 2022; Marengoni et al., 2018; Ngandu et al., 2015; Wimo et al., 2023). The FINGER lifestyle intervention included the combination of five domains: exercise, diet, cognitive training, social activities, and vascular risk monitoring (Ngandu et al., 2015). In the trial, a reduction in dementia risk in the intervention group was associated with decreased loss in hippocampal volume (Stephen et al., 2021), yet it is still unknown what are the molecular working mechanisms behind the beneficial cognitive effects of multimodal interventions.

To gain a mechanistic understanding of how a multimodal lifestyle intervention influences the brain, we reverse-translated the FINGER model to mice and developed the **P**revention of **A**lzheimer’s by a Multimodal Lifestyle Protocol – **W**orking-mechanisms for Memory Gains (Mouse-PAW). The impact of this intervention on molecular, cognitive, and physical aspects was established by applying the protocol to adult wild type mice. We designed an experimental setup using the IntelliCage (IC) system (Kiryk et al., 2020) connected to running wheels where mice received cognitive training, physical activity, and diet containing Fortasyn Connect. Fortasyn is a medical food with a brain health-promoting combination of nutrients docosahexaenoic acid and eicosapentaenoic acid; uridine monophosphate; choline; vitamins B12, B6, C, E, and folic acid; phospholipids; and selenium (Van Wijk et al., 2014). In the LipiDiDiet clinical trial, prodromal AD patients receiving Fortasyn showed slower cognitive decline as well as reduced hippocampal atrophy compared to placebo controls (Soininen et al., 2017, 2021). Moreover, in the proof-of-concept study MIND-AD_mini_, prodromal AD patients showed good adherence to a combination of the FINGER protocol and Fortasyn medical food, which resulted in both cognitive and cardiovascular benefits (Sindi et al., 2022; Thunborg et al., 2024). In essence, in this new model, we combined different modalities simultaneously aiming to achieve synergistic intervention effects. In addition, we included a vascular risk management group that underwent a pharmacological intervention receiving a combination of atorvastatin and enalapril – the most prescribed compounds in Sweden in their respective drug class (Wastesson et al., 2018) – to compare with the lifestyle intervention. Atorvastatin is a hydroxy-methyl-glutaryl-coenzyme A reductase inhibitor used to lower blood LDL-cholesterol; enalapril is an angiotensin-converting enzyme inhibitor, used to treat hypertension. A transition from studying the effects of only single-domain interventions to studying multimodal lifestyle changes in animal models is necessary to learn about mechanisms that underlie lifestyle-induced dementia prevention initiatives.

## Materials & methods

### Ethical statement

All procedures on mice were performed following the national animal care and use guidelines of Sweden and following the 3R principles. The study has been approved by the Swedish Board of Agriculture (Dnr 18252-2021, Stockholm ethical committee) and the local committee at Karolinska Institutet. All possible efforts were made to minimise the suffering and distress of the animals.

### Animals and general animal welfare

We used female C57BL/6JRj mice (Janvier Labs, France). At 6 months of age, a total of 36 mice were randomly distributed into three groups of 12 mice. The three groups were constituted as follows: 1) The Prevention of Alzheimer’s Working-mechanisms (PAW) group, which received the multimodal lifestyle intervention comprising cognitive training, physical activity, and brain health-promoting diet; 2) The Pharma group, which received a pharmacological intervention; And 3) the Control group that did not undergo any intervention (**Table 1**). The intervention lasted for a total of eight weeks (**Fig. 1A**). At the start of the intervention, mice were 6.5 months of age.

**Figure 1.**
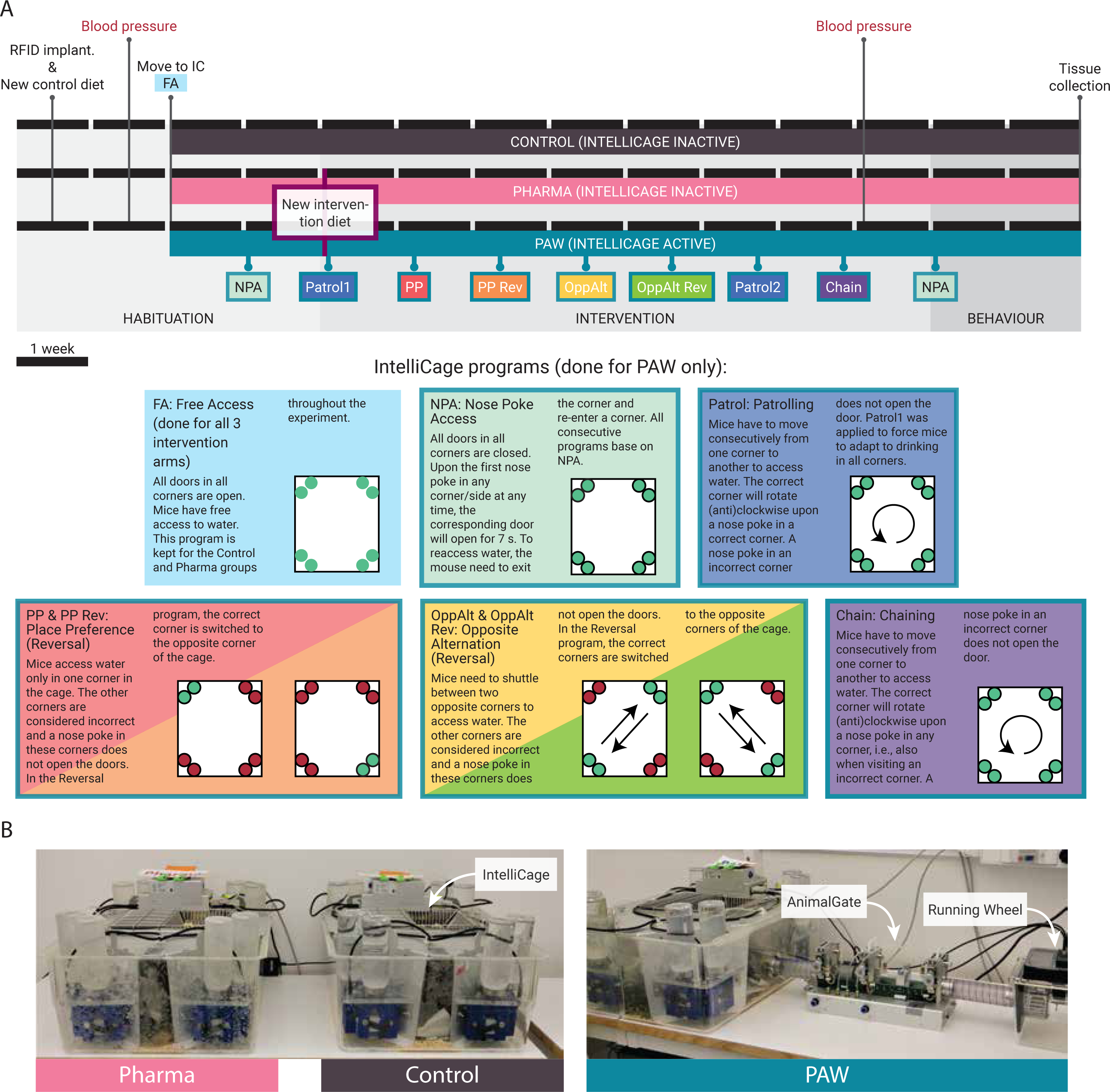
Schematic overview of the study protocol. (**A**) Timeline depicting the experimental phases of the Mouse PAW study including descriptions of the different cognitive training programs. All three groups underwent the same experimental procedures except for the intervention diet (PAW and Pharma groups only) and the IntelliCage programs (PAW only). The habituation period lasted for four weeks, intervention for eight weeks, and behavioural testing for two weeks. (**B**) Pictures presenting the housing conditions for all three study groups. The three groups were housed in their respective IntelliCages. The cage that housed the PAW group was additionally connected to two AnimalGates that were further attached to running wheels (other gate and wheel on the opposite side of the cage).

**Table 1.** Summary of intervention domains in the three different groups.

| Group | Animals (N) | Intervention domains |  |  |  |  |
| --- | --- | --- | --- | --- | --- | --- |
|  |  | Diet | Physical activity | Cognitive training | Vascular risk management | Social interaction |
| PAW | 12 | Fortasyn Connect | Voluntary wheel running | IntelliCage programs | None | Yes |
| Pharma | 12 | Base diet | None | None | Atorvastatin and Enalapril | Yes |
| Control | 12 | Base diet | None | None | None | Yes |

Before moving to the IC system, all mice were housed in individually ventilated cages (IVC) with 3–5 mice per cage, with water and regular chow provided *ad libitum*. IVCs had standard enrichment with cardboard tunnels, tissue paper, shredded cardboard, and wooden sticks on chip bedding. Once transferred to the ICs, the enrichment and bedding were maintained similar to the one in IVCs, with the addition of three red plastic houses in the middle of the cage. In the ICs, the water was changed weekly, while bedding changes and cage cleaning were performed biweekly. Mice were weighed weekly or when needed to monitor their health and individual drinking activity was monitored daily throughout the study by assessing lick numbers. At any moment, if a mouse had performed less than 50 licks/day, they were taken out from the cage and placed in an IVC with a water bottle for up to 30 min to avoid dehydration. All groups of mice learned how to drink in the IC within the first week of living in the cage and did not show signs of dehydration (skin tents).

Mice were housed under a 12-hour dark/light cycle from 7 AM to 7 PM, at 20–22℃ with 45–50% humidity throughout the study. Handling and lifting of mice were performed exclusively using a tube – a refining method previously described by Hurst & West (Hurst & West, 2010). All activities in the room were performed by the same female researcher (VA).

### Diet

Before the intervention started, all mice were habituated to a semi-synthetic control diet (ssniff Spezialdiäten, Germany) for three and a half weeks. The Control group continued with this diet throughout the intervention period, while the PAW and Pharma groups received their respective isocaloric diets. The diet specifications are presented in **Supplementary Table 1**. Diets were stored at –20°C, thawed pellets were replaced every three days and food intake was registered per cage. Food was provided *ad libitum* to all three groups throughout the study.

#### The dietary domain of PAW

At the beginning of the intervention period, the control diet was replaced with the Fortasyn diet (Danone Nutricia Research, The Netherlands; ssniff Spezialdiäten). The diet specifications are listed in **Supplementary Table 1**.

#### The pharmacological (Pharma) intervention

As a vascular preventive intervention, mice in the Pharma group received a lipid-lowering and an antihypertensive drug dosed in the diet. In detail, the diet (ssniff Spezialdiäten) was prepared with Atorvastatin (10493, Cayman Chemical, USA) and Enalapril (16041, Cayman Chemical) to final concentrations of 128.6 mg/kg and 85.7 mg/kg, respectively (**Supplementary Table 1**). The final average daily doses (expected dose in parentheses) for the animals based on average food intake and body weight during the study were 14 (15) mg/kg/day for Atorvastatin and 9.5 (10) mg/kg/day for Enalapril. The dosages have previously been tested in rodents without any unwanted side effects and they were selected to match the human therapeutic range (Trebicka et al., 2007; Vandresen-Filho et al., 2015; Yamada et al., 2010).

### The IntelliCage (IC) system

The IC system (TSE Systems, Germany) is a fully automated home cage-like monitoring system (Kiryk et al., 2020). This system is designed to evaluate the behaviour of mice living in social groups, up to 16 animals per cage, largely without human interference for long periods. The IC apparatus is fitted in a standard large, not individually ventilated, rodent cage of size 21 x 55 x 39 cm (h x l x w). There are four recording chambers, one in each corner (hereafter referred to as corner) of the apparatus that fit one mouse at a time. Mice access the corners through a ring-shaped tube containing an antenna for mouse identification. The antenna reads the radio-frequency identification (RFID) transponders that have been implanted subcutaneously in mice. The presence of a mouse in the corner is detected by the RFID and a temperature sensor. Within the triangular-shaped corner, ports to two respective water bottles are located on both the left and right walls from the entrance. The ports can be closed by doors. Nose pokes to the ports and licks are detected by their respective sensors. The ICs are connected to a PC from where they are operated using the IC software (v. 3.6.3, TSE Systems). We used three ICs where each cage housed one of the three groups (**Fig. 1B**). The IC system was not cleaned during the study to avoid loss of smell cues.

#### Habituation to the IntelliCage system

The habituation period anticipating the intervention period lasted for four weeks in total (see schematic in **Figure 1A**). All mice were first habituated to the study room by living in the IVCs placed on tables and without filters for two weeks. During the first week, mice were subcutaneously implanted with the RFID transponders (Uno Micro ID/12 ISO Transponder, Uno Life Science, The Netherlands) under 2% isoflurane (Attane vet) inhalation anaesthesia according to manufacturer’s instructions (TSE Systems). At the beginning of the third week, mice were transferred from their IVCs to their respective ICs. The transfer was performed at the beginning of the dark phase. During the first week in the IC, all three cages were on a Free Access (FA) program. During FA, all doors in the corners were open for mice to learn where they receive water (**Fig. 1A**). The Control and Pharma groups continued the FA program for the whole study, thus having *ad libitum* access to water.

#### The cognitive domain of PAW

During the second week of habituation in the IC, the PAW group was adapted to Nose Poke Access (NPA; **Fig. 1A**). During NPA, all the doors to the water bottles were closed. When a mouse visited a corner, the first nose poke to either side opened the corresponding door for 7 seconds, allowing the mouse to drink. After 7 seconds, the door closed and to re-open the door, the mouse needed to exit and re-enter a corner. The NPA program was the base for the more complex programs where a nose poke to a correct corner granted access to water, while nose pokes to incorrect corners did not open the doors.

The cognitive training was based on positive reinforcement, with water given to mice as a reward for learning different IC programs. To avoid unnecessary stress for the animals, the water could be accessed at any time, i.e., there was no temporal restriction of water (no drinking sessions that are often used in ICs). During the eight-week intervention, seven cognitive training programs were run in the IC in the following order: Patrolling 1 (Patrol1), Place Preference (PP), Place Preference Reversal (PP Rev), Opposite Alternation (OppAlt), Opposite Alternation Reversal (OppAlt Rev), Patrolling 2 (Patrol), and Chaining (Chain). The different programs are described and visualised in **Figure 1A**. Each program ran for six consecutive nights after which the mice could rest for two consecutive nights on the NPA program before the next learning program started. The first Patrolling program, Patrol1, was performed to motivate mice to visit and drink from all four corners to adjust them to actively alternate between corners. All mice were able to perform all seven cognitive programs without adjustments. All animal handling and water changes were done during the resting days to avoid disturbances in learning.

#### The physical activity domain of PAW

The cage that housed the PAW group was equipped with two AnimalGates (AG; TSE Systems), located on opposite sides of the cage, and connected to running wheels (RW; TSE Systems; **Fig. 1B**). RWs had a diameter of 117 mm and even spaced metal rods. Similarly to the IC system, also the AGs and the RWs are automated and operated through the IC software. The PAW group was introduced to the AGs and RWs three days before the intervention started. AGs were, for the first days, kept completely open to allow mice to habituate to the gates and RWs. Later, AGs were set to “Training” mode to familiarise the mice with opening and closing the AG doors. AG entries and running activities during this habituation phase have not been considered in the analysis. Three days into the intervention period, all mice had visited the RWs, and thus the AGs were switched from “Training” to “EasySingle” mode. In the “EasySingle” mode, only one mouse at a time was allowed to visit the RW, enabling tracking of the physical activity parameters of individual mice. The RWs registered the running velocity, duration, and distance. The “EasySingle” mode was kept until the end of the study.

### Measurements and sampling

#### Blood pressure

Blood pressure (BP) was determined at baseline (second week of habituation) and at the end of the intervention (last week of intervention; **Fig. 1A**). For each time point, measurements were divided on three consecutive days to ensure a similar time of the day (between 2 and 4 PM) for all animals, thus minimising the effect of timing on BP between groups (Zhang et al., 2021). BP was measured using the CODA non-invasive tail-cuff system according to the manufacturer’s instructions (CODA-HT4, Kent Scientific Corporation, USA). Mice were acclimated to the experimental room for a minimum of 45 min. Before the measurement, mice were habituated to the restrainer while simultaneously warming up on a heating platform for a minimum of 5 min. Four mice from each group were measured per day. The first five measurements of each mouse were regarded as habituation to the cuffs and were thus excluded from the analysis. An N of 12, 10, and 8 mice in the Control, PAW, and Pharma groups, respectively, had a minimum of seven accepted measurements on both time points. Values deviating 3 standard deviations (SD) from the mean for an individual animal were considered outliers and excluded. The median value of all accepted cycles was considered as the level for systolic, diastolic, and mean BP as well as the heart rate for each animal.

#### Behavioural testing

The behavioural battery was performed in the last part of the intervention (**Fig. 1A**). This battery included classical behavioural tests, run in the following order: Open Field (OF), Elevated Plus Maze (EPM), Y maze, Novel Object Recognition (NOR), and Fear Conditioning (FCond). We used the same instrumentation and followed the same test procedures as described in previous publications from the group, with minor differences for the objects used during the NOR test (Alanko et al., 2023; Eroli et al., 2020). For NOR, two different objects were used: an oval-shaped (egg, 6 cm wide x 5 cm high) object with light red colour and a taller (T25 flask, 5 cm wide x 10 cm high) object filled with sand. During day 2, half of the mice were habituated to explore the egg object and half of the mice to the flask object. On day 3, those mice that had the egg as the familiar object on day 2 got the flask as a novel object and those mice that had the flask as the familiar object on day 2 got the egg as a novel object. Before each behavioural test, mice were moved from their ICs into IVCs, brought to the experimental room and left there to acclimatise for a minimum of 1 h before starting the procedure. Activity during the OF test was acquired using the TSE ActiMot apparatus and software (TSE Systems). EPM, Y maze, and NOR were performed on arenas from Ugo Basile (Italy) and data were acquired using the video-tracking software Ethovision XT 17 (Noldus Information Technology, The Netherlands). FCond data were acquired using the TSE Multi Conditioning System (TSE Systems). All behavioural tests were run between 8:30 AM and 12:30 PM (or 3:00 PM in case of OF) by female researchers (VA and FE).

#### Tissue collection and preparation

At the end of the study, mice were sacrificed by cervical dislocation. Hippocampi were dissected, snap-frozen on dry ice, and stored at –80°C until further use. For protein extraction, tissues were first homogenised in a Milli-Q water buffer containing 1% protease inhibitors (P8340, Sigma, USA) and 1% phosphatase inhibitors (P0044, Sigma) with a manual homogeniser. Next, 250 µl of RIPA protein extraction buffer (89900, Thermo Fischer Scientific, USA) containing 1% protease inhibitors and 1% phosphatase inhibitors was mixed with 55 µl of tissue homogenate. Samples were kept on ice and after lysis samples were centrifuged at 10,000 rcf for 5 min at 4°C. The supernatant was collected for protein analyses.

#### Mass spectrometry analysis/protein determination

After tissue sample preparation, lysates were supplemented with a 4-fold volume of chilled acetone and kept at –20°C overnight. Following spun down at 14,000 g at 4°C for 20 min acetone was removed and the protein pellet was dried before being solubilised in 10 µl of 8M urea (U5378, Sigma), sonicated in a water bath for 10 min, supplemented with 80 µl of 50 mM Tris-HCl, pH 8.5 and sonicated again for 10 min. The protein concentrations were determined by BCA assay (23225, Thermo Fischer Scientific). A volume of lysate corresponding to 25 µg of protein was taken and supplemented with Tris-HCl buffer up to 100 µl. Proteins were reduced by adding 2.5 µl of 250 mM dithiothreitol (D0632, Sigma) and incubated at 45°C for 37 min while shaking at 400 rpm on a block heater. Alkylation was performed with addition of 5 µl of 500 mM iodoacetamide (I6125, Sigma) at room temperature (RT) for 30 min in the dark. Then 0.5 µg of sequencing grade modified trypsin (V5111, Promega, USA) was added to the samples and incubated for 16 h at 37°C. The digestion was stopped with 6 µl cc. formic acid (FA; 1.00264.1000, Sigma), incubating the solutions at RT for 5 min. The sample was cleaned on a C18 Hypersep plate with 40 µl bed volume (60300-525, Thermo Fisher Scientific), and dried using a vacuum concentrator (Concentrator Plus, Eppendorf, Germany).

Samples were labelled with TMTpro (A52045, Thermo Fisher Scientific) isobaric reagents in two sets. Peptides were solubilized in 70 µl of 50 mM triethylammonium bicarbonate (90114, Thermo Fisher Scientific) and mixed with 100 µg TMTpro reagent in anhydrous acetonitrile (ACN; 364100010, Thermo Fisher Scientific) and incubated for 2 h at RT. The unreacted reagents were quenched with 11 µl of hydroxyamine (90115, Thermo Fisher Scientific) for 15 at RT. Biological samples were then combined, dried in vacuum and cleaned on a C18 Hypersep plate.

The combined TMTpro-labelled biological replicates were fractionated by high-pH reversed-phase after dissolving in 50 µl of 20 mM ammonium hydroxide and were loaded onto an Acquity bridged ethyl hybrid C18 UPLC column (2.1 mm inner diameter × 150 mm, 1.7 μm particle size, Waters), and profiled with a linear gradient of 5–60% 20 mM ammonium hydroxide (205840010, Acros, Thermo Fisher Scientific) in ACN (pH 9.0) over 48 min, at a flow rate of 200 µl/min. The chromatographic performance was monitored with a UV detector (Ultimate 3000 UPLC, Thermo Fisher Scientific) at 214 nm. Fractions were collected at 30 s intervals into a 96-well plate and combined in 12 samples concatenating 8-8 fractions representing peak peptide elution.

The peptide fractions were in solvent A and approximately 2 µg samples were injected on a 50 cm long EASY-Spray C18 column (Thermo Fisher Scientific) connected to an Ultimate 3000 nanoUPLC system (Thermo Fisher Scientific) using a 90 min long gradient: 4–26% of solvent B (98% ACN, 0.1% FA) in 90 min, 26–95% in 5 min, and 95% of solvent B for 5 min at a flow rate of 300 nl/min. Mass spectra were acquired Q Exactive HF hybrid quadrupole-Orbitrap mass spectrometer (Thermo Fisher Scientific) ranging from m/z 350 to 1800 at a resolution of R=120,000 (at m/z 200) targeting 5x106 ions for maximum injection time of 100 ms, followed by data-dependent higher-energy collisional dissociation (HCD) fragmentations of top 17 precursor ions with a charge state 2+ to 7+, using 45 s dynamic exclusion. The tandem mass spectra were acquired with a resolution of R=60,000, targeting 2x105 ions for a maximum injection time of 54 ms, setting quadrupole isolation width to 1.4 Th and normalized collision energy to 34%.

Acquired raw data files were analysed using Proteome Discoverer v3.0 (Thermo Fisher Scientific) with the SequestHT search engine against the mouse protein database (UniProt, downloaded on 27 March 2024). A maximum of two missed cleavage sites were allowed for full tryptic digestion, while setting the precursor and the fragment ion mass tolerance to 10 ppm and 0.02 Da, respectively. Carbamidomethylation of cysteine was specified as a fixed modification. Oxidation on methionine, deamidation of asparagine and glutamine, TMTpro (+304.207 Da) of lysine and peptide N-termini were set as dynamic modifications. Initial search results were filtered with 5% FDR using the Percolator node in Proteome Discoverer. Quantification was based on the reporter ion intensities.

#### Ingenuity Pathway Analysis (IPA)

Hippocampal proteomics data were analysed using the QIAGEN IPA (QIAGEN Inc., https://digitalinsights.qiagen.com/IPA) to identify significantly enriched pathways in the intervention groups compared to the Control group. Proteins that had an FDR of less than 5% and a fold change of more or less than 1.25 or –1.25, respectively, were considered for the canonical pathways core analysis. Pathways with an absolute Z-score of at least 2 and a *p*-value of less than 0.05 were regarded as significant.

### Statistical analysis

Data handling, statistical analyses, and visualisation were performed using the R software (version 4.3.0) with the following packages: vroom, tidyverse, lubridate, scales, ggpubr, ggbeeswarm, rstatix, plotrix, corrplot. Differences between groups were analysed with one-way ANOVA, two- or three-way mixed model ANOVA or Kruskal Wallis rank sum test, when appropriate. For post hoc tests, paired or unpaired two-tailed Student’s *t*-tests or Wilcoxon tests with Bonferroni correction were performed, when appropriate. The distribution of data was assessed with the Shapiro-Wilk normality test and homogeneity of variance with Levene’s test. Correlations were analysed using Spearman’s correlation. The area under the curve (AUC) for individual mice’s learning curves in the different cognitive programs was calculated in R using the functions integrate (stats) and approxfun (stats). The null hypotheses were rejected with *p*-values ≤ 0.05.

## Results

### Health and activity parameters over the intervention period

The PAW and Pharma interventions were generally well-tolerated, and all the mice completed the full study protocol. No mice died or had to be sacrificed prematurely due to illness. We observed some differences between groups in general health parameters. The body weight changes during the intervention were dependent on both the intervention group and time point (two-way mixed model ANOVA, *F*_(30, 495)_ = 8.801, *p* < 0.0001; **Fig. 2A**). At baseline, there were no statistically significant differences in body weights between groups. After two weeks of intervention, both PAW (28.9±2.6 g) and Control (27.9±2.2 g) groups gained significantly more weight compared to the Pharma group (25.8±1.7 g) (post hoc comparisons between groups, two-tailed Student’s *t*-test with Bonferroni correction, *p* = 0.008 and *p* = 0.046, respectively; **Fig. 2A**). On the last weighing, at the time of behavioural testing, PAW mice (32.2±2.2 g) were significantly heavier than both Control (29.1±2.2 g) and Pharma (26.0±2.8 g) mice (*p* = 0.006 and *p* < 0.0001, respectively), and the Control group kept showing increased body weight compared to the Pharma group (*p* = 0.023).

**Figure 2.**
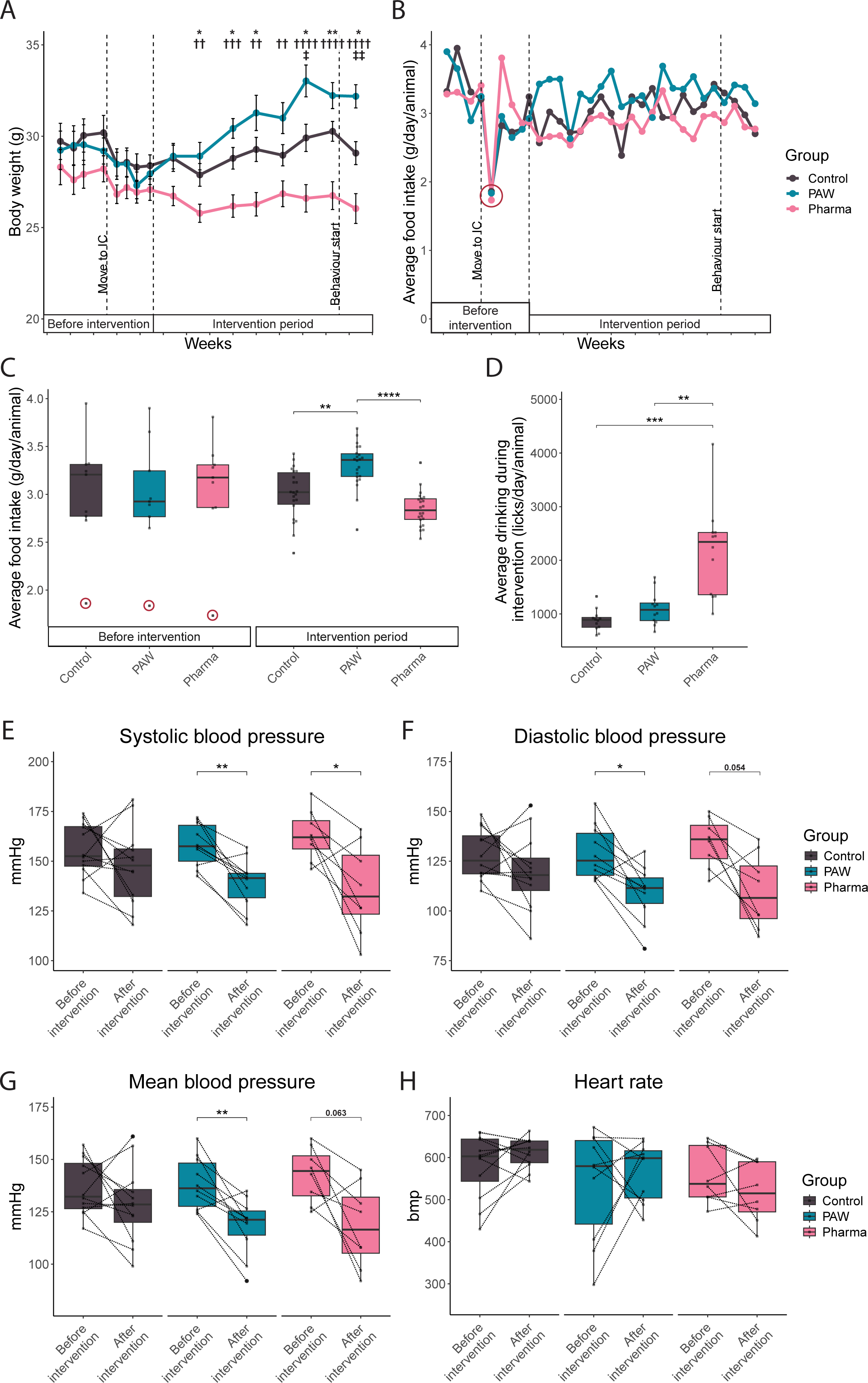
General health parameters during the study. (**A**) Body weight and (**B**) food intake development throughout the study period. The first dashed line indicates the time point when mice were moved to IntelliCages (IC), the second when the intervention started, and the third when behavioural testing was initiated. (**C**) Summary of food intake before or during the intervention. Each data point corresponds to a data point presented in (**B**). (**B, C**) Data points circulated in red correspond to the first time point food intake was recorded in the IC. (**D**) Drinking activity was measured as the average lick number performed per animal per day. (**E**) Systolic, (**F**) diastolic, and (**G**) mean blood pressure and (**H**) heart rate (beats per minute, bpm) measured before the intervention start and at the end of the intervention. Data points from the animals that had repeated measurements on both time points were included. (**A**) Data presented as mean±SEM. (**C–H**) Data presented as median (thick black line), 25^th^ and 75^th^ percentile (box) and 1.5 x interquartile range (whiskers). (**A**) * Control vs. Pharma, † PAW vs. Pharma, ‡ Control vs. PAW. (**A–H**) */ ‡ *p* < 0.05, **/††/‡‡ *p* < 0.01, ***/††† *p* < 0.001, ****/†††† *p* < 0.0001

There were no differences in food intake between groups during the habituation period (**Fig. 2B, C**), however, as expected, we registered a transient drop in food consumption and body weight for all three groups at the first time point after moving into the ICs (**Fig. 2A–C**). It is worth mentioning that since each IC system housed 12 mice together, the food intake was not registered individually but as the average food intake (g) per animal per day. On average, during the intervention, PAW mice ate 3.3±0.2 g/day/animal, which was more than what the Control mice (3.0±0.3 g/day/animal) and Pharma mice (2.9±0.2 g/day/animal) ate (one-way ANOVA for intervention food intake *F*_(2, 63)_ = 22.21, *p* < 0.0001, post hoc two-tailed Student’s *t*-test with Bonferroni correction, PAW vs. Control *p* = 0.001, PAW vs. Pharma *p* > 0.0001, Control vs. Pharma *p* = 0.072; **Fig. 2C**).

During the intervention, the drinking activity, evaluated by the number of licks each mouse performed during a day, was significantly different between groups (Kruskal-Wallis rank sum test, *p* < 0.0001; **Fig. 2D**). For the Pharma group, the median daily lick number during this period was 2342 (IQR 1160), which was significantly higher than for PAW (1077 [328], post hoc two-tailed Wilcoxon test with Bonferroni correction, *p* = 0.004) and the Control group (888 [182], *p* = 0.0008). The baseline drinking activity was not analysed, as drinking activity during the habituation period depended on the individual learning.

BPs were similar between all three groups both before and after the intervention for systolic (two-way mixed model ANOVA, *F*_(2, 27)_ = 0.18, *p* = 0.835 for one-way group effect) diastolic (*F* = 0.45, *p* = 0.645), and mean BP (*F* = 0.37, *p* = 0.693; **Fig. 2E–G**). Similarly, the heart rates did not significantly differ between the groups (*F* = 2.78, *p* = 0.080; **Fig. 2H**). However, we observed a significant one-way time effect for the BPs (systolic *F* = 24.54, *p* < 0.0001; diastolic *F* = 23.53, *p* < 0.0001; mean *F* = 23.11, *p* < 0.0001). The PAW intervention reduced systolic BP from an average of 158±11 mmHg at baseline to 139±13 mmHg at the end of the intervention (post hoc paired Student’s *t*-test with Bonferroni correction, *p* = 0.003), and the Pharma intervention reduced systolic BP from 163±13 mmHg to 136±22 (*p* = 0.042). Similar reductions (although not significant for the Pharma group) were observed for both diastolic (PAW 129±13 mmHg to 109±14 mmHg, *p* = 0.012, Pharma 134±12 mmHg to 110±19 mmHg, *p* = 0.054) and mean BP (PAW 139±12 mmHg to 118±14 mmHg, *p* = 0.006, Pharma 143±13 mmHg to 118±20 mmHg, *p* = 0.063).

At baseline, all groups showed similar activity in the ICs measured as the number of visits during the first hour in the cage (**Supplementary Fig. 1A**), and during the first 24 hours in the cage (**Supplementary Fig. 1B**). The average number of visits during daytime and nighttime and duration of corner visits were similar between the groups during the period before the intervention (**Supplementary Fig. 1C**, D). During the intervention period, the PAW group performed more visits (**Supplementary Fig. 1C**) while the Pharma group spent significantly more time in the corners (**Supplementary Fig. 1D**). The findings are described in more detail in the **Supplementary Material**.

### Activity and learning during the PAW intervention

The PAW group was subjected to a battery of seven cognitive training programs in the IC (see schematic illustration in **Fig. 1A**). **Figure 3A** summarises the average learning curves relative to the individual programs of cognitive training in the IC. Starting with the PP program, the mice performed below the chance level of 75% already during the first night (mean±SEM 64.5±3.8%). During each consecutive night, on average, mice improved their performance until the last day of the training period (one-way repeated measures ANOVA, *F*_(5, 55)_ = 15.1, *p* < 0.0001, post hoc paired two-tailed Student’s *t*-test with Bonferroni correction, Day 1 vs. Day 6, *p* = 0.002). In the following PP Rev program, the mice started the first night on a similar level to the PP (61.1±3.2%) and significantly corrected their performance during the training period (*F* = 13.5, *p* < 0.0001). They reduced their error rate significantly already from the first to the second day (36.9±3.5%, *p* = 0.0001) and remained on the same level until the last day (34.5±3.0%, Day 1 vs. Day 6, *p* < 0.0001). In the more demanding OppAlt programs that require behavioural flexibility (Endo et al., 2011), already during the first night, mice performed below the 75% chance level and in both programs (OppAlt 63.4±2.8% and Rev 61.3±3.6%), and they demonstrated significant learning ability (OppAlt *F* = 9.9, *p* < 0.0001, Rev *F* = 9.6, *p* < 0.0001). In the first OppAlt mice had their lowest average error percentage on Day 3 (40.2±5.6%, vs. Day 1 *p* = 0.030) and they remained on a similar level until the sixth day (41.6±2.8%, vs. Day 1 *p* < 0.0001). In the second OppAlt Rev, the mice behaved similarly, with the best performance on average on Day 3 (45.5±3.5%, vs. Day 1 *p* = 0.014) and thereafter maintaining the level until the last day (45.9±4.3%, vs. Day 1 *p* = 0.080). In the fifth and most demanding program Patrolling (Patrol2 in **Fig. 1A**), the mice did not show a clear learning tendency (*F* = 2.0, *p* = 0.096). Still, on average the mice performed below the chance level of 75% throughout the training period (Day 1 59.7±2.5%, Day 6 55.2±3.7%). Although the program is similar to Patrolling, in the Chaining program mice exhibited a clearer learning curve (*F* = 5.5, *p* = 0.0004). Again, on the first day, the error rate was 64.1±2.3% and it was lowest on the fourth day (49.7±4.0%, vs. Day 1 *p* = 0.026) after which no significant improvement occurred (Day 6 51.1±3.1%, vs. Day 1 *p* = 0.100).

**Figure 3.**
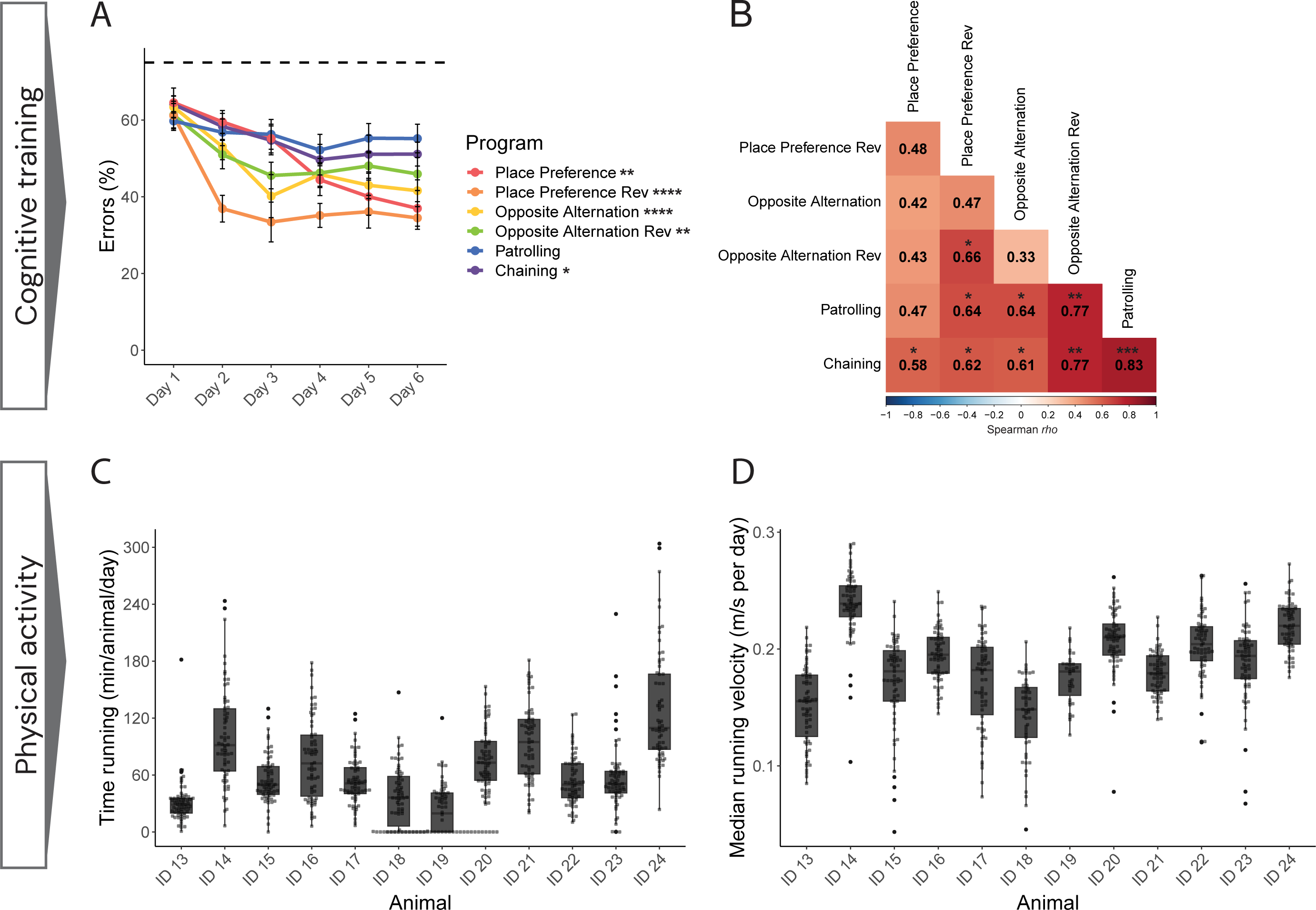
The lifestyle intervention domains in the PAW group. (**A**) All mice in the PAW group successfully underwent the cognitive training domain of the intervention period. Learning curves for the six different cognitive training programs are plotted for the six consecutive days that each program was running. Asterisks next to program names indicate the difference observed between the training day 1 and day 6 (day 4 for Opposite Alternation Rev and Chaining) as assessed in the post hoc test. The dashed line represents the 75% chance level. (**B**) Correlations between the performances in the different cognitive training programs as measured by learning curve AUCs are annotated by Spearman *rho*. Individual physical activity was measured as the (**C**) time in minutes each animal ran per day and the (**D**) velocity in meters per second they ran each day. ID 18 did not run for 14 days and ID 19 did not for 31 days (0 min/animal/day in **C**); These days are not considered in (**D**). (**A**) Data presented as mean±SEM. (**C, D**) Data presented as median (thick black line), 25^th^ and 75^th^ percentile (box) and 1.5 x interquartile range (whiskers). * *p* < 0.05, ** *p* < 0.01, *** *p* < 0.001, **** *p* < 0.0001

Despite some individual variability in the learning curves, the AUCs representing the overall performance of individual animals in the different programs correlated positively with each other (**Fig. 3B**). Specifically, we observed a strong correlation between the performance in the two similar programs, Patrolling and Chaining (Spearman’s rank correlation, *rho* = 0.83, unadjusted *p* = 0.001). Additionally, the performance in these two programs strongly correlated with the performance in OppAlt Rev (*rho* = 0.77, *p* = 0.003 for both Chaining and Patrolling). Performance in Chaining was furthermore significantly correlated with the performance in all the other programs (PP *rho* = 0.58, *p* = 0.048, PP Rev *rho* = 0.68, *p* = 0.033, OppAlt *rho* = 0.61, *p* = 0.036). Performance in Patrolling significantly correlated also with the performance in PP Rev and OppAlt (*rho* = 0.64, *p* = 0.024 for both). An additional significant correlation was observed between PP Rev and OppAlt Rev performances (*rho* = 0.66, *p* = 0.018).

In addition to cognitive training, the PAW group received physical training through free access to two running wheels. Over the intervention period, significant differences were detected between individual mice in running activity, which was calculated as the number of minutes spent running each day (Kruskal Wallis rank sum test, *H*_(11)_ = 338.2, *p* < 0.0001; **Fig. 3C**, **Supplementary Table 2**). Some mice did not run at all or very little on certain days (e.g., ID 13, 18, 19, and 23), while some other animals ran for hours (e.g., ID 14 and 24). The differences in running activity were reflected also in the distances ran (*H*_(11)_ = 412.4, *p* < 0.0001; **Supplementary Fig. 2A**). Moreover, the running speed for the individual mice varied substantially (*H*_(11)_ = 350.8, *p* < 0.0001; **Fig. 3D**, **Supplementary Table 3**). We further investigated whether there were any relationships between the performance in the cognitive training tasks and physical activity (**Supplementary Fig. 2B**). We observed a trend towards a negative correlation between the learning curve AUC in PP and median daily running time (Spearman’s rank correlation, *rho* = –0.55, *p* = 0.071).

### Evaluation of locomotion, anxiety-like behaviour and cognitive functions upon interventions

After completion of the intervention protocols, we first assessed whether the interventions impacted the locomotor activity of the mice by using the OF test. All three groups moved a similar distance over the 30-minute trial (one-way ANOVA, *F*_(2, 33)_ = 0.03, *p* = 0.970) with a similar speed (*F* = 1.50, *p* = 0.237; **Supplementary Fig. 3A**, B). Neither were there differences in the frequency (*F* = 0.51, *p* = 0.608) or time spent rearing (*F* = 0.46, *p* = 0.636; **Supplementary Fig. 3C**, D). On average, all three groups also spent similar amounts of time in the centre (*F* = 1.26, *p* = 0.298; **Supplementary Fig. 3E**). When analysing the trial period in 10-minute intervals, we observed significant differences within the groups. For the distance moved, there was a significant time effect (two-way mixed model ANOVA, *F*_(2, 66)_ = 45.53, *p* < 0.0001) and both Control and PAW mice moved less during the 10–20 min and 20–30 min intervals compared to the 0–10 min interval (post hoc paired Student’s *t*-test with Bonferroni correction, Control *p* = 0.003 and *p* = 0.002, PAW *p* = 0.0005 and *p* < 0.0001, respectively; **Fig. 4A**), showing the expected progressive habituation of the animals to the new environment over time. On the other hand, this effect was not observed in the Pharma group, where no statistically significant differences between the intervals were found. The average speed remained similar for all groups throughout the intervals (**Supplementary Fig. 3F**), while the percentage of time spent in the arena centre increased significantly only for the Pharma group (*F* = 10.46, *p* = 0.0001, time effect) between the first and the last 10 minutes of the trial (*p* = 0.005; **Fig. 4B**). Mice in all three groups increased the time spent rearing from the first to last interval (*F* = 26.64, *p* < 0.0001, time effect; Control *p* = 0.036, PAW *p* = 0.045, Pharma *p* = 0.009; **Fig. 4C**). Still, the number of rearings (*F* = 29.34, *p* < 0.0001, time effect) was only increased for the Pharma group from first to last interval (*p* = 0.002; **Fig. 4D**). Representative OF activity patterns of the 30-min trial for each group are presented in **Figure 4E**.

**Figure 4.**
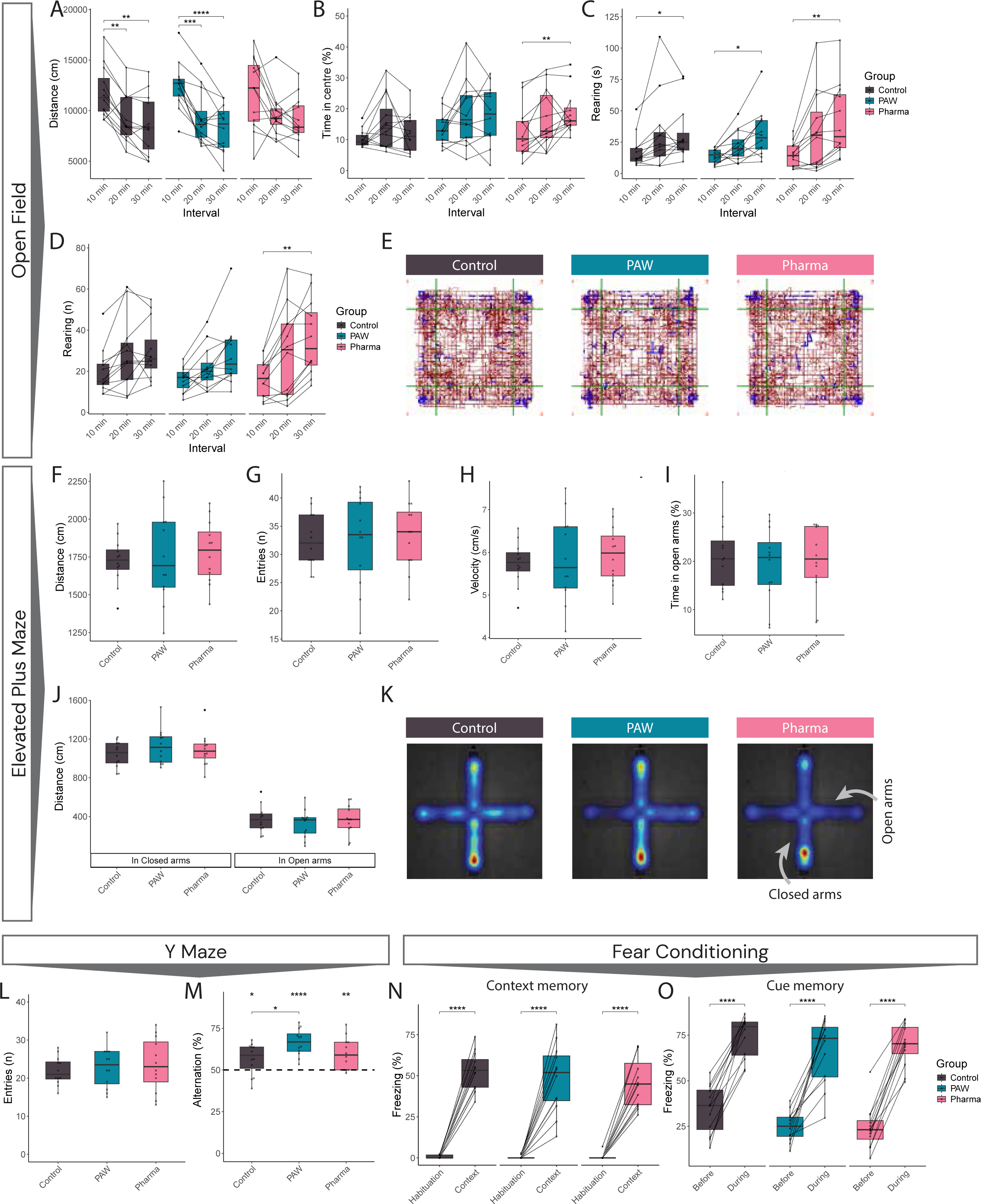
Behavioural outcomes after the intervention period. (**A–E**) Open Field test results; Changes in (**A**) total distance moved, (**B**) percentage of time spent in the centre of the arena, (**C**) duration in seconds spent rearing, and (**D**) number of rearings performed throughout the ten-minute intervals constituting the full 30 min trial. (**E**) Representative images of the total distance moved during the trials by one animal in each group. (**F–K**) Elevated Plus Maze test results; (**F**) Total distance moved, (**G**) total number of entries performed, (**H**) velocity, and (**I**) percentage of time spent in the open arms during the five-minute trial. (**J**) Distance moved in closed and open arms, respectively. (**K**) Representative heatmaps of the activity during the trials by one animal in each group. Performance in the Y Maze test was assessed by (**L**) the total number of entries performed during the trials and (**M**) the percentage of alternation. The dashed line represents the 50% chance level in (**M**). In the Fear Conditioning test, the percentage of freezing was measured for (**N**) context memory during habituation on the first day and the following day in the same context and for (**O**) cue memory on the last day in the new context before the sound cue and during the sound cue. Data presented as median (thick black line), 25^th^ and 75^th^ percentile (box) and 1.5 x interquartile range (whiskers). * *p* < 0.05, ** *p* < 0.01, *** *p* < 0.001, **** *p* < 0.0001

Next, anxiety-like behaviour was evaluated in the EPM test. Overall, all three groups moved on average a similar distance (one-way ANOVA, *F*_(2, 33)_ = 0.21, *p* = 0.813), did a similar number of entries (*F* = 0.09, *p* = 0.917) and showed similar speed (*F* = 0.21, *p* = 0.814; **Fig. 4F–H**). Moreover, there were no differences in the time spent in the open arms (*F* = 0.23, *p* = 0.797; **Fig. 4I**) nor distance moved neither in closed or open arms (*F* = 0.73, *p* = 0.492 and *F* = 0.43, *p* = 0.656, respectively; **Fig. 4J**). Representative EPM activity patterns for each group are presented in **Figure 4K**.

The exploration and short-term spatial working memory were tested using the Y maze. All groups performed on average a similar number of entries into the three arms (*F* = 0.25, *p* = 0.782; **Fig. 4L**). All groups also demonstrated good cognitive performance by performing a percentage of alternation above the chance level of 50% (unadjusted *p*-values: Control 56.7±2.8%, *p* = 0.033; PAW 66.5±2.3%, *p* < 0.0001; Pharma 59.8±2.8%, *p* = 0.005). Still, the alternation percentage differed between the groups (*F* = 3.72, *p* = 0.035; **Fig. 4M**), with the PAW (but not Pharma) group outperforming the Control group, displaying a significantly better alternation percentage (*p* = 0.036). Lastly, mice underwent the FCond test. All mice learned and remembered the association of the context with the electric shock stimulus, as shown by a significant increase in the freezing time after 24 h (two-way mixed models ANOVA, *F*_(1, 33)_ = 331.66, *p* < 0.0001, for the one-way time interaction) but we did not observe group differences (*F* = 0.525, *p* = 0.596, for the one-way group interaction; **Fig. 4N**). After two days, all mice also remembered the cue-shock association (*F* = 279.48, *p* < 0.0001, for the time interaction), again with no group differences (*F* = 2.41, *p* = 0.106, for the group interaction; **Fig. 4O**).

Additionally, mice underwent the NOR test to assess recognition memory. During the second day, there were no group differences in the total time spent exploring the objects (*F* = 0.71, *p* = 0.501; **Supplementary Fig. 3G**). However, when analysing the object exploration time by object type (flask or egg), we observed a preference toward the flask object in all three groups (two-way ANOVA, *F*_(1, 30)_ = 25.75, *p* < 0.0001, object effect), which was significant for the PAW group (*p* = 0.043; **Supplementary Fig. 3H**). The discrimination index between familiar and novel objects measured on the third day did not show group differences (*F* = 0.05, *p* = 0.953; **Supplementary Fig. 3I**). Still, we found a significant object preference based on the object type (*F* = 308.68, *p* < 0.0001), as indicated by the discrimination index which was significantly higher for the flask than egg as novel object (*p* < 0.0001 for all groups; **Supplementary Fig. 3J**), independently from the group.

Finally, we explored whether the performance of the PAW group in the Y maze and FCond tests was associated with their running activity or learning during the intervention. In the case of the Y maze test, there was a trend of median daily running time being positively correlated to alternation percentage (Spearman’s *rho* = 0.50, *p* = 0.099; **Supplementary Fig. 4A**). For the FCond tests we did not observe significant correlations (**Supplementary Fig. 4B**, C). Alternation percentage in the Y maze test was not significantly correlated with any of the cognitive training programs but there was a trend towards a correlation with PP Rev (*rho* = 0.55, *p* = 0.071; **Supplementary Fig. 4D**). No significant correlations were observed between FCond and cognitive training performances either, except between the change in freezing percentage in the cue test and the AUCs for Chaining learning curve (*rho* = 0.61, *p* = 0.040; **Supplementary Fig. 4E**, F).

### Intervention-induced effects on hippocampal proteomics

To investigate how the interventions impacted the hippocampi of the mice, proteomics and subsequent pathways analyses were performed. In the comparison of Control to PAW mice, we detected 4404 proteins out of which 241 were upregulated and 55 were downregulated in the PAW hippocampi (absolute Fold Change ≥ 1.25, *p* < 0.05 with FDR 5%; **Fig. 5A**, **Supplementary Table 4**). The 20 proteins with the largest difference compared to Controls are listed in **Table 2**. In the comparison of Control and Pharma mice, 4418 proteins were detected yet, the levels were not significantly different for any of the proteins in the Pharma mice (**Fig. 5B**, proteins with unadjusted *p*-value < 0.05 are presented in **Supplementary Table 5**). The two comparisons had 4403 proteins in common, 1 protein was detected in the Control vs. PAW comparison only, and 15 proteins were detected in the Control vs. Pharma comparison only.

**Figure 5.**
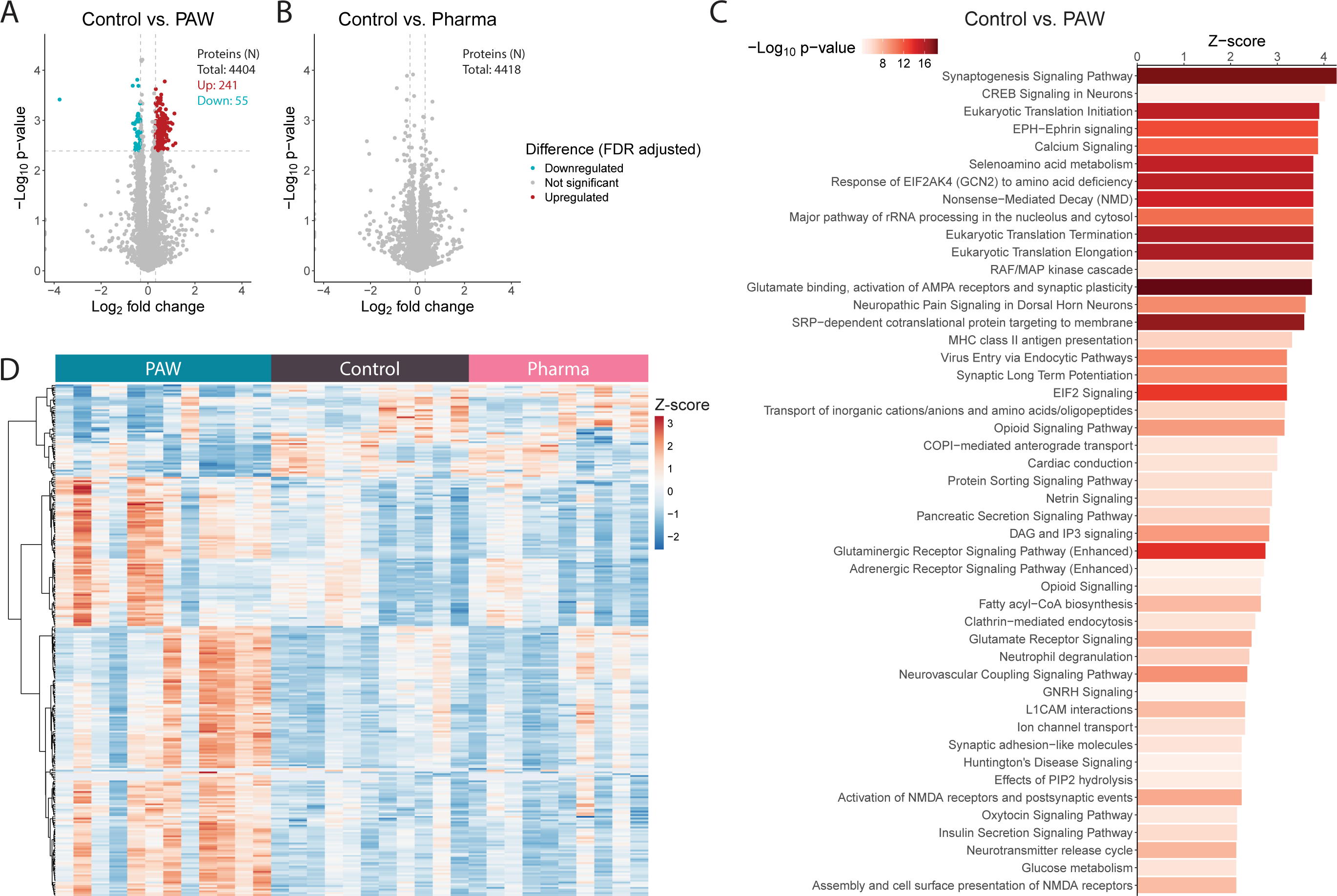
Differences in hippocampal proteomics upon intervention. (**A, B**) Volcano plots representing the differently expressed protein levels when comparing (**A**) PAW to Control and (**B**) Pharma to Control. The dashed lines indicate the limits for significantly altered protein levels (|fold change| > 1.25, FDR adjusted *p*-value < 0.05). (**C**) Significantly altered pathways in the PAW hippocampus compared to the Control were identified in the Ingenuity Pathway Analysis (IPA) canonical pathway analysis. Analysis was based on the significantly different proteins found in (**A**); IPA identified 288 out of the total 296 differently expressed proteins. There were 47 pathways with an absolute Z-score of more than 2 and a *p*-value < 0.0001. (**D**) Heatmap showing the relative protein levels of individual mice for the 288 significantly different proteins between PAW and Control that were included in pathway analysis. Proteins were clustered using hierarchical clustering with dissimilarity measured by Euclidean distance and by the agglomeration method Ward D2. Mice were not clustered.

**Table 2.**
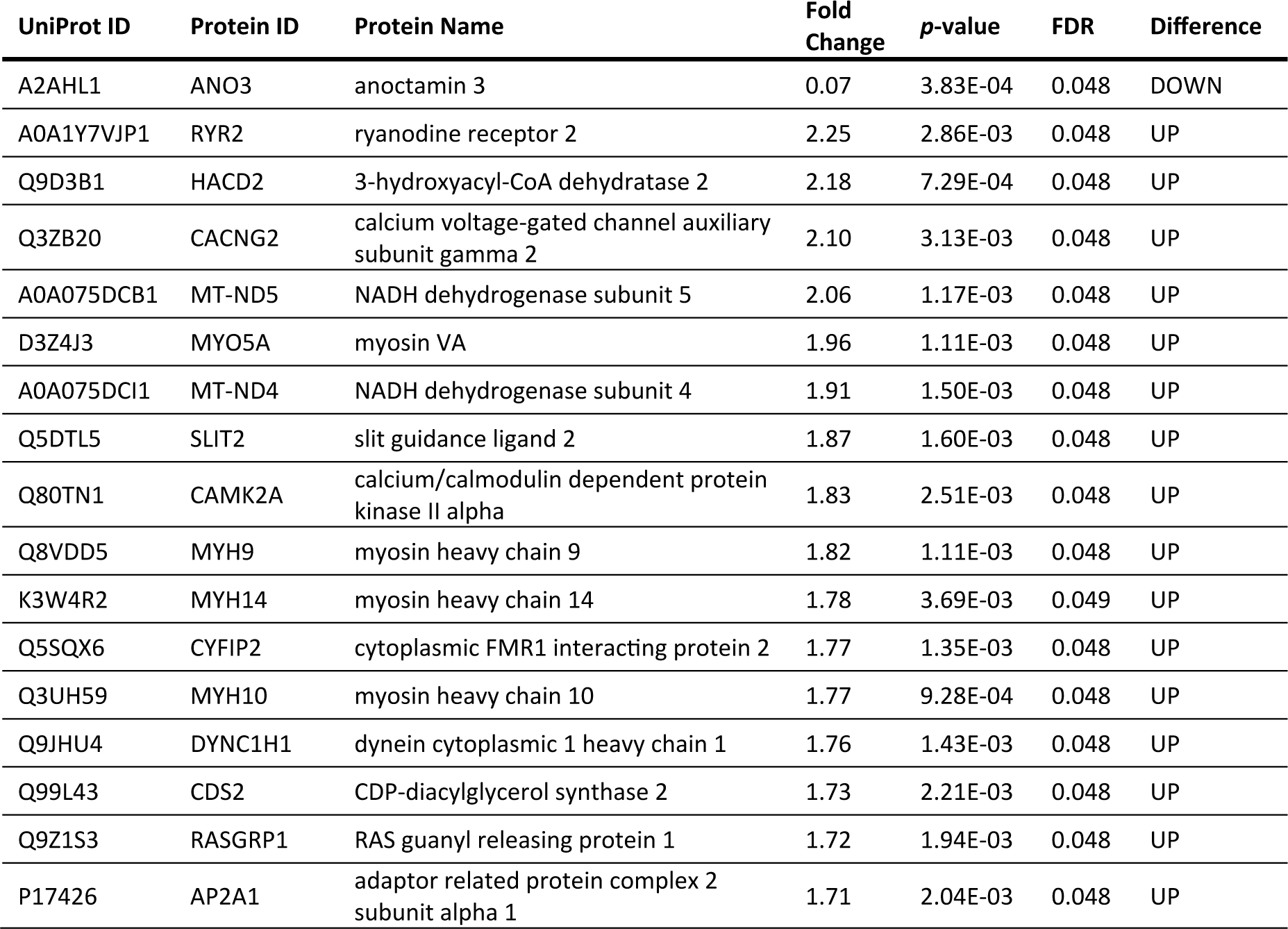
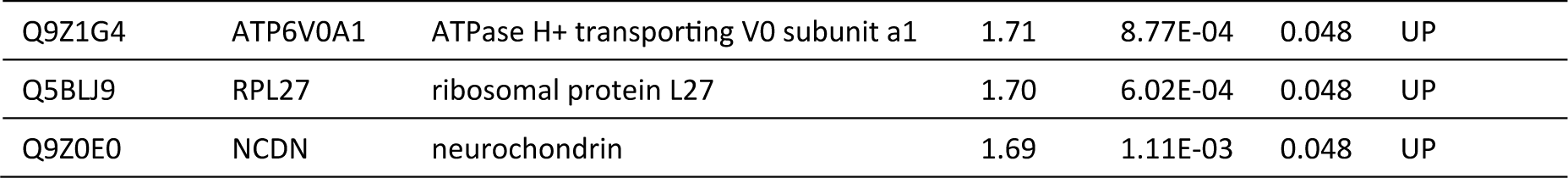
List of 20 differentially expressed proteins in the PAW group compared to Control, ordered by the magnitude of fold change.

In the PAW group, the significantly differently expressed proteins were associated with 47 pathways (|Z-score| ≥ 2, *p* < 0.0001; **Fig. 5C**, the full list is presented in **Supplementary Table 6**). The most strongly influenced pathways were related to the upregulation of the synaptogenesis signalling pathway (Z = 4.27, *p* = 1.26E^–18^) and glutamate binding, activation of AMPA receptors and synaptic plasticity (Z = 3.74, *p* = 3.16E^–19^). Other upregulated pathways that are linked to synaptic and neuronal plasticity and function were, for example, CREB signalling in neurons, EPH-Ephrin signalling, calcium signalling, and synaptic long-term potentiation. The only significantly downregulated pathway was related to RhoGDI signalling. The differences in relative protein levels between individual mice for the significantly different proteins are presented in **Figure 5D**.

## Discussion

With the increasing evidence of the potential of dementia prevention with multimodal lifestyle interventions, the need to understand the working mechanisms behind the beneficial cognitive effects becomes inevitable. Here, we propose a new model to study a multimodal lifestyle intervention in mice. To our knowledge, our study is the first to combine physical activity, cognitive training, and dietary intervention to enhance or maintain cognitive performance in mice. Our setup provided a platform to not only study the intervention effects but also the performance and adherence to the intervention itself. Importantly, the intervention was fully run in the home cage without the requirement of social isolation – a factor known the be harmful to mice (Benfato et al., 2022; Wang et al., 2018).

Mice included in the multimodal lifestyle intervention were active in all the different lifestyle domains. We observed that the PAW group on average performed below the chance level in all six different cognitive training programs, indicating good learning abilities throughout the intervention. Moreover, the overall performances of individual mice as measured by the AUC of the learning curves in the different programs positively correlated with each other, suggesting mice tended to perform similarly independent of the program. The difficulty level of the programs was increased throughout the intervention, with PP programs being the easiest requiring simple place learning, and Patrolling and Chaining the most difficult demanding good spatial working memory (Kiryk et al., 2011; Voikar et al., 2018). This increment in difficulty was also reflected in the learning curves; on a group level, the mice did not significantly reduce the error rate from day one to day six in Patrolling and Chaining. However, the flatter learning curves may be a result of implementing only weak positive reinforcement to learn the programs. Previous IC studies have used negative or strong positive reinforcement either by the restriction of water access to specific times or by the introduction of sweetened water, respectively (Kiryk et al., 2011, 2020; Voikar et al., 2018). Still, learning without temporal water restriction in, for instance, PP, Patrolling, and Chaining tasks, has been reported (Bramati et al., 2023; Voikar et al., 2018). Although rodents are capable of adaptation, water restriction may cause stress and other physiological changes in animals (Rowland, 2007; Vasilev et al., 2021). Water restriction could also impact wheel running activity because physical activity increases water consumption in mice (Goodrick, 1978). Since the PAW intervention aimed to improve the lifestyle, we wanted to avoid unnecessary stress interfering with wellbeing and behaviour. To that end, we decided (1) not to introduce temporal water restriction, (2) we let mice learn and adjust to each program for six consecutive nights, and (3) we included rest days between each program to let mice recover. Other stress-inducing elements in cognitive training studies are human handling, removal from home cages and cage mates, and possibly aversive test arenas (Alanko et al., 2022). By introducing cognitive training in a home cage setting we were able to avoid these elements. The body weight development, similar water consumption, and similar behaviour in the OF and EPM tests between Control and PAW mice suggest that the PAW mice did not experience stress due to the cognitive training.

Although all mice in the PAW group actively used the RW, there was some individual variance. The variation observed may be attributed to differences in motivation and perceived reward (Novak et al., 2012). As with cognitive training, we did not want to force physical activity and endurance training by using, for instance, treadmill running or swimming which could be stressful for the mice. By using the AGs, mice could use the RWs undisturbed without the need for individual housing. While individually housed, 6-month-old C57BL/6 females run approximately 2 km/day (Venezia et al., 2016), mice in the current study ran on average between 0.1 and 1.4 km/day. Considering that 12 mice shared two running wheels in our study the running distance could be expected to be lower than in previous studies. The detailed and individual data collection allowed for the analysis of associations between physical activity and cognitive training. We observed that the running activity was not related to the performance in any cognitive training programs. A recent study, on the other hand, showed that mice with high physical activity made more errors in PP than less active mice (Niiranen et al., 2023). However, we observed positive, yet non-significant, correlations between wheel running activity and spatial working memory.

Previous studies in mice aiming for enhancing or maintaining cognitive performance have mainly focused on single-domain interventions or a combination of two domains, the most common combination being exercise and cognitive enrichment, mainly in the form of environmental enrichment with running wheels (EE-RW). One study has combined EE-RW with a dietary intervention (Esselun et al., 2021). Even though findings from single-domain interventions have largely supported the beneficial effect of lifestyle interventions in mouse models of cognitive decline, it is important to understand the synergistic effects of different domains (Alanko et al., 2022). Our finding of improved short-term working memory as a result of a multimodal lifestyle intervention is largely in line with previous reports from EE-RW interventions. Some EE-RW intervention studies have resulted in improved spatial working memory in wild type mouse strains (Bramati et al., 2023; Hüttenrauch et al., 2016; Rabadán et al., 2019; Schmidt et al., 2022; Stuart et al., 2017), while some studies have lacked the effect (Stazi et al., 2023; Wang et al., 2018; Wu et al., 2022). In contrast to our findings, a previous study reported worse short-term spatial working memory in female wild type mice exposed to EE-RW for nine months compared to standard-housed mice (Sansevero et al., 2016). It has been proposed that, in C57BL/6, the effects on neurogenesis and spatial learning are influenced by physical activity, with little or no effect from the EE component (Kobilo et al., 2011; Lambert et al., 2005; Mustroph et al., 2012). Considering that reported EE-RW interventions greatly vary concerning intervention length, the exact environmental configuration, and the age of the tested mice, variation in outcomes can be expected.

A combination of dietary intervention together with EE-RW has previously been associated with improved cognitive performance in a mouse model. In this study with female NMRI mice, however, the PUFA-enriched walnut diet alone restored age-induced decline in cognitive performance to the same extent as the diet and EE-RW combination (Esselun et al., 2021). We selected to add Fortasyn diet to our intervention because of its known brain health-enhancing properties (Jansen et al., 2013; Wiesmann et al., 2016). Fortasyn has been found to have clinical benefits in patients with mild cognitive impairment due to Alzheimer’s disease alone as well as in combination with the FINGER lifestyle intervention, yet the results from preclinical models have been modest (Jansen et al., 2014; Koivisto et al., 2014; Soininen et al., 2021; Thunborg et al., 2024; Wiesmann et al., 2013). We hypothesise that the molecular effects that Fortasyn has on the mouse brain may benefit from the synergistic impact of combining Fortasyn diet with physical activity and cognitive training and therefore that the addition of those domains was responsible for the stronger cognitive performance in the current study.

To investigate the influence of improved vascular health achieved by medication rather than by lifestyle, a group of mice received a lipid-lowering and an antihypertensive drug dosed in the diet. These drug types have been shown to alter brain function in mice (Guo et al., 2021; Yamada et al., 2010; Yoo et al., 2021). Simvastatin, which is a lipophilic statin like atorvastatin, has been found to cause impairment in synaptic plasticity, leading to poor cognitive performance in healthy C57BL/6J males (Guo et al., 2021). Conversely, a combination therapy with atorvastatin and captopril, which is a centrally active antihypertensive drug unlike enalapril, induced neurogenesis in C57BL/6J males without effects on spatial memory (Yoo et al., 2021). In a mouse model of Alzheimer’s disease, enalapril did not ameliorate cognitive deficits (Yamada et al., 2010). Vascular risk monitoring is an essential part of the FINGER model as previous research has demonstrated the importance of cardiovascular diseases as risk factors for dementia (Hughes et al., 2020; Solomon et al., 2009). However, controlling for individual vascular risk factors alone can be insufficient in reducing the dementia risk (Mcguinness et al., 2016; Williamson et al., 2019). In line with this notion, we did not observe effects on cognition upon the combination treatment with atorvastatin and enalapril regardless of the effects on vascular health.

Since the Pharma group was receiving an antihypertensive drug in the diet, a reduction in BP was expected. Intriguingly, the PAW intervention had a similar BP-lowering effect. Recently, exercise has been demonstrated to lower high-fat-diet-induced elevation in BP, even to a similar extent as enalapril (Lino Rodrigues et al., 2022; Salles et al., 2024). In contrast, exercise did not reduce BP in mouse models of atherosclerosis or hypertension (Bochi et al., 2022; Chang et al., 2022). PUFA-enriched diets, on the other hand, have had little or no effect on BP in mouse models of metabolic and cardiovascular disorders (Gladine et al., 2014; Redondo Useros et al., 2019; Takashima et al., 2016). Compared to the Control and PAW groups that gained weight during the study, mice undergoing the pharmacological intervention kept the same body weight throughout the study and on average ate less than the two other groups. Despite lower food consumption, Pharma mice performed significantly more licks during the intervention compared to the other groups. Similar antihypertensive-mediated effects on water intake have previously been observed in rats (Rowland & Fregly, 1988).

Although resulting in cognitive benefits, EE-RW has in some cases been shown to increase anxiety-like behaviour in wild type mouse strains. Both NMRI and C57BL/6 mice in EE-RW groups have been observed to spend less time in open arms in the EPM test compared to their sedentary controls (Pietropaolo et al., 2014; Rabadán et al., 2019; Zhu et al., 2006). Such behaviour was not seen for the PAW mice. Furthermore, EE-RW and RW interventions have been shown to reduce locomotor activity in the OF test (Kobilo et al., 2011; Zhu et al., 2006), yet this effect was not observed in the current study as the PAW group had a similar activity pattern to the Control group where the distance travelled declined from the first to the last test interval. In contrast, the Pharma group did not have a significant decline in the activity from the first to the last interval in the test. Moreover, during the last 10-minute interval the Pharma mice performed more rearings and spent more time in the centre of the arena than during the first 10-minute interval, altogether implying enhanced explorative behaviour compared to the other groups.

To unravel in a non-hypothesis-driven way which mechanisms were associated with the hippocampal-dependent effect on spatial memory, we analysed the hippocampal proteome of the mice. In the PAW group, we found several upregulated pathways related to synaptic functioning and plasticity. Our findings are in line with previous research that has shown enhanced synaptic plasticity upon EE-RW interventions (Hüttenrauch et al., 2016; Schmidt et al., 2022; Stuart et al., 2017; Wang et al., 2018). Levels of synaptic proteins in the hippocampus have also been observed to increase as a result of single-domain interventions of EE, exercise, and Fortasyn diet (Lambert et al., 2005; Wiesmann et al., 2016). Previous EE-RW studies have additionally shown increased numbers of neurons or region volumes in the hippocampus (Bramati et al., 2023; Hüttenrauch et al., 2016). In comparison to PAW, there were no significantly altered pathways in the Pharma group. Previous research indicates that some statins and angiotensin-converting enzyme inhibitors – alone or in combination – may ameliorate deficits in long-term potentiation, reduce inflammatory and pathological markers, and improve cognition in Alzheimer’s disease mouse models (Collu et al., 2023; Yamada et al., 2010; Zhao et al., 2016), while such effects have not been observed in healthy mice (Guo et al., 2021; Yoo et al., 2021; Zhao et al., 2016). These findings imply that the beneficial effects these drug classes (considering the different properties of individual compounds) have on the brain depends on pathology.

This study has several limitations. First, since our study only included female mice, our results cannot be extrapolated to male mice. It is known that there are sex-dependent differences in both behavioural, physiological, and biological effects as a result of lifestyle-related interventions between female and male mice (Venezia et al., 2016; Zhu et al., 2006). Sex differences for both Alzheimer’s disease models and non-transgenic mice have previously been reported also for the PP and PP Rev performances in the IC (Mifflin et al., 2021). Second, the exercise intervention was based on voluntary wheel running, which may not increase physical fitness alike forced training, such as treadmill training (Bódis et al., 2024). The physiological and biological effects may be divergent depending on the type of exercise intervention (Bódis et al., 2024). Still, high-resistance physical training is not necessarily required to observe beneficial effects (Han et al., 2023; Leuchtmann et al., 2023). Another limitation related to voluntary wheel running is the individual variability which we also observed in our study. Third, all three groups of mice were exposed to similar enhanced social interaction during the study. Social activities are one of the key elements in the FINGER model (Kivipelto et al., 2013) and could have been incorporated into the current study design by having Control and Pharma mice living in smaller groups. However, to keep the basic conditions between groups as similar as possible and to collect activity data during the intervention, we decided to use the same home cage environment for all groups which resulted in large mouse groups also for Pharma and Control.

To conclude, in adult wild type mice without cognitive impairment -causing brain pathologies, a multimodal lifestyle intervention enhanced spatial working memory, which was associated with the activation of pathways related to synaptic plasticity in the hippocampus. Although the pharmacological intervention was effective in mediating vascular health, it was insufficient to alter cognition. The lifestyle intervention had both cardiovascular and brain health -promoting effects that were most likely independent of each other. While our findings indicate that a multimodal lifestyle intervention can positively influence molecular pathways in the brain that improve or maintain cognitive function, it is to be investigated whether the effect is similar when the brain is affected by ongoing pathologies. Our findings endorse future animal studies that combine several lifestyle domains to further decipher molecular mechanisms related to memory gains, particularly in animal models relevant to dementia research.

## Acknowledgements

The authors would like to thank: Dr. Vootele Vöikar and the Mouse Behavioural Phenotyping Facility (Helsinki Institute of Life Science) for training and help with the IntelliCage method; Dr. Qian Yu and the Animal Behavioural Core Facility (ABCF) at Karolinska Institutet for providing the behavioural equipment; Beatrice Scarpa for assisting in animal care take and behavioural testing; Anna Lauer for assistance in tissue dissection. VA, AS, TH and AM are members of the EU Joint Programs – Neurodegenerative Disease Research Multi-MeMo project and funding was in part provided by the BMBF 03ED2306 grant and Research Council of Finland grant 357810.

The study was further funded by: Alzheimerfonden (Sweden); Region Stockholm (ALF grant); European Research Council (ERC, 804371); EU Joint Programme—Neurodegenerative Disease Research (JPND) EURO-FINGERS and Multi-MeMo grants; Swedish Research Council; Center for Innovative Medicine (CIMED) at Region Stockholm (Sweden); Stiftelsen Stockholms Sjukhem (Sweden); Hjärnfonden (Sweden); Margaretha af Ugglas Foundation (Sweden); King Gustav V:s and Queen Victorias Foundation (Sweden); Gun och Bertil Stohnes Stiftelse (Sweden); Stiftelsen för Gamla Tjänarinnor (Sweden).

---

**Supplementary Figure 1. Activity patterns in the IntelliCage.** (**A**) Total number of visits to the IntelliCage (IC) corners during the first hour when mice were placed in the cages. Mice were placed into their respective cages at 7 PM at the beginning of the dark phase. (**B**) The familiar cage activity was measured as corner visits for each hour during the following 24 hours after the initial hour after moving into the ICs. The grey box indicates the dark phase. (**C**) Daytime activity was measured as the average number of corner visits per animal during day and night both before and during the intervention. (**D**) The average duration of a corner visit per animal before and during the intervention. (**E**) General activity patterns were measured as corner visits throughout the study period. The study period is colour-annotated according to the ongoing program. Except for the Free Access program, the programs applied only to the PAW groups. Control and Pharma groups were on the Free Access program throughout the study. The drop in activity during the second week is an artefact caused by software issues. (**A, C, D**) Data presented as median (thick black line), 25^th^ and 75^th^ percentile (box) and 1.5 x interquartile range (whiskers). (**B, E**) Data presented as mean±SEM. * *p* < 0.05, ** *p* < 0.01, *** *p* < 0.001, **** *p* < 0.0001

**Supplementary Figure 2. Running distance and correlation between cognitive training performance and physical activity.** (**A**) Running distance in meters for each animal on each day during the intervention. ID 18 did not run for 14 days and ID 19 did not for 31 days (0 m/animal/day). (**B**) The individual plots represent the correlation between the AUC of the learning curves in the different cognitive training programs (y-axis) and the median running time in minutes for each animal (x-axis). Each plot is annotated with the Spearman *rho* and unadjusted *p*-value. Plotted trendlines were acquired by robust linear regressions.

**Supplementary Figure 3. Behavioural outcomes after the intervention period.** (**A–F**) Open Field test results; Total (**A**) distance moved, (**B**) velocity, (**C**) number of rearings performed, (**D**) duration in seconds spent rearing, and (**E**) percentage of time spent in the centre of the arena during the 30-minute trial. (**F**) Changes in velocity between the ten-minute intervals during the Open Field test. (**G– J**) Novel Object Recognition test results; (**G**) Total duration used for exploring the objects during the habituation day. The dashed line represents a minimum level (20 s) of exploration required to be considered for the novel object test. (**H**) Duration of time spent exploring the objects during the habituation day divided by the object type. (**I**) The discrimination index between the novel and the familiar object on the last test day. (**J**) The discrimination index divided based on the type of novel object. (**I, J**) The dashed line represents the level of spending equal time exploring both objects. (**A–J**) Data presented as median (thick black line), 25^th^ and 75^th^ percentile (box) and 1.5 x interquartile range (whiskers). * *p* < 0.05, **** *p* < 0.0001

**Supplementary Figure 4. Correlation between physical activity, cognitive training, and performance in cognitive tests.** (**A**) Correlation between the alternation percentage in the Y Maze test and the median time an animal ran per day. (**B**) Correlation between the freezing percentage during the context memory test day and (**C**) change in freezing percentage during the cue memory test day of Fear Conditioning and the median time an animal ran per day. (**D–F**) The individual plots represent the correlation between (**D**) the alternation percentage in the Y maze test, (**E**) the freezing percentage during the context memory test day, (**F**) the change in freezing percentage during the cue memory test day, and the AUC of the learning curves in the different cognitive training programs. (**A–F**) Plots are annotated with Spearman *rho* and *p*-value. Plotted trendlines were acquired by robust linear regressions.

