## Supplementary figures and images for "Mouse PAW: reverse-translating the FINGER multimodal lifestyle intervention enhances synaptic plasticity and cognition in adult wild type female mice"

### Supplementary figure 1

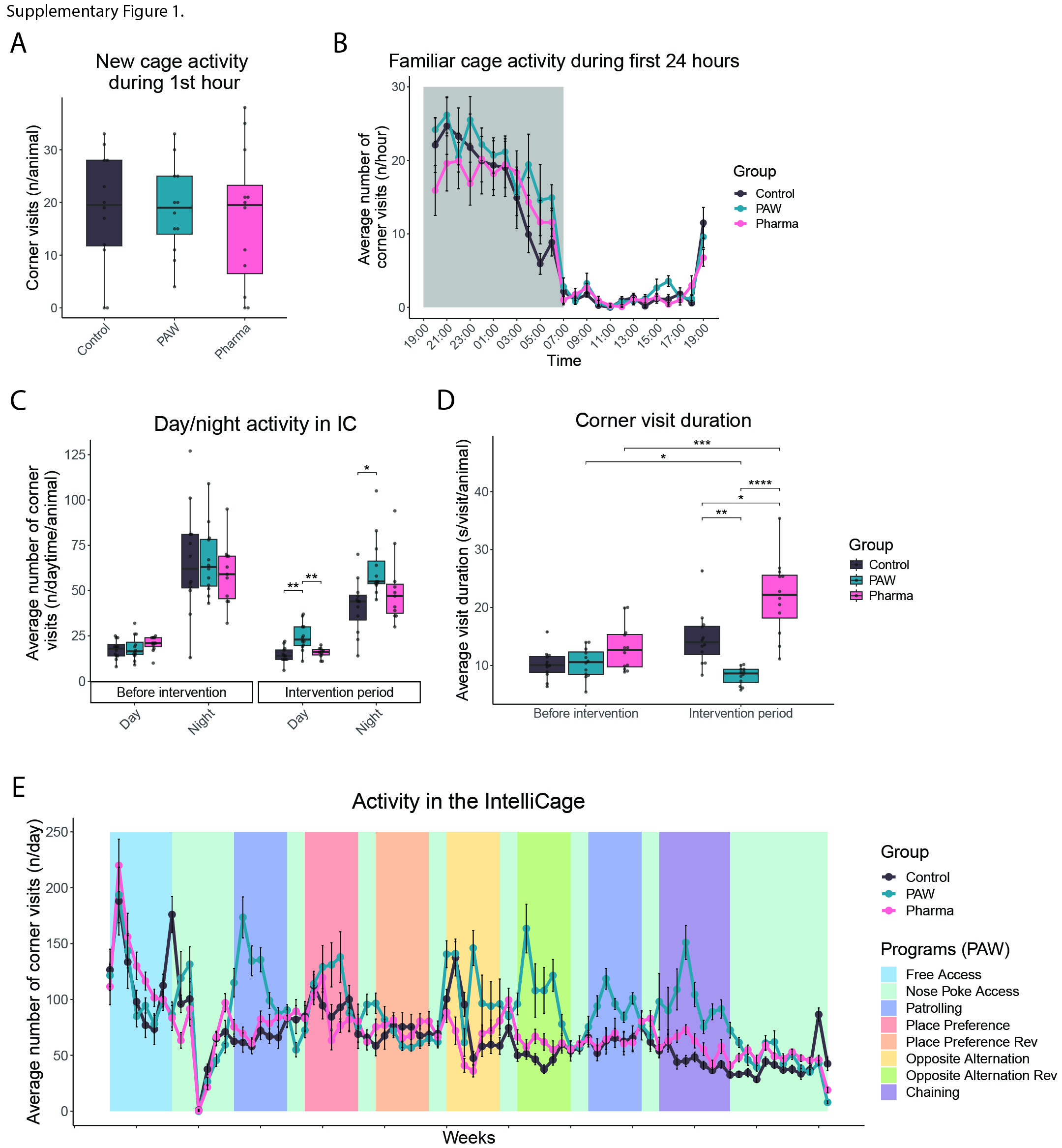

### Supplementary figure 2

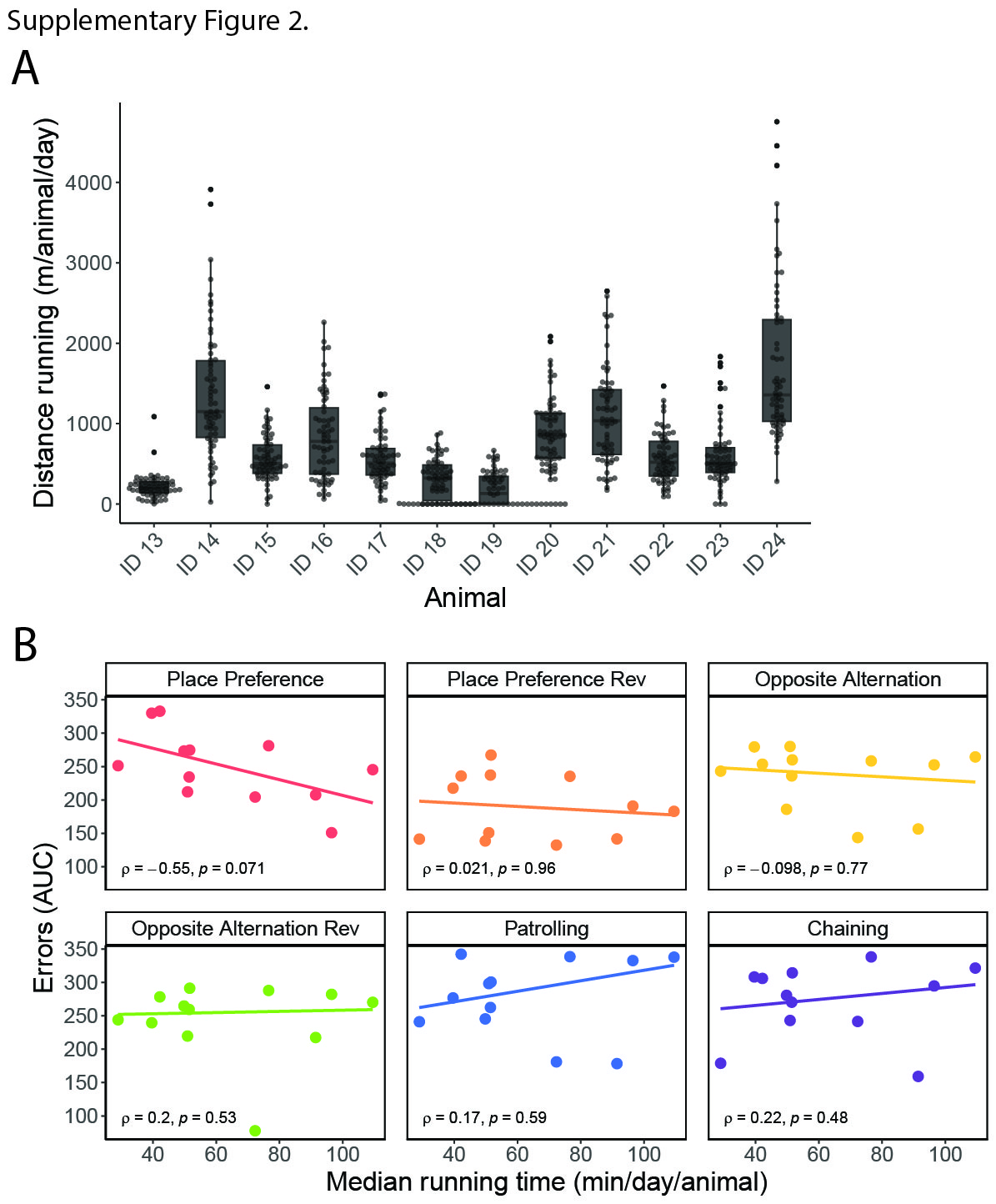

### Supplementary figure 3

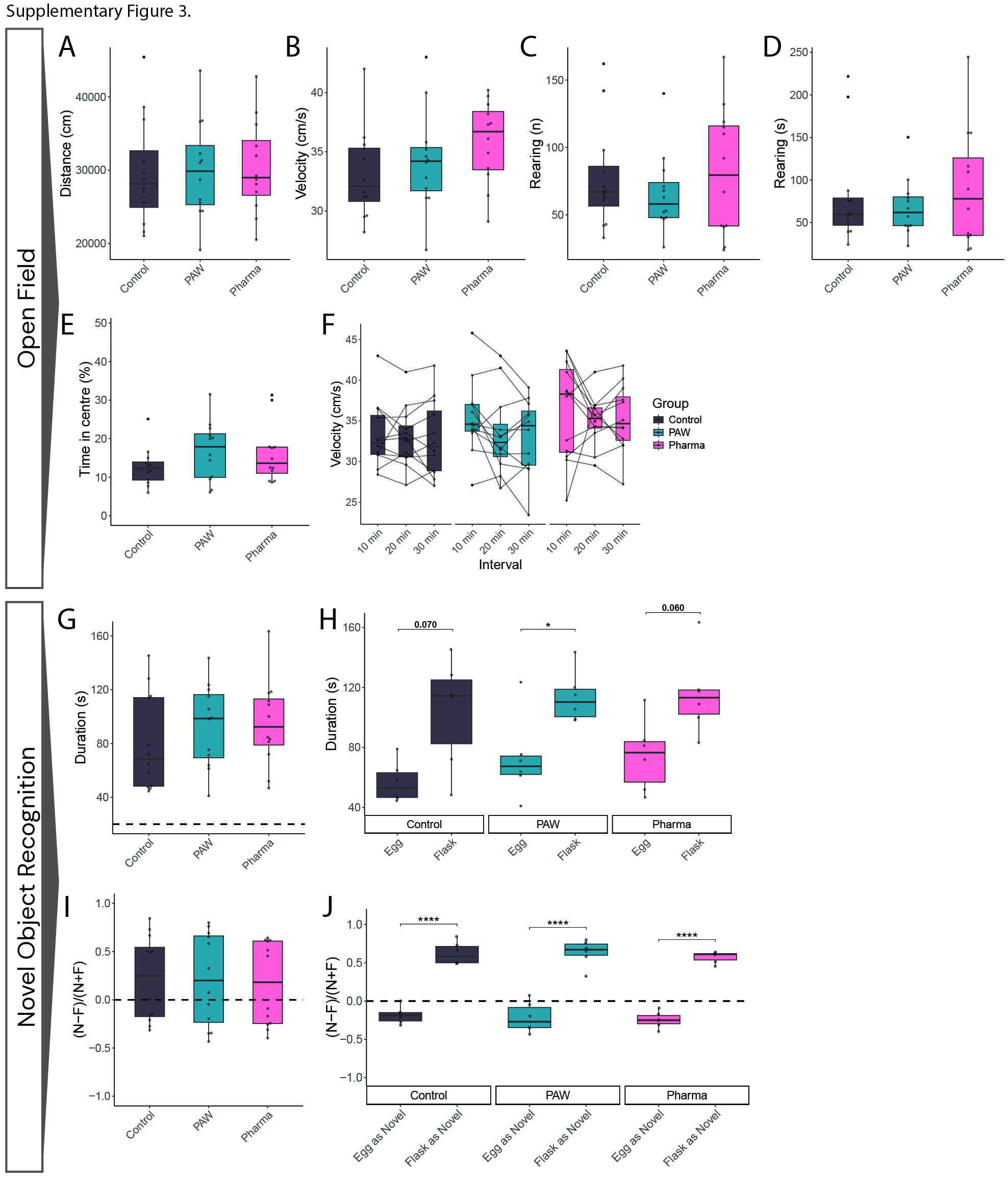

### Supplementary figure 4

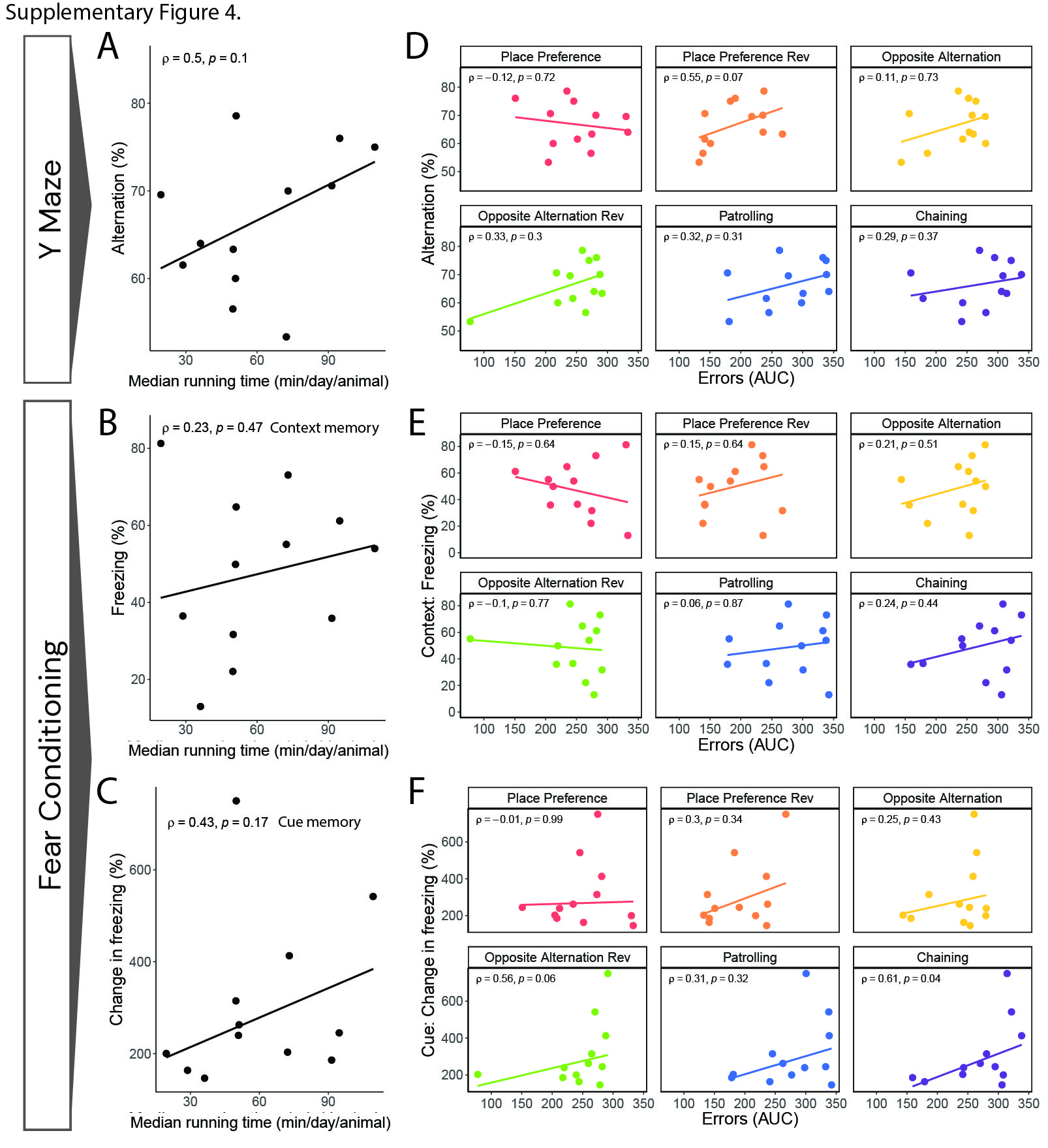
