## Supplementary tables 1-3 for "Mouse PAW: reverse-translating the FINGER multimodal lifestyle intervention enhances synaptic plasticity and cognition in adult wild type female mice"

**Supplementary Table 1. Diet specifications.**

| Ingredients | Control g / 100 g diet | Pharma g / 100 g diet | PAW g / 100 g diet |
|---|---|---|---|
| Cornstarch, pre-gelatinized | 35.59 | 35.57 | 32.51 |
| Caseine | 14.00 | 14.00 | 14.00 |
| Maltodextrin, 10 DE | 15.50 | 15.50 | 15.50 |
| Sucrose | 10.00 | 10.00 | 10.00 |
| Dextrose | 10.00 | 10.00 | 10.00 |
| Cellulose powder | 5.00 | 5.00 | 5.00 |
| Mineral & trace element premix (AIN-93M-MX) | 3.50 | 3.50 | 3.50 |
| Vitamin mix (AIN-93-VX) | 1.00 | 1.00 | 1.00 |
| **Fats:** | | | |
| Soy oil | 1.900 | 1.900 | |
| Coconut oil | 0.900 | 0.900 | 0.100 |
| Corn oil | 2.200 | 2.200 | 1.700 |
| Tuna fish oil | | | 3.200 |
| **Additions:** | | | |
| L-cystine | 0.180 | 0.180 | 0.180 |
| Choline chloride (50%, 0.434 g / g) | 0.230 | 0.230 | 0.922 |
| Tert-butylhydroquinone | 0.0008 | 0.0008 | 0.0008 |
| **Extra's:** | | | |
| Atorvastatin | | 0.01286 | |
| Enalapril | | 0.00857 | |
| Soy lecithin | | | 0.7547 |
| UMP disodium (24% $H_2O$) | | | 1.0000 |
| Ascorbic acid (100%) | | | 0.1600 |
| Vit. E (Tocopherol acetate, 50%) | | | 0.4650 |
| Vit. B6 (Pyridoxin HCl, 82%) | | | 0.0033 |
| Folic acid (100%) | | | 0.0006 |
| Vit. B12 (Cyanocobalamin 0,1%) | | | 0.0035 |
| Selenium (Na selenite • 5 $H_2O$) | | | 0.0003 |
| Total | 100.0 | 100.0 | 100.0 |
| Energy (kcal / 100 g chow) | 377.7 | 377.6 | 365.4 |
| Fat (g / 100 g chow) | 5.0 | 5.0 | 5.0 |

**Supplementary Table 2. Differences between the time running (min/day/animal) between the PAW mice.**

| Animal ID 1 | Animal ID 2 | n1 | n2 | H statistic | p-value | Adj. p-value |
|---|---|---|---|---|---|---|
| ID 13 | ID 14 | 68 | 68 | 288 | 1,29E-18 | 8,51E-17 |
| ID 13 | ID 15 | 68 | 67 | 811 | 1,09E-10 | 7,19E-09 |
| ID 13 | ID 16 | 68 | 68 | 808 | 6,00E-11 | 3,96E-09 |
| ID 13 | ID 17 | 68 | 68 | 847 | 1,84E-10 | 1,21E-08 |
| ID 13 | ID 18 | 68 | 56 | 1251 | 0,001 | 0,069 |
| ID 13 | ID 19 | 68 | 38 | 832 | 0,002 | 0,163 |
| ID 13 | ID 20 | 68 | 68 | 324 | 5,14E-18 | 3,39E-16 |
| ID 13 | ID 21 | 68 | 68 | 333 | 7,24E-18 | 4,78E-16 |
| ID 13 | ID 22 | 68 | 68 | 950 | 3,11E-09 | 2,05E-07 |
| ID 13 | ID 23 | 68 | 66 | 890 | 1,71E-09 | 1,13E-07 |
| ID 13 | ID 24 | 68 | 68 | 123 | 1,65E-21 | 1,09E-19 |
| ID 14 | ID 15 | 68 | 67 | 3685 | 6,03E-10 | 3,98E-08 |
| ID 14 | ID 16 | 68 | 68 | 2954 | 0,005 | 0,346 |
| ID 14 | ID 17 | 68 | 68 | 3667 | 3,74E-09 | 2,47E-07 |
| ID 14 | ID 18 | 68 | 56 | 3245 | 1,69E-11 | 1,12E-09 |
| ID 14 | ID 19 | 68 | 38 | 2318 | 1,42E-11 | 9,37E-10 |
| ID 14 | ID 20 | 68 | 68 | 2910 | 0,009 | 0,614 |
| ID 14 | ID 21 | 68 | 68 | 2396 | 0,716 | 1 |
| ID 14 | ID 22 | 68 | 68 | 3636 | 8,40E-09 | 5,54E-07 |
| ID 14 | ID 23 | 68 | 66 | 3550 | 6,25E-09 | 4,12E-07 |
| ID 14 | ID 24 | 68 | 68 | 1670 | 0,005 | 0,346 |
| ID 15 | ID 16 | 67 | 68 | 1660 | 0,007 | 0,434 |
| ID 15 | ID 17 | 67 | 68 | 2206 | 0,753 | 1 |
| ID 15 | ID 18 | 67 | 56 | 2289 | 0,036 | 1 |
| ID 15 | ID 19 | 67 | 38 | 1745 | 0,002 | 0,11 |
| ID 15 | ID 20 | 67 | 68 | 1192 | 1,78E-06 | 0,000117 |
| ID 15 | ID 21 | 67 | 68 | 932 | 3,20E-09 | 2,11E-07 |
| ID 15 | ID 22 | 67 | 68 | 2237 | 0,859 | 1 |
| ID 15 | ID 23 | 67 | 66 | 2161 | 0,824 | 1 |
| ID 15 | ID 24 | 67 | 68 | 395 | 1,19E-16 | 7,85E-15 |
| ID 16 | ID 17 | 68 | 68 | 2909 | 0,009 | 0,622 |
| ID 16 | ID 18 | 68 | 56 | 2703 | 6,09E-05 | 0,004 |
| ID 16 | ID 19 | 68 | 38 | 1925 | 3,09E-05 | 0,002 |
| ID 16 | ID 20 | 68 | 68 | 2142 | 0,461 | 1 |
| ID 16 | ID 21 | 68 | 68 | 1730 | 0,011 | 0,752 |
| ID 16 | ID 22 | 68 | 68 | 2907 | 0,01 | 0,638 |
| ID 16 | ID 23 | 68 | 66 | 2806 | 0,013 | 0,825 |
| ID 16 | ID 24 | 68 | 68 | 1106 | 1,55E-07 | 1,02E-05 |
| ID 17 | ID 18 | 68 | 56 | 2348 | 0,026 | 1 |
| ID 17 | ID 19 | 68 | 38 | 1789 | 0,001 | 0,071 |
| ID 17 | ID 20 | 68 | 68 | 1289 | 8,58E-06 | 0,000566 |
| ID 17 | ID 21 | 68 | 68 | 1018 | 1,80E-08 | 1,19E-06 |
| ID 17 | ID 22 | 68 | 68 | 2322 | 0,967 | 1 |
| ID 17 | ID 23 | 68 | 66 | 2262 | 0,938 | 1 |
| ID 17 | ID 24 | 68 | 68 | 468 | 1,03E-15 | 6,80E-14 |
| ID 18 | ID 19 | 56 | 38 | 1162 | 0,453 | 1 |

| | | | | | | |
|---|---|---|---|---|---|---|
| ID 18 | ID 20 | 56 | 68 | 772 | 1,34E-08 | 8,84E-07 |
| ID 18 | ID 21 | 56 | 68 | 608 | 7,79E-11 | 5,14E-09 |
| ID 18 | ID 22 | 56 | 68 | 1477 | 0,032 | 1 |
| ID 18 | ID 23 | 56 | 66 | 1440 | 0,036 | 1 |
| ID 18 | ID 24 | 56 | 68 | 248 | 9,39E-17 | 6,20E-15 |
| ID 19 | ID 20 | 38 | 68 | 353 | 6,30E-10 | 4,16E-08 |
| ID 19 | ID 21 | 38 | 68 | 313 | 1,15E-10 | 7,59E-09 |
| ID 19 | ID 22 | 38 | 68 | 839 | 0,003 | 0,189 |
| ID 19 | ID 23 | 38 | 66 | 785 | 0,002 | 0,103 |
| ID 19 | ID 24 | 38 | 68 | 98 | 3,76E-15 | 2,48E-13 |
| ID 20 | ID 21 | 68 | 68 | 1797 | 0,025 | 1 |
| ID 20 | ID 22 | 68 | 68 | 3294 | 1,94E-05 | 0,001 |
| ID 20 | ID 23 | 68 | 66 | 3230 | 1,16E-05 | 0,000766 |
| ID 20 | ID 24 | 68 | 68 | 1054 | 4,42E-08 | 2,92E-06 |
| ID 21 | ID 22 | 68 | 68 | 3582 | 3,29E-08 | 2,17E-06 |
| ID 21 | ID 23 | 68 | 66 | 3477 | 4,13E-08 | 2,73E-06 |
| ID 21 | ID 24 | 68 | 68 | 1594 | 0,002 | 0,118 |
| ID 22 | ID 23 | 68 | 66 | 2273 | 0,899 | 1 |
| ID 22 | ID 24 | 68 | 68 | 493 | 2,48E-15 | 1,64E-13 |
| ID 23 | ID 24 | 66 | 68 | 464 | 2,39E-15 | 1,58E-13 |

**Supplementary Table 3. Differences between the running velocity (m/s per day) between the PAW mice.**

| Animal ID 1 | Animal ID 2 | n1 | n2 | H statistic | p-value | Adj. p-value |
|---|---|---|---|---|---|---|
| ID 13 | ID 14 | 68 | 68 | 195 | 3,21E-20 | 2,12E-18 |
| ID 13 | ID 15 | 68 | 67 | 1437 | 0,000217 | 0,014 |
| ID 13 | ID 16 | 68 | 68 | 748 | 1,01E-11 | 6,67E-10 |
| ID 13 | ID 17 | 68 | 68 | 1588 | 0,002 | 0,108 |
| ID 13 | ID 18 | 68 | 56 | 2198 | 0,141 | 1 |
| ID 13 | ID 19 | 68 | 38 | 791 | 0,000976 | 0,064 |
| ID 13 | ID 20 | 68 | 68 | 472 | 1,18E-15 | 7,79E-14 |
| ID 13 | ID 21 | 68 | 68 | 1217 | 1,90E-06 | 0,000125 |
| ID 13 | ID 22 | 68 | 68 | 584 | 5,53E-14 | 3,65E-12 |
| ID 13 | ID 23 | 68 | 66 | 962 | 1,18E-08 | 7,79E-07 |
| ID 13 | ID 24 | 68 | 68 | 211 | 6,13E-20 | 4,05E-18 |
| ID 14 | ID 15 | 68 | 67 | 4139 | 2,66E-16 | 1,76E-14 |
| ID 14 | ID 16 | 68 | 68 | 4040 | 5,53E-14 | 3,65E-12 |
| ID 14 | ID 17 | 68 | 68 | 4240 | 4,90E-17 | 3,23E-15 |
| ID 14 | ID 18 | 68 | 56 | 3689 | 3,25E-19 | 2,14E-17 |
| ID 14 | ID 19 | 68 | 38 | 2391 | 4,59E-13 | 3,03E-11 |
| ID 14 | ID 20 | 68 | 68 | 3709 | 1,22E-09 | 8,05E-08 |
| ID 14 | ID 21 | 68 | 68 | 4282 | 1,02E-17 | 6,73E-16 |
| ID 14 | ID 22 | 68 | 68 | 3812 | 6,74E-11 | 4,45E-09 |
| ID 14 | ID 23 | 68 | 66 | 3908 | 1,33E-13 | 8,78E-12 |
| ID 14 | ID 24 | 68 | 68 | 3357 | 5,47E-06 | 0,000361 |
| ID 15 | ID 16 | 67 | 68 | 1567 | 0,002 | 0,117 |
| ID 15 | ID 17 | 67 | 68 | 2352 | 0,746 | 1 |
| ID 15 | ID 18 | 67 | 56 | 2857 | 6,37E-07 | 4,20E-05 |
| ID 15 | ID 19 | 67 | 38 | 1385 | 0,457 | 1 |
| ID 15 | ID 20 | 67 | 68 | 1011 | 2,50E-08 | 1,65E-06 |
| ID 15 | ID 21 | 67 | 68 | 2325 | 0,838 | 1 |
| ID 15 | ID 22 | 67 | 68 | 1208 | 2,52E-06 | 0,000166 |
| ID 15 | ID 23 | 67 | 66 | 1716 | 0,026 | 1 |
| ID 15 | ID 24 | 67 | 68 | 629 | 4,03E-13 | 2,66E-11 |
| ID 16 | ID 17 | 68 | 68 | 3077 | 0,000877 | 0,058 |
| ID 16 | ID 18 | 68 | 56 | 3436 | 1,48E-14 | 9,77E-13 |
| ID 16 | ID 19 | 68 | 38 | 1816 | 0,000563 | 0,037 |
| ID 16 | ID 20 | 68 | 68 | 1595 | 0,002 | 0,12 |
| ID 16 | ID 21 | 68 | 68 | 3158 | 0,000233 | 0,015 |
| ID 16 | ID 22 | 68 | 68 | 1810 | 0,029 | 1 |
| ID 16 | ID 23 | 68 | 66 | 2402 | 0,483 | 1 |
| ID 16 | ID 24 | 68 | 68 | 1044 | 3,46E-08 | 2,28E-06 |
| ID 17 | ID 18 | 68 | 56 | 2761 | 1,70E-05 | 0,001 |
| ID 17 | ID 19 | 68 | 38 | 1295 | 0,987 | 1 |
| ID 17 | ID 20 | 68 | 68 | 1059 | 5,00E-08 | 3,30E-06 |
| ID 17 | ID 21 | 68 | 68 | 2209 | 0,656 | 1 |
| ID 17 | ID 22 | 68 | 68 | 1240 | 3,11E-06 | 0,000205 |
| ID 17 | ID 23 | 68 | 66 | 1690 | 0,014 | 0,911 |
| ID 17 | ID 24 | 68 | 68 | 664 | 7,47E-13 | 4,93E-11 |
| ID 18 | ID 19 | 56 | 38 | 439 | 1,50E-06 | 9,90E-05 |

| | | | | | | |
|---|---|---|---|---|---|---|
| ID 18 | ID 20 | 56 | 68 | 210 | 1,85E-17 | 1,22E-15 |
| ID 18 | ID 21 | 56 | 68 | 655 | 3,64E-10 | 2,40E-08 |
| ID 18 | ID 22 | 56 | 68 | 295 | 6,68E-16 | 4,41E-14 |
| ID 18 | ID 23 | 56 | 66 | 524 | 1,05E-11 | 6,93E-10 |
| ID 18 | ID 24 | 56 | 68 | 61 | 2,22E-20 | 1,47E-18 |
| ID 19 | ID 20 | 38 | 68 | 439 | 1,95E-08 | 1,29E-06 |
| ID 19 | ID 21 | 38 | 68 | 1189 | 0,5 | 1 |
| ID 19 | ID 22 | 38 | 68 | 560 | 1,44E-06 | 9,50E-05 |
| ID 19 | ID 23 | 38 | 66 | 857 | 0,007 | 0,491 |
| ID 19 | ID 24 | 38 | 68 | 233 | 3,09E-12 | 2,04E-10 |
| ID 20 | ID 21 | 68 | 68 | 3790 | 1,27E-10 | 8,38E-09 |
| ID 20 | ID 22 | 68 | 68 | 2526 | 0,353 | 1 |
| ID 20 | ID 23 | 68 | 66 | 3052 | 0,000326 | 0,022 |
| ID 20 | ID 24 | 68 | 68 | 1714 | 0,009 | 0,614 |
| ID 21 | ID 22 | 68 | 68 | 1059 | 5,00E-08 | 3,30E-06 |
| ID 21 | ID 23 | 68 | 66 | 1618 | 0,005 | 0,354 |
| ID 21 | ID 24 | 68 | 68 | 417 | 1,64E-16 | 1,08E-14 |
| ID 22 | ID 23 | 68 | 66 | 2843 | 0,008 | 0,51 |
| ID 22 | ID 24 | 68 | 68 | 1506 | 0,000455 | 0,03 |
| ID 23 | ID 24 | 66 | 68 | 970 | 1,45E-08 | 9,57E-07 |
