## Supplementary table 4 for "Mouse PAW: reverse-translating the FINGER multimodal lifestyle intervention enhances synaptic plasticity and cognition in adult wild type female mice"

| UniProt ID | Protein ID | Protein Name | Location | Type | Fold Change | p-value | FDR | Difference |
|---|---|---|---|---|---|---|---|---|
| Q61102 | ABCB7 | ATP binding cassette subfamily B member 7 | Cytoplasm | transporter | 1,53 | 2,43E-03 | 0,048 | UP |
| A0A571BG95 | ABR | ABR activator of RhoGEF and GTPase | Cytoplasm | other | 1,42 | 1,66E-03 | 0,048 | UP |
| Q5SWU9 | ACACA | acetyl-CoA carboxylase alpha | Cytoplasm | enzyme | 1,42 | 2,49E-03 | 0,048 | UP |
| Q8JZN5 | ACAD9 | acyl-CoA dehydrogenase family member 9 | Cytoplasm | enzyme | 1,26 | 3,32E-03 | 0,048 | UP |
| Q3V117 | ACLY | ATP citrate lyase | Cytoplasm | enzyme | 1,65 | 1,51E-03 | 0,048 | UP |
| Q99KI0 | ACO2 | aconitase 2 | Cytoplasm | enzyme | 1,26 | 2,61E-03 | 0,048 | UP |
| Q99PU5 | ACSBG1 | acyl-CoA synthetase bubblegum family member 1 | Cytoplasm | enzyme | 1,34 | 2,31E-03 | 0,048 | UP |
| Q3TGW0 | ACTR3 | actin related protein 3 | Plasma Membrane | other | 1,28 | 2,62E-03 | 0,048 | UP |
| P51830 | ADCY9 | adenylate cyclase 9 | Plasma Membrane | enzyme | 1,38 | 3,97E-03 | 0,049 | UP |
| Q3UHD1 | ADGRB1 | adhesion G protein-coupled receptor B1 | Plasma Membrane | G-protein coupled receptor | 1,41 | 1,66E-03 | 0,048 | UP |
| Q3UYG0 | ADORA1 | adenosine A1 receptor | Plasma Membrane | G-protein coupled receptor | 1,41 | 6,32E-04 | 0,048 | UP |
| Q3UHD9 | AGAP2 | ArfGAP with GTPase domain, ankyrin repeat and PH domain 2 | Nucleus | enzyme | 1,48 | 3,02E-03 | 0,048 | UP |
| E9QKI4 | AKT1S1 | AKT1 substrate 1 | Cytoplasm | other | 0,75 | 1,05E-03 | 0,048 | DOWN |
| A0A0G2JEG8 | AMPH | amphiphysin | Plasma Membrane | other | 0,79 | 3,14E-03 | 0,048 | DOWN |
| S4R2F3 | Ank2 | ankyrin 2, brain | Plasma Membrane | other | 1,36 | 4,13E-04 | 0,048 | UP |
| A2AHL1 | ANO3 | anoctamin 3 | Plasma Membrane | transporter | 0,07 | 3,83E-04 | 0,048 | DOWN |
| O35643 | AP1B1 | adaptor related protein complex 1 subunit beta 1 | Cytoplasm | other | 1,52 | 2,62E-03 | 0,048 | UP |
| Q3UKX8 | AP1G1 | adaptor related protein complex 1 subunit gamma 1 | Cytoplasm | other | 1,60 | 1,46E-03 | 0,048 | UP |
| P35585 | AP1M1 | adaptor related protein complex 1 subunit mu 1 | Cytoplasm | transporter | 1,39 | 2,63E-03 | 0,048 | UP |
| P17426 | AP2A1 | adaptor related protein complex 2 subunit alpha 1 | Cytoplasm | transporter | 1,71 | 2,04E-03 | 0,048 | UP |
| Q69ZW4 | AP2A2 | adaptor related protein complex 2 subunit alpha 2 | Cytoplasm | transporter | 1,52 | 3,09E-03 | 0,048 | UP |
| Q5SWR1 | Ap2b1 | adaptor-related protein complex 2, beta 1 subunit | Plasma Membrane | other | 1,63 | 1,67E-03 | 0,048 | UP |
| Q5FWI9 | AP2M1 | adaptor related protein complex 2 subunit mu 1 | Cytoplasm | other | 1,45 | 3,28E-03 | 0,048 | UP |
| P62743 | AP2S1 | adaptor related protein complex 2 subunit sigma 1 | Cytoplasm | transporter | 1,33 | 1,69E-03 | 0,048 | UP |
| Q9JME5 | AP3B2 | adaptor related protein complex 3 subunit beta 2 | Cytoplasm | other | 1,35 | 3,70E-03 | 0,049 | UP |
| O54774 | AP3D1 | adaptor related protein complex 3 subunit delta 1 | Cytoplasm | other | 1,65 | 7,89E-04 | 0,048 | UP |
| Q6GR78 | APP | amyloid beta precursor protein | Plasma Membrane | other | 0,74 | 3,10E-03 | 0,048 | DOWN |
| A0A0R4J0Z3 | AQP4 | aquaporin 4 | Plasma Membrane | transporter | 1,53 | 1,93E-03 | 0,048 | UP |
| Q6ZQ82 | ARHGAP26 | Rho GTPase activating protein 26 | Cytoplasm | other | 1,29 | 3,27E-03 | 0,048 | UP |
| H3BJX8 | ARHGEF2 | Rho/Rac guanine nucleotide exchange factor 2 | Cytoplasm | other | 1,43 | 2,26E-03 | 0,048 | UP |
| Q8BH66 | ATL1 | atlastin GTPase 1 | Cytoplasm | enzyme | 1,45 | 2,03E-03 | 0,048 | UP |
| Q5DTI2 | ATP2A2 | ATPase sarcoplasmic/endoplasmic reticulum Ca2+ transporting 2 | Cytoplasm | transporter | 1,45 | 1,59E-03 | 0,048 | UP |
| G5E829 | ATP2B1 | ATPase plasma membrane Ca2+ transporting 1 | Plasma Membrane | transporter | 1,56 | 1,73E-03 | 0,048 | UP |
| F8WHB1 | ATP2B2 | ATPase plasma membrane Ca2+ transporting 2 | Plasma Membrane | transporter | 1,45 | 1,67E-03 | 0,048 | UP |
| Q9DCX2 | ATP5PD | ATP synthase peripheral stalk subunit d | Cytoplasm | enzyme | 0,79 | 3,64E-03 | 0,048 | DOWN |
| Q9Z1G4 | ATP6V0A1 | ATPase H+ transporting V0 subunit a1 | Cytoplasm | transporter | 1,71 | 8,77E-04 | 0,048 | UP |
| Q3U861 | ATP6V1D | ATPase H+ transporting V1 subunit D | Cytoplasm | transporter | 0,77 | 7,83E-04 | 0,048 | DOWN |
| Q8BVE3 | ATP6V1H | ATPase H+ transporting V1 subunit H | Cytoplasm | transporter | 1,32 | 3,60E-03 | 0,048 | UP |
| O35607 | BMPR2 | bone morphogenetic protein receptor type 2 | Plasma Membrane | kinase | 1,44 | 4,57E-04 | 0,048 | UP |
| Q920P3 | BRINP1 | BMP/retinoic acid inducible neural specific 1 | Nucleus | peptidase | 1,40 | 3,86E-03 | 0,049 | UP |
| Q3ZB20 | CACNG2 | calcium voltage-gated channel auxiliary subunit gamma 2 | Plasma Membrane | ion channel | 2,10 | 3,13E-03 | 0,048 | UP |
| Q80TJ1 | CADPS | calcium dependent secretion activator | Plasma Membrane | other | 1,56 | 1,83E-03 | 0,048 | UP |
| B2MWM9 | CALR | calreticulin | Cytoplasm | transcription regulator | 0,78 | 2,85E-03 | 0,048 | DOWN |
| Q80TN1 | CAMK2A | calcium/calmodulin dependent protein kinase II alpha | Cytoplasm | kinase | 1,83 | 2,51E-03 | 0,048 | UP |
| A0A0G2JGS4 | CAMK2D | calcium/calmodulin dependent protein kinase II delta | Cytoplasm | kinase | 1,32 | 8,09E-04 | 0,048 | UP |
| A0A2I3BQP6 | CAMK2G | calcium/calmodulin dependent protein kinase II gamma | Cytoplasm | kinase | 1,52 | 1,26E-03 | 0,048 | UP |
| Q6ZQ38 | CAND1 | cullin associated and neddylation dissociated 1 | Cytoplasm | transcription regulator | 1,46 | 1,14E-03 | 0,048 | UP |
| Q8VDP4 | CCAR2 | cell cycle and apoptosis regulator 2 | Cytoplasm | peptidase | 1,30 | 2,89E-03 | 0,048 | UP |
| Q8C1Y8 | CCZ1/CCZ1B | CCZ1 homolog, vacuolar protein trafficking and biogenesis associated | Cytoplasm | other | 1,33 | 3,85E-03 | 0,049 | UP |
| Q99L43 | CDS2 | CDP-diacylglycerol synthase 2 | Cytoplasm | enzyme | 1,73 | 2,21E-03 | 0,048 | UP |
| Q9CXS4 | CENPV | centromere protein V | Nucleus | other | 1,25 | 4,59E-04 | 0,048 | UP |
| Q08EB5 | CLASP2 | cytoplasmic linker associated protein 2 | Cytoplasm | other | 1,46 | 3,20E-03 | 0,048 | UP |
| A0A0R4J0B4 | CMAS | cytidine monophosphate N-acetylneuraminic acid synthetase | Nucleus | enzyme | 1,31 | 1,58E-03 | 0,048 | UP |
| Q9D1A2 | CNDP2 | carnosine dipeptidase 2 | Cytoplasm | peptidase | 0,76 | 1,86E-03 | 0,048 | DOWN |
| F8WHL2 | COPA | COPI coat complex subunit alpha | Cytoplasm | transporter | 1,46 | 1,77E-03 | 0,048 | UP |
| Q9JIF7 | COPB1 | COPI coat complex subunit beta 1 | Cytoplasm | transporter | 1,36 | 3,99E-03 | 0,049 | UP |
| Q9QZE5 | COPG1 | COPI coat complex subunit gamma 1 | Cytoplasm | transporter | 1,39 | 1,14E-03 | 0,048 | UP |
| Q8K1Z0 | COQ9 | coenzyme Q9 | Cytoplasm | other | 0,80 | 3,45E-03 | 0,048 | DOWN |
| Q3TEU8 | CORO1C | coronin 1C | Cytoplasm | other | 1,39 | 6,19E-04 | 0,048 | UP |
| Q8BH44 | CORO2B | coronin 2B | Plasma Membrane | other | 1,49 | 2,89E-03 | 0,048 | UP |
| Q5ND51 | CRK | CRK proto-oncogene, adaptor protein | Cytoplasm | other | 0,80 | 3,24E-03 | 0,048 | DOWN |
| Q4FJX4 | CSRP1 | cysteine and glycine rich protein 1 | Nucleus | other | 0,73 | 9,51E-04 | 0,048 | DOWN |
| G3X914 | CUL5 | cullin 5 | Nucleus | ion channel | 1,43 | 1,43E-03 | 0,048 | UP |
| Q7TMB8 | CYFIP1 | cytoplasmic FMR1 interacting protein 1 | Cytoplasm | translation regulator | 1,57 | 2,73E-03 | 0,048 | UP |
| Q5SQX6 | CYFIP2 | cytoplasmic FMR1 interacting protein 2 | Cytoplasm | other | 1,77 | 1,35E-03 | 0,048 | UP |
| Q3US73 | CYP46A1 | cytochrome P450 family 46 subfamily A member 1 | Cytoplasm | enzyme | 1,41 | 1,66E-03 | 0,048 | UP |
| Q921M7 | CYRIB | CYFIP related Rac1 interactor B | Extracellular Space | other | 1,27 | 1,98E-03 | 0,048 | UP |
| P61804 | DAD1 | defender against cell death 1 | Cytoplasm | other | 1,37 | 2,42E-03 | 0,048 | UP |
| Q9JLM8 | DCLK1 | doublecortin like kinase 1 | Plasma Membrane | kinase | 1,26 | 2,41E-03 | 0,048 | UP |
| O08788 | DCTN1 | dynactin subunit 1 | Cytoplasm | other | 1,35 | 2,41E-03 | 0,048 | UP |
| Q9CWS0 | DDAH1 | dimethylarginine dimethylaminohydrolase 1 | Cytoplasm | enzyme | 0,78 | 1,00E-03 | 0,048 | DOWN |
| Q3U1J4 | DDB1 | damage specific DNA binding protein 1 | Nucleus | other | 1,33 | 1,54E-03 | 0,048 | UP |
| O54734 | DDOST | dolichyl-diphosphooligosaccharide--protein glycosyltransferase non-catalytic subunit | Cytoplasm | enzyme | 1,31 | 3,71E-03 | 0,049 | UP |
| A0A1S6GWH2 | DDX39B | DExD-box helicase 39B | Nucleus | enzyme | 1,25 | 2,33E-03 | 0,048 | UP |
| Q3V0Z8 | DDX5 | DEAD-box helicase 5 | Nucleus | enzyme | 1,35 | 2,59E-03 | 0,048 | UP |
| D3YXJ0 | DGKH | diacylglycerol kinase eta | Cytoplasm | kinase | 1,35 | 2,12E-03 | 0,048 | UP |
| A2AHK0 | DGKZ | diacylglycerol kinase zeta | Cytoplasm | kinase | 1,29 | 3,00E-03 | 0,048 | UP |
| A0A338P6I6 | DLG4 | discs large MAGUK scaffold protein 4 | Plasma Membrane | kinase | 1,37 | 1,07E-03 | 0,048 | UP |
| Q8BPN8 | DMXL2 | Dmx like 2 | Cytoplasm | other | 1,66 | 1,22E-03 | 0,048 | UP |
| Q9QYJ0 | DNAJA2 | DnaJ heat shock protein family (Hsp40) member A2 | Nucleus | enzyme | 0,78 | 2,94E-03 | 0,048 | DOWN |
| A0A0J9YUN4 | DNM1 | dynamin 1 | Cytoplasm | enzyme | 1,40 | 4,04E-03 | 0,049 | UP |
| Q9JHU4 | DYNC1H1 | dynein cytoplasmic 1 heavy chain 1 | Cytoplasm | peptidase | 1,76 | 1,43E-03 | 0,048 | UP |
| Q542J7 | DYNLT3 | dynein light chain Tctex-type 3 | Cytoplasm | other | 0,79 | 1,04E-03 | 0,048 | DOWN |
| P10630 | EIF4A2 | eukaryotic translation initiation factor 4A2 | Cytoplasm | translation regulator | 1,26 | 3,64E-03 | 0,048 | UP |
| Q8VEH5 | EPM2AIP1 | EPM2A interacting protein 1 | Cytoplasm | other | 1,25 | 2,01E-03 | 0,048 | UP |
| Q8CGC7 | EPRS1 | glutamyl-prolyl-tRNA synthetase 1 | Cytoplasm | enzyme | 1,56 | 2,81E-03 | 0,048 | UP |
| Q9D4H1 | EXOC2 | exocyst complex component 2 | Cytoplasm | transporter | 1,39 | 1,63E-03 | 0,048 | UP |
| Q8K0E2 | EXOC3 | exocyst complex component 3 | Plasma Membrane | transporter | 1,31 | 2,49E-03 | 0,048 | UP |
| Q6PGF7 | EXOC8 | exocyst complex component 8 | Plasma Membrane | other | 1,27 | 1,24E-03 | 0,048 | UP |
| O08914 | FAAH | fatty acid amide hydrolase | Plasma Membrane | enzyme | 1,38 | 1,74E-03 | 0,048 | UP |
| F8VPU2 | FARP1 | FERM, ARH/RhoGEF and pleckstrin domain protein 1 | Plasma Membrane | other | 1,33 | 2,08E-03 | 0,048 | UP |
| P19096 | FASN | fatty acid synthase | Cytoplasm | enzyme | 1,62 | 1,08E-03 | 0,048 | UP |
| Q9WV18 | GABBR1 | gamma-aminobutyric acid type B receptor subunit 1 | Plasma Membrane | G-protein coupled receptor | 1,52 | 3,45E-03 | 0,048 | UP |
| Q3URD4 | GABRA1 | gamma-aminobutyric acid type A receptor subunit alpha1 | Plasma Membrane | ion channel | 1,49 | 2,02E-03 | 0,048 | UP |
| Q8BHJ7 | GABRA5 | gamma-aminobutyric acid type A receptor subunit alpha5 | Plasma Membrane | ion channel | 1,37 | 3,75E-04 | 0,048 | UP |
| P22723 | GABRG2 | gamma-aminobutyric acid type A receptor subunit gamma2 | Plasma Membrane | ion channel | 1,46 | 3,55E-04 | 0,048 | UP |
| Q88741 | GDAP1 | ganglioside induced differentiation associated protein 1 | Cytoplasm | other | 1,31 | 3,20E-03 | 0,048 | UP |
| Q68FF6 | GIT1 | GIT ArfGAP 1 | Cytoplasm | other | 1,27 | 3,25E-03 | 0,048 | UP |
| Q64516 | Gk | glycerol kinase | Cytoplasm | kinase | 1,31 | 3,56E-04 | 0,048 | UP |
| Q8C0L9 | GPCPD1 | glycerophosphocholine phosphodiesterase 1 | Cytoplasm | enzyme | 1,25 | 3,55E-03 | 0,048 | UP |
| C9K0Y4 | GRIA1 | glutamate ionotropic receptor AMPA type subunit 1 | Plasma Membrane | ion channel | 1,67 | 2,64E-03 | 0,048 | UP |
| E9QKC0 | GRIA2 | glutamate ionotropic receptor AMPA type subunit 2 | Plasma Membrane | ion channel | 1,57 | 1,94E-03 | 0,048 | UP |
| A2VDF5 | GRIA3 | glutamate ionotropic receptor AMPA type subunit 3 | Plasma Membrane | ion channel | 1,56 | 1,11E-03 | 0,048 | UP |

| | | | | | | | | |
|---|---|---|---|---|---|---|---|---|
| A2AI21 | GRIN1 | glutamate ionotropic receptor NMDA type subunit 1 | Plasma Membrane | ion channel | 1,58 | 1,25E-03 | 0,048 | UP |
| G3X9V4 | GRIN2B | glutamate ionotropic receptor NMDA type subunit 2B | Plasma Membrane | ion channel | 1,50 | 7,72E-04 | 0,048 | UP |
| Q3UVX5 | GRM5 | glutamate metabotropic receptor 5 | Plasma Membrane | G-protein coupled receptor | 1,32 | 3,86E-03 | 0,049 | UP |
| O09131 | GSTO1 | glutathione S-transferase omega 1 | Cytoplasm | enzyme | 0,79 | 3,05E-03 | 0,048 | DOWN |
| Q9D3B1 | HACD2 | 3-hydroxyacyl-CoA dehydratase 2 | Cytoplasm | phosphatase | 2,18 | 7,29E-04 | 0,048 | UP |
| Q8K2C9 | HACD3 | 3-hydroxyacyl-CoA dehydratase 3 | Cytoplasm | enzyme | 1,47 | 2,96E-03 | 0,048 | UP |
| Q3UMU9 | HDGFL2 | HDGF like 2 | Nucleus | other | 0,74 | 3,64E-03 | 0,048 | DOWN |
| Q6ZQ77 | HIP1R | huntingtin interacting protein 1 related | Cytoplasm | other | 1,31 | 2,33E-03 | 0,048 | UP |
| P17710 | HK1 | hexokinase 1 | Cytoplasm | kinase | 1,48 | 1,94E-03 | 0,048 | UP |
| Q9CX86 | HNRNPA0 | heterogeneous nuclear ribonucleoprotein A0 | Nucleus | other | 1,39 | 2,62E-03 | 0,048 | UP |
| G3X9H5 | HTT | huntingtin | Cytoplasm | transcription regulator | 1,40 | 9,62E-04 | 0,048 | UP |
| Q9JKR6 | HYOU1 | hypoxia up-regulated 1 | Cytoplasm | other | 0,77 | 3,64E-03 | 0,048 | DOWN |
| Q8BIJ6 | IARS2 | isoleucyl-tRNA synthetase 2, mitochondrial | Cytoplasm | enzyme | 1,28 | 6,90E-04 | 0,048 | UP |
| Q8BKC5 | IPO5 | importin 5 | Nucleus | transporter | 1,35 | 2,51E-03 | 0,048 | UP |
| Q9EPL8 | IPO7 | importin 7 | Nucleus | transporter | 1,39 | 3,85E-03 | 0,049 | UP |
| Q3UQ44 | IQGAP2 | IQ motif containing GTPase activating protein 2 | Cytoplasm | other | 1,45 | 9,19E-04 | 0,048 | UP |
| A0A1D5RM83 | IQSEC1 | IQ motif and Sec7 domain ArfGEF 1 | Cytoplasm | other | 1,63 | 1,20E-03 | 0,048 | UP |
| Q3TES0 | IQSEC3 | IQ motif and Sec7 domain ArfGEF 3 | Cytoplasm | other | 1,57 | 1,93E-03 | 0,048 | UP |
| P11881 | ITPR1 | inositol 1,4,5-trisphosphate receptor type 1 | Cytoplasm | ion channel | 1,61 | 3,59E-03 | 0,048 | UP |
| Q9D8B7 | JAM3 | junctional adhesion molecule 3 | Plasma Membrane | other | 0,76 | 3,58E-03 | 0,048 | DOWN |
| A2CG49 | KALRN | kalirin RhoGEF kinase | Cytoplasm | kinase | 1,51 | 2,75E-03 | 0,048 | UP |
| Q9Z0V2 | KCND2 | potassium voltage-gated channel subfamily D member 2 | Plasma Membrane | ion channel | 1,46 | 1,19E-03 | 0,048 | UP |
| A0A087WQE8 | KIF1A | kinesin family member 1A | Cytoplasm | other | 1,34 | 2,23E-03 | 0,048 | UP |
| Q9JJK2 | LANCL2 | LanC like glutathione S-transferase 2 | Plasma Membrane | other | 1,31 | 3,11E-03 | 0,048 | UP |
| A0A1B0GSX0 | LDHA | lactate dehydrogenase A | Cytoplasm | enzyme | 1,32 | 3,77E-03 | 0,049 | UP |
| Q9JIA1 | LGI1 | leucine rich glioma inactivated 1 | Plasma Membrane | other | 1,40 | 2,79E-03 | 0,048 | UP |
| Q9D7I5 | LHPP | phospholysine phosphohistidine inorganic pyrophosphate phosphatase | Cytoplasm | phosphatase | 0,72 | 1,74E-03 | 0,048 | DOWN |
| O88952 | LIN7C | lin-7 homolog C, crumbs cell polarity complex component | Cytoplasm | other | 0,67 | 1,18E-03 | 0,048 | DOWN |
| A0A087WNU6 | LRRFIP1 | LRR binding FLII interacting protein 1 | Cytoplasm | transcription regulator | 0,78 | 3,03E-03 | 0,048 | DOWN |
| Q9CYI4 | LUC7L | LUC7 like | Nucleus | other | 0,77 | 8,04E-04 | 0,048 | DOWN |
| A2AGQ4 | MADD | MAP kinase activating death domain | Cytoplasm | other | 1,39 | 3,73E-03 | 0,049 | UP |
| P20917 | MAG | myelin associated glycoprotein | Plasma Membrane | other | 0,69 | 3,38E-03 | 0,048 | DOWN |
| Q2KHK7 | MAL2 | mal, T cell differentiation protein 2 | Plasma Membrane | transporter | 1,48 | 2,70E-03 | 0,048 | UP |
| Q9CXI5 | MANF | mesencephalic astrocyte derived neurotrophic factor | Extracellular Space | other | 0,68 | 3,86E-03 | 0,049 | DOWN |
| Q3U2X5 | MAOA | monoamine oxidase A | Cytoplasm | enzyme | 1,37 | 1,31E-03 | 0,048 | UP |
| Q14BB9 | MAP6D1 | MAP6 domain containing 1 | Cytoplasm | other | 1,26 | 1,13E-03 | 0,048 | UP |
| Q8BMF3 | ME3 | malic enzyme 3 | Cytoplasm | enzyme | 1,33 | 1,92E-03 | 0,048 | UP |
| Q3TEX7 | MFN2 | mitofusin 2 | Cytoplasm | enzyme | 1,47 | 3,42E-03 | 0,048 | UP |
| G3X9G2 | MINK1 | misshapen like kinase 1 | Cytoplasm | kinase | 1,54 | 4,80E-04 | 0,048 | UP |
| Q8R3Q6 | MIX23 | mitochondrial matrix import factor 23 | Cytoplasm | other | 0,76 | 3,94E-03 | 0,049 | DOWN |
| Q9D023 | MPC2 | mitochondrial pyruvate carrier 2 | Plasma Membrane | transporter | 1,32 | 2,66E-03 | 0,048 | UP |
| Q61474 | MSI1 | musashi RNA binding protein 1 | Cytoplasm | other | 0,78 | 2,76E-03 | 0,048 | DOWN |
| Q9D050 | MTCH2 | mitochondrial carrier 2 | Cytoplasm | other | 1,55 | 1,73E-03 | 0,048 | UP |
| A0A075DCI1 | MT-ND4 | NADH dehydrogenase subunit 4 | Cytoplasm | enzyme | 1,91 | 1,50E-03 | 0,048 | UP |
| A0A075DCB1 | MT-ND5 | NADH dehydrogenase subunit 5 | Cytoplasm | enzyme | 2,06 | 1,17E-03 | 0,048 | UP |
| Q3UH59 | MYH10 | myosin heavy chain 10 | Cytoplasm | enzyme | 1,77 | 9,28E-04 | 0,048 | UP |
| K3W4R2 | MYH14 | myosin heavy chain 14 | Extracellular Space | enzyme | 1,78 | 3,69E-03 | 0,049 | UP |
| Q8VDD5 | MYH9 | myosin heavy chain 9 | Cytoplasm | enzyme | 1,82 | 1,11E-03 | 0,048 | UP |
| E9QAX2 | MYO18A | myosin XVIIIA | Cytoplasm | other | 1,50 | 7,12E-04 | 0,048 | UP |
| D3Z4J3 | MYO5A | myosin VA | Cytoplasm | enzyme | 1,96 | 1,11E-03 | 0,048 | UP |
| Q3UK27 | NAE1 | NEDD8 activating enzyme E1 subunit 1 | Cytoplasm | enzyme | 1,43 | 2,02E-03 | 0,048 | UP |
| Q9EPN1 | NBEA | neurobeachin | Cytoplasm | other | 1,65 | 1,48E-03 | 0,048 | UP |
| A0A0R4IZX5 | NCAN | neurocan | Extracellular Space | other | 0,79 | 3,01E-03 | 0,048 | DOWN |
| Q9Z0E0 | NCDN | neurochondrin | Cytoplasm | other | 1,69 | 1,11E-03 | 0,048 | UP |
| Q8BLF1 | NCEH1 | neutral cholesterol ester hydrolase 1 | Plasma Membrane | enzyme | 1,34 | 2,90E-03 | 0,048 | UP |
| A2AS98 | NCKAP1 | NCK associated protein 1 | Plasma Membrane | other | 1,65 | 9,73E-04 | 0,048 | UP |
| Q9DC69 | NDUFA9 | NADH:ubiquinone oxidoreductase subunit A9 | Cytoplasm | enzyme | 1,46 | 3,96E-04 | 0,048 | UP |
| A0A494BAX0 | NEDD4L | NEDD4 like E3 ubiquitin protein ligase | Cytoplasm | enzyme | 1,25 | 1,90E-03 | 0,048 | UP |
| A0A087WPX3 | NFASC | neurofascin | Plasma Membrane | other | 0,80 | 1,69E-03 | 0,048 | DOWN |
| P30415 | NKTR | natural killer cell triggering receptor | Plasma Membrane | enzyme | 0,64 | 1,15E-03 | 0,048 | DOWN |
| Q3TYC2 | NTM | neurotrimin | Plasma Membrane | other | 0,75 | 3,52E-03 | 0,048 | DOWN |
| Q8CGY8 | OGT | O-linked N-acetylglucosamine (GlcNAc) transferase | Cytoplasm | enzyme | 1,45 | 2,96E-03 | 0,048 | UP |
| Q8K212 | PACS1 | phosphofurin acidic cluster sorting protein 1 | Cytoplasm | other | 1,25 | 1,19E-03 | 0,048 | UP |
| Q61206 | PAFAH1B2 | platelet activating factor acetylhydrolase 1b catalytic subunit 2 | Cytoplasm | enzyme | 0,76 | 1,78E-03 | 0,048 | DOWN |
| Q3TJC2 | PAFAH1B3 | platelet activating factor acetylhydrolase 1b catalytic subunit 3 | Cytoplasm | enzyme | 0,77 | 3,40E-03 | 0,048 | DOWN |
| F7BJK1 | PCDH1 | protocadherin 1 | Plasma Membrane | other | 0,80 | 3,30E-03 | 0,048 | DOWN |
| Q9QYX7 | PCLO | piccolo presynaptic cytomatrix protein | Cytoplasm | transporter | 1,25 | 2,68E-03 | 0,048 | UP |
| Q80Y09 | PDCD6IP | programmed cell death 6 interacting protein | Cytoplasm | other | 1,28 | 3,12E-03 | 0,048 | UP |
| Q8BKZ9 | PDHX | pyruvate dehydrogenase complex component X | Cytoplasm | enzyme | 0,78 | 1,83E-03 | 0,048 | DOWN |
| Q62048 | PEA15 | proliferation and apoptosis adaptor protein 15 | Cytoplasm | transporter | 0,73 | 8,08E-04 | 0,048 | DOWN |
| Q6ZWY7 | PFDN5 | prefoldin subunit 5 | Nucleus | transcription regulator | 0,73 | 1,55E-04 | 0,048 | DOWN |
| P12382 | PFKL | phosphofructokinase, liver type | Cytoplasm | kinase | 1,44 | 2,78E-03 | 0,048 | UP |
| P47857 | PFKM | phosphofructokinase, muscle | Cytoplasm | kinase | 1,56 | 7,94E-04 | 0,048 | UP |
| Q8C605 | PFKP | phosphofructokinase, platelet | Cytoplasm | kinase | 1,41 | 2,96E-03 | 0,048 | UP |
| Q3UAG2 | PGD | phosphogluconate dehydrogenase | Cytoplasm | enzyme | 1,39 | 1,35E-03 | 0,048 | UP |
| Q8CAA7 | PGM2L1 | phosphoglucomutase 2 like 1 | Cytoplasm | enzyme | 1,37 | 1,70E-03 | 0,048 | UP |
| Q3TH84 | PGRMC1 | progesterone receptor membrane component 1 | Plasma Membrane | transmembrane receptor | 0,77 | 2,02E-03 | 0,048 | DOWN |
| A0A1S6GWJ7 | PI4KA | phosphatidylinositol 4-kinase alpha | Cytoplasm | kinase | 1,60 | 1,32E-03 | 0,048 | UP |
| Q8BWR2 | PITHD1 | PITH domain containing 1 | Nucleus | other | 0,68 | 1,49E-03 | 0,048 | DOWN |
| O35954 | PITPNM1 | phosphatidylinositol transfer protein membrane associated 1 | Cytoplasm | transporter | 1,46 | 2,34E-03 | 0,048 | UP |
| Q6ZPQ6 | PITPNM2 | phosphatidylinositol transfer protein membrane associated 2 | Cytoplasm | enzyme | 1,51 | 9,70E-04 | 0,048 | UP |
| Q9QXS1 | PLEC | plectin | Cytoplasm | other | 1,52 | 1,46E-03 | 0,048 | UP |
| Q99JY8 | PLPP3 | phospholipid phosphatase 3 | Plasma Membrane | phosphatase | 0,78 | 3,77E-03 | 0,049 | DOWN |
| Q7TME0 | PLPPR4 | phospholipid phosphatase related 4 | Plasma Membrane | phosphatase | 1,28 | 1,12E-03 | 0,048 | UP |
| B1AX58 | PLS3 | plastin 3 | Cytoplasm | other | 1,33 | 4,56E-04 | 0,048 | UP |
| Q5DTM9 | PPFIA2 | PTPRF interacting protein alpha 2 | Plasma Membrane | phosphatase | 1,29 | 2,78E-03 | 0,048 | UP |
| C7G3P1 | PPFIA3 | PTPRF interacting protein alpha 3 | Plasma Membrane | phosphatase | 1,43 | 1,12E-03 | 0,048 | UP |
| P68181 | PRKACB | protein kinase cAMP-activated catalytic subunit beta | Cytoplasm | kinase | 1,35 | 1,08E-03 | 0,048 | UP |
| P16054 | PRKCE | protein kinase C epsilon | Cytoplasm | kinase | 1,33 | 3,07E-03 | 0,048 | UP |
| P63318 | PRKCG | protein kinase C gamma | Cytoplasm | kinase | 1,41 | 3,07E-03 | 0,048 | UP |
| J3QPG5 | PSAP | prosaposin | Extracellular Space | other | 0,64 | 2,03E-04 | 0,048 | DOWN |
| Q3TXS7 | PSMD1 | proteasome 26S subunit, non-ATPase 1 | Cytoplasm | other | 1,41 | 3,54E-03 | 0,048 | UP |
| Q99JI4 | PSMD6 | proteasome 26S subunit, non-ATPase 6 | Cytoplasm | enzyme | 1,34 | 2,16E-03 | 0,048 | UP |
| Q3TJG6 | PTGES3 | prostaglandin E synthase 3 | Cytoplasm | enzyme | 0,79 | 9,60E-04 | 0,048 | DOWN |
| Q66JR8 | Ptms | parathymosin | Cytoplasm | other | 0,75 | 1,92E-03 | 0,048 | DOWN |
| O35239 | PTPN9 | protein tyrosine phosphatase non-receptor type 9 | Cytoplasm | phosphatase | 1,37 | 1,33E-03 | 0,048 | UP |
| Q3UYH9 | PYGB | glycogen phosphorylase B | Cytoplasm | enzyme | 1,40 | 1,53E-03 | 0,048 | UP |
| A0A1D5RLG3 | RAB3GAP1 | RAB3 GTPase activating protein catalytic subunit 1 | Cytoplasm | other | 1,42 | 4,39E-04 | 0,048 | UP |
| P63321 | RALA | RAS like proto-oncogene A | Cytoplasm | enzyme | 0,74 | 1,98E-03 | 0,048 | DOWN |
| A0A0A6YWG7 | RAPGEF2 | Rap guanine nucleotide exchange factor 2 | Cytoplasm | other | 1,51 | 1,01E-03 | 0,048 | UP |
| Q9Z1S3 | RASGRP1 | RAS guanyl releasing protein 1 | Cytoplasm | other | 1,72 | 1,94E-03 | 0,048 | UP |
| P52760 | RIDA | reactive intermediate imine deaminase A homolog | Cytoplasm | enzyme | 0,69 | 2,88E-03 | 0,048 | DOWN |
| Q8VCT3 | RNPEP | arginyl aminopeptidase | Cytoplasm | peptidase | 1,34 | 3,36E-03 | 0,048 | UP |
| A0A1Y7VMN0 | ROCK2 | Rho associated coiled-coil containing protein kinase 2 | Cytoplasm | kinase | 1,31 | 1,82E-03 | 0,048 | UP |
| Q5BLK0 | RPL12 | ribosomal protein L12 | Nucleus | other | 1,27 | 1,19E-03 | 0,048 | UP |

| | | | | | | | | |
|---|---|---|---|---|---|---|---|---|
| P47963 | RPL13 | ribosomal protein L13 | Nucleus | other | 1,27 | 1,02E-03 | 0,048 | UP |
| Q9CWK0 | RPL14 | ribosomal protein L14 | Cytoplasm | other | 1,48 | 1,36E-03 | 0,048 | UP |
| Q3U7D2 | RPL15 | ribosomal protein L15 | Cytoplasm | other | 1,42 | 1,76E-03 | 0,048 | UP |
| A0A1D5RM85 | RPL18A | ribosomal protein L18a | Cytoplasm | other | 1,39 | 3,72E-03 | 0,049 | UP |
| Q3UW40 | RPL24 | ribosomal protein L24 | Cytoplasm | other | 1,46 | 3,47E-03 | 0,048 | UP |
| Q4FZH2 | RPL26 | ribosomal protein L26 | Cytoplasm | other | 1,64 | 1,67E-04 | 0,048 | UP |
| Q5BLJ9 | RPL27 | ribosomal protein L27 | Cytoplasm | other | 1,70 | 6,02E-04 | 0,048 | UP |
| P27659 | RPL3 | ribosomal protein L3 | Nucleus | other | 1,36 | 2,19E-03 | 0,048 | UP |
| Q9D1R9 | Rpl34 (includes others) | ribosomal protein L34 | Cytoplasm | other | 1,56 | 7,32E-04 | 0,048 | UP |
| Q6ZWX1 | RPL35A | ribosomal protein L35a | Cytoplasm | other | 1,62 | 9,73E-04 | 0,048 | UP |
| P47964 | Rpl36 | ribosomal protein L36 | Nucleus | other | 0,75 | 2,05E-04 | 0,048 | DOWN |
| P47911 | RPL6 | ribosomal protein L6 | Nucleus | other | 1,26 | 1,48E-03 | 0,048 | UP |
| Q80UT7 | RPL7A | ribosomal protein L7a | Cytoplasm | other | 1,38 | 6,25E-04 | 0,048 | UP |
| P62918 | RPL8 | ribosomal protein L8 | Cytoplasm | other | 1,37 | 2,76E-03 | 0,048 | UP |
| Q9DB79 | RPS11 | ribosomal protein S11 | Cytoplasm | other | 1,54 | 1,74E-03 | 0,048 | UP |
| Q5CZY9 | RPS16 | ribosomal protein S16 | Cytoplasm | other | 1,48 | 3,06E-04 | 0,048 | UP |
| P62908 | RPS3 | ribosomal protein S3 | Cytoplasm | enzyme | 1,32 | 2,58E-03 | 0,048 | UP |
| P97461 | RPS5 | ribosomal protein S5 | Cytoplasm | other | 0,76 | 7,85E-04 | 0,048 | DOWN |
| Q6ZWN5 | RPS9 | ribosomal protein S9 | Cytoplasm | translation regulator | 1,36 | 2,95E-03 | 0,048 | UP |
| A0A1Y7VJP1 | RYR2 | ryanodine receptor 2 | Plasma Membrane | ion channel | 2,25 | 2,86E-03 | 0,048 | UP |
| Q6PDS3 | SARM1 | sterile alpha and TIR motif containing 1 | Plasma Membrane | transmembrane receptor | 1,37 | 3,17E-03 | 0,048 | UP |
| Q6ZPE2 | SBF1 | SET binding factor 1 | Plasma Membrane | phosphatase | 1,57 | 1,59E-03 | 0,048 | UP |
| Q8C8N2 | SCAI | suppressor of cancer cell invasion | Nucleus | transcription regulator | 1,26 | 2,37E-04 | 0,048 | UP |
| A0A0R5RP28 | SCN2A | sodium voltage-gated channel alpha subunit 2 | Plasma Membrane | ion channel | 1,43 | 2,35E-03 | 0,048 | UP |
| Q925N0 | SFXN5 | sideroflexin 5 | Cytoplasm | transporter | 1,43 | 3,11E-03 | 0,048 | UP |
| E9QM38 | SLC12A2 | solute carrier family 12 member 2 | Plasma Membrane | transporter | 1,27 | 3,53E-03 | 0,048 | UP |
| A0A076FRG6 | SLC12A5 | solute carrier family 12 member 5 | Plasma Membrane | transporter | 1,58 | 8,07E-04 | 0,048 | UP |
| Q3TXX4 | SLC17A7 | solute carrier family 17 member 7 | Plasma Membrane | transporter | 1,39 | 2,29E-03 | 0,048 | UP |
| Q3UYK6 | SLC1A2 | solute carrier family 1 member 2 | Plasma Membrane | transporter | 1,39 | 3,01E-03 | 0,048 | UP |
| Q8BUN9 | SLC24A2 | solute carrier family 24 member 2 | Plasma Membrane | transporter | 1,35 | 2,57E-03 | 0,048 | UP |
| Q9CR62 | SLC25A11 | solute carrier family 25 member 11 | Cytoplasm | transporter | 1,39 | 1,23E-03 | 0,048 | UP |
| Q8BH59 | SLC25A12 | solute carrier family 25 member 12 | Cytoplasm | transporter | 1,52 | 3,92E-03 | 0,049 | UP |
| Q6GQS1 | SLC25A23 | solute carrier family 25 member 23 | Cytoplasm | transporter | 1,25 | 1,07E-03 | 0,048 | UP |
| Q3THU8 | SLC25A3 | solute carrier family 25 member 3 | Cytoplasm | transporter | 1,42 | 2,75E-03 | 0,048 | UP |
| P48962 | SLC25A4 | solute carrier family 25 member 4 | Cytoplasm | transporter | 1,44 | 4,03E-03 | 0,049 | UP |
| A2AKW0 | SLC25A51 | solute carrier family 25 member 51 | Cytoplasm | transporter | 1,57 | 9,36E-04 | 0,048 | UP |
| Q3TPL8 | SLC2A3 | solute carrier family 2 member 3 | Plasma Membrane | transporter | 1,58 | 1,27E-03 | 0,048 | UP |
| Q60738 | SLC30A1 | solute carrier family 30 member 1 | Plasma Membrane | transporter | 1,41 | 2,56E-03 | 0,048 | UP |
| O35633 | SLC32A1 | solute carrier family 32 member 1 | Plasma Membrane | transporter | 1,28 | 2,73E-03 | 0,048 | UP |
| Q8K2P7 | SLC38A1 | solute carrier family 38 member 1 | Plasma Membrane | transporter | 1,37 | 1,96E-03 | 0,048 | UP |
| B1AWV9 | SLC4A10 | solute carrier family 4 member 10 | Plasma Membrane | transporter | 1,44 | 5,80E-04 | 0,048 | UP |
| E9Q8N8 | SLC4A4 | solute carrier family 4 member 4 | Plasma Membrane | transporter | 1,47 | 3,19E-03 | 0,048 | UP |
| P31650 | SLC6A11 | solute carrier family 6 member 11 | Plasma Membrane | transporter | 1,47 | 3,84E-03 | 0,049 | UP |
| Q8BJI1 | SLC6A17 | solute carrier family 6 member 17 | Cytoplasm | transporter | 1,37 | 3,83E-03 | 0,049 | UP |
| Q9QXW9 | SLC7A8 | solute carrier family 7 member 8 | Plasma Membrane | transporter | 1,32 | 2,17E-03 | 0,048 | UP |
| Q5DTL5 | SLIT2 | slit guidance ligand 2 | Extracellular Space | other | 1,87 | 1,60E-03 | 0,048 | UP |
| Q9JJY3 | SMPD3 | sphingomyelin phosphodiesterase 3 | Cytoplasm | enzyme | 1,26 | 1,43E-03 | 0,048 | UP |
| Q9ER58 | SPOCK2 | SPARC (osteonectin), cwcv and kazal like domains proteoglycan 2 | Extracellular Space | other | 0,72 | 2,05E-03 | 0,048 | DOWN |
| Q62261 | SPTBN1 | spectrin beta, non-erythrocytic 1 | Plasma Membrane | other | 1,41 | 1,37E-03 | 0,048 | UP |
| Q68FG2 | SPTBN2 | spectrin beta, non-erythrocytic 2 | Cytoplasm | other | 1,61 | 1,61E-03 | 0,048 | UP |
| F8VPQ4 | SRGAP3 | SLIT-ROBO Rho GTPase activating protein 3 | Cytoplasm | other | 1,38 | 3,53E-03 | 0,048 | UP |
| A0A286YCG8 | SSR1 | signal sequence receptor subunit 1 | Cytoplasm | other | 0,75 | 7,34E-04 | 0,048 | DOWN |
| A0A1S6GWK0 | STT3A | STT3 oligosaccharyltransferase complex catalytic subunit A | Plasma Membrane | enzyme | 1,55 | 2,38E-03 | 0,048 | UP |
| Q5DQR4 | STXBP5L | syntaxin binding protein 5L | Cytoplasm | other | 1,45 | 2,15E-03 | 0,048 | UP |
| O88935 | SYN1 | synapsin I | Plasma Membrane | other | 1,29 | 5,09E-04 | 0,048 | UP |
| Q3KN99 | SYN3 | synapsin III | Plasma Membrane | other | 1,55 | 2,67E-03 | 0,048 | UP |
| F6SEU4 | SYNGAP1 | synaptic Ras GTPase activating protein 1 | Plasma Membrane | other | 1,58 | 1,06E-03 | 0,048 | UP |
| O55100 | SYNGR1 | synaptogyrin 1 | Plasma Membrane | transporter | 1,34 | 3,05E-03 | 0,048 | UP |
| F7BQW7 | SYNJ1 | synaptojanin 1 | Cytoplasm | phosphatase | 1,41 | 2,17E-03 | 0,048 | UP |
| F8VQ95 | TACC1 | transforming acidic coiled-coil containing protein 1 | Nucleus | transcription regulator | 0,80 | 1,72E-03 | 0,048 | DOWN |
| Q3UUG6 | TBC1D24 | TBC1 domain family member 24 | Cytoplasm | other | 1,48 | 2,37E-03 | 0,048 | UP |
| Q80VP0 | TECPR1 | tectonin beta-propeller repeat containing 1 | Cytoplasm | other | 1,29 | 1,97E-03 | 0,048 | UP |
| A0A5F8MQC8 | TECR | trans-2,3-enoyl-CoA reductase | Plasma Membrane | enzyme | 1,47 | 2,14E-03 | 0,048 | UP |
| E9PWD2 | THSD7A | thrombospondin type 1 domain containing 7A | Cytoplasm | other | 0,78 | 1,84E-03 | 0,048 | DOWN |
| B2RWB7 | TIGAR | TP53 induced glycolysis regulatory phosphatase | Cytoplasm | enzyme | 1,32 | 8,33E-04 | 0,048 | UP |
| A0A1L1SQ51 | TLN2 | talin 2 | Nucleus | other | 1,29 | 1,95E-03 | 0,048 | UP |
| Q9CR67 | TMEM33 | transmembrane protein 33 | Cytoplasm | other | 1,53 | 2,08E-03 | 0,048 | UP |
| Q8BFY9 | TNPO1 | transportin 1 | Nucleus | transporter | 1,39 | 2,21E-03 | 0,048 | UP |
| P17751 | TPI1 | triosephosphate isomerase 1 | Cytoplasm | enzyme | 0,78 | 3,23E-03 | 0,048 | DOWN |
| Q9CQ69 | UQCRQ | ubiquinol-cytochrome c reductase complex III subunit VII | Cytoplasm | enzyme | 1,29 | 2,48E-03 | 0,048 | UP |
| Q9Z1Z0 | USO1 | USO1 vesicle transport factor | Cytoplasm | other | 1,44 | 2,81E-03 | 0,048 | UP |
| P70398 | USP9X | ubiquitin specific peptidase 9 X-linked | Plasma Membrane | peptidase | 1,33 | 2,87E-03 | 0,048 | UP |
| E9PYH0 | VCAN | versican | Extracellular Space | other | 0,79 | 4,61E-04 | 0,048 | DOWN |
| Q5EBQ0 | VDAC3 | voltage dependent anion channel 3 | Cytoplasm | ion channel | 1,29 | 1,93E-03 | 0,048 | UP |
| A1ILG8 | VPS13C | vacuolar protein sorting 13 homolog C | Cytoplasm | other | 1,56 | 8,73E-04 | 0,048 | UP |
| Q3TJ43 | VPS35 | VPS35 retromer complex component | Cytoplasm | transporter | 1,40 | 3,34E-03 | 0,048 | UP |
| Q91XH6 | VTI1B | vesicle transport through interaction with t-SNAREs 1B | Plasma Membrane | transporter | 0,78 | 2,37E-03 | 0,048 | DOWN |
| Q8CGF6 | WDR47 | WD repeat domain 47 | Cytoplasm | other | 1,55 | 3,83E-03 | 0,049 | UP |
| Q920I9 | WDR7 | WD repeat domain 7 | Cytoplasm | other | 1,44 | 2,14E-03 | 0,048 | UP |
| Q99J09 | WDR77 | WD repeat domain 77 | Nucleus | transcription regulator | 0,76 | 1,66E-03 | 0,048 | DOWN |
