## Supplementary table 5 for "Mouse PAW: reverse-translating the FINGER multimodal lifestyle intervention enhances synaptic plasticity and cognition in adult wild type female mice"

| UniProt ID | Protein ID | Protein Name | Location | Type | Fold Change | p-value | FDR |
|---|---|---|---|---|---|---|---|
| Q6P542 | ABCF1 | ATP binding cassette subfamily F member 1 | Cytoplasm | transporter | 1,26 | 0,013 | 0,891 |
| Q9D358 | ACP1 | acid phosphatase 1 | Cytoplasm | phosphatase | 1,49 | 0,008 | 0,817 |
| E9QJT5 | ACYP1 | acylphosphatase 1 | Cytoplasm | enzyme | 1,22 | 0,005 | 0,697 |
| E9PVE8 | ADCYAP1R1 | ADCYAP receptor type I | Plasma Membrane | G-protein coupled receptor | 0,61 | 0,028 | 0,996 |
| Q80YS6 | AFAP1 | actin filament associated protein 1 | Cytoplasm | other | 1,45 | 0,004 | 0,697 |
| Q4FJL9 | AIF1 | allograft inflammatory factor 1 | Nucleus | other | 1,14 | 0,047 | 0,997 |
| Q9R0Y5 | AK1 | adenylate kinase 1 | Cytoplasm | kinase | 1,18 | 0,001 | 0,452 |
| B2RRE0 | AKAP12 | A-kinase anchoring protein 12 | Cytoplasm | transporter | 1,28 | 0,049 | 0,997 |
| G5E895 | Akr1b10 | aldo-keto reductase family 1, member B10 | Cytoplasm | enzyme | 0,86 | 0,010 | 0,821 |
| Q8CG76 | AKR7A2 | aldo-keto reductase family 7 member A2 | Cytoplasm | enzyme | 0,90 | 0,047 | 0,997 |
| Q6NXW0 | AKT3 | AKT serine/threonine kinase 3 | Cytoplasm | kinase | 0,81 | 0,019 | 0,973 |
| Q546G4 | ALB | albumin | Extracellular Space | transporter | 0,77 | 0,036 | 0,997 |
| P61967 | AP1S1 | adaptor related protein complex 1 subunit sigma 1 | Cytoplasm | transporter | 1,48 | 0,041 | 0,997 |
| Q3U0D7 | ARF6 | ADP ribosylation factor 6 | Plasma Membrane | transporter | 1,28 | 0,027 | 0,996 |
| B7ZCJ1 | ARHGAP21 | Rho GTPase activating protein 21 | Cytoplasm | other | 0,50 | 0,035 | 0,997 |
| Q9D0L7 | ARMC10 | armadillo repeat containing 10 | Cytoplasm | other | 1,18 | 0,009 | 0,821 |
| Q9CVB6 | ARPC2 | actin related protein 2/3 complex subunit 2 | Cytoplasm | other | 0,90 | 0,011 | 0,842 |
| Q9D898 | ARPC5L | actin related protein 2/3 complex subunit 5 like | Cytoplasm | other | 1,18 | 0,033 | 0,996 |
| A0A0R4J138 | ARSB | arylsulfatase B | Cytoplasm | enzyme | 1,12 | 0,043 | 0,997 |
| Q8R3P0 | ASPA | aspartoacylase | Cytoplasm | enzyme | 0,83 | 0,012 | 0,846 |
| Q3UR55 | ATP1B2 | ATPase Na+/K+ transporting subunit beta 2 | Plasma Membrane | transporter | 0,88 | 0,030 | 0,996 |
| P56383 | ATP5MC2 | ATP synthase membrane subunit c locus 2 | Cytoplasm | transporter | 1,30 | 0,024 | 0,996 |
| Q9DCX2 | ATP5PD | ATP synthase peripheral stalk subunit d | Cytoplasm | enzyme | 1,14 | 0,030 | 0,996 |
| Q3U861 | ATP6V1D | ATPase H+ transporting V1 subunit D | Cytoplasm | transporter | 1,11 | 0,039 | 0,997 |
| Q6DI68 | ATP9A | ATPase phospholipid transporting 9A (putative) | Cytoplasm | transporter | 0,76 | 0,043 | 0,997 |
| Q80U56 | AVL9 | AVL9 cell migration associated | Cytoplasm | other | 1,33 | 0,025 | 0,996 |
| Q8CI32 | BAG5 | BAG cochaperone 5 | Cytoplasm | other | 0,91 | 0,045 | 0,997 |
| Q6P3A8 | BCKDHB | branched chain keto acid dehydrogenase E1 subunit beta | Cytoplasm | enzyme | 0,77 | 0,019 | 0,973 |
| O88597 | BECN1 | beclin 1 | Cytoplasm | other | 1,15 | 0,005 | 0,697 |
| Q8K2Q7 | BROX | BRO1 domain and CAAX motif containing | Cytoplasm | other | 1,36 | 0,037 | 0,997 |
| Q91WE4 | C18orf32 | chromosome 18 open reading frame 32 | Other | other | 1,35 | 0,042 | 0,997 |
| Q8BGN2 | C4orf33 | chromosome 4 open reading frame 33 | Other | other | 1,44 | 0,048 | 0,997 |
| B2RS43 | C6orf47 | chromosome 6 open reading frame 47 | Cytoplasm | other | 1,27 | 0,032 | 0,996 |
| Q9ERQ8 | CA7 | carbonic anhydrase 7 | Cytoplasm | enzyme | 1,35 | 0,024 | 0,996 |
| Q8BGR3 | CAMK4 | calcium/calmodulin dependent protein kinase IV | Nucleus | kinase | 1,16 | 0,033 | 0,996 |
| G3X975 | CARS2 | cysteinyl-tRNA synthetase 2, mitochondrial | Cytoplasm | enzyme | 0,83 | 0,045 | 0,997 |
| P48758 | CBR1 | carbonyl reductase 1 | Cytoplasm | enzyme | 1,11 | 0,018 | 0,973 |
| Q7TQK5 | CCDC93 | coiled-coil domain containing 93 | Extracellular Space | other | 0,79 | 0,044 | 0,997 |
| P35762 | CD81 | CD81 molecule | Plasma Membrane | other | 0,74 | 0,020 | 0,973 |
| Q3UII2 | CD82 | CD82 molecule | Plasma Membrane | other | 0,73 | 0,000 | 0,285 |
| Q9JKC6 | CEND1 | cell cycle exit and neuronal differentiation 1 | Cytoplasm | other | 1,38 | 0,013 | 0,891 |
| Q8C6E0 | CFAP36 | cilia and flagella associated protein 36 | Extracellular Space | other | 1,44 | 0,010 | 0,821 |
| Q9QXG2 | CHM | CHM Rab escort protein | Cytoplasm | enzyme | 0,78 | 0,016 | 0,973 |
| Q91WS0 | CISD1 | CDGSH iron sulfur domain 1 | Cytoplasm | enzyme | 0,81 | 0,011 | 0,842 |
| Q9QYB1 | CLIC4 | chloride intracellular channel 4 | Plasma Membrane | ion channel | 1,24 | 0,049 | 0,997 |
| Q8CJ61 | CMTM4 | CKLF like MARVEL transmembrane domain containing 4 | Extracellular Space | cytokine | 1,26 | 0,039 | 0,997 |
| A0A1S6GWJ6 | CNOT2 | CCR4-NOT transcription complex subunit 2 | Nucleus | transcription regulator | 1,55 | 0,000 | 0,312 |
| Q3V231 | CNOT7 | CCR4-NOT transcription complex subunit 7 | Nucleus | transcription regulator | 1,80 | 0,047 | 0,997 |
| P16330 | CNP | 2',3'-cyclic nucleotide 3' phosphodiesterase | Cytoplasm | enzyme | 0,74 | 0,002 | 0,521 |
| Q3UQL2 | COPS3 | COP9 signalosome subunit 3 | Cytoplasm | other | 0,92 | 0,010 | 0,821 |
| Q8VBV7 | COPS8 | COP9 signalosome subunit 8 | Nucleus | other | 1,35 | 0,013 | 0,891 |
| Q8BLR2 | CPNE4 | copine 4 | Cytoplasm | other | 0,80 | 0,032 | 0,996 |
| Q561M8 | CPS1 | carbamoyl-phosphate synthase 1 | Cytoplasm | enzyme | 1,32 | 0,040 | 0,997 |
| Q9DCT8 | Crip2 | cysteine rich protein 2 | Plasma Membrane | other | 0,83 | 0,049 | 0,997 |
| Q921W4 | CRYZL1 | crystallin zeta like 1 | Cytoplasm | enzyme | 0,81 | 0,008 | 0,817 |
| Q9JMK2 | CSNK1E | casein kinase 1 epsilon | Cytoplasm | kinase | 1,32 | 0,030 | 0,996 |
| Q4FJX4 | CSRP1 | cysteine and glycine rich protein 1 | Nucleus | other | 0,84 | 0,008 | 0,817 |
| P21460 | CST3 | cystatin C | Extracellular Space | other | 1,73 | 0,006 | 0,763 |
| A0A0A6YX71 | DCLK2 | doublecortin like kinase 2 | Cytoplasm | kinase | 1,47 | 0,022 | 0,996 |
| Q9DAR7 | DCPS | decapping enzyme, scavenger | Nucleus | enzyme | 0,88 | 0,000 | 0,285 |
| A2ADY9 | DDI2 | DNA damage inducible 1 homolog 2 | Plasma Membrane | transporter | 1,18 | 0,017 | 0,973 |
| A0A1S6GWH2 | DDX39B | DExD-box helicase 39B | Nucleus | enzyme | 0,90 | 0,038 | 0,997 |
| Q9CQT7 | DESI1 | desumoylating isopeptidase 1 | Cytoplasm | peptidase | 1,34 | 0,032 | 0,996 |
| Q99LB2 | DHRS4 | dehydrogenase/reductase 4 | Cytoplasm | enzyme | 0,87 | 0,021 | 0,995 |
| Q9D2G2 | DLST | dihydrolipoamide S-succinyltransferase | Cytoplasm | enzyme | 1,13 | 0,022 | 0,996 |
| O08553 | DPYSL2 | dihydropyrimidinase like 2 | Cytoplasm | enzyme | 1,16 | 0,016 | 0,973 |
| Q9R0P5 | DSTN | destrin, actin depolymerizing factor | Cytoplasm | other | 0,88 | 0,042 | 0,997 |
| Q9DD18 | DTD1 | D-aminoacyl-tRNA deacylase 1 | Cytoplasm | enzyme | 1,10 | 0,028 | 0,996 |
| A0A0R4J1E2 | EEF1D | eukaryotic translation elongation factor 1 delta | Cytoplasm | translation regulator | 1,16 | 0,039 | 0,997 |
| Q544L9 | EFNB1 | ephrin B1 | Plasma Membrane | other | 1,13 | 0,015 | 0,962 |
| Q69ZW3 | EHBP1 | EH domain binding protein 1 | Cytoplasm | other | 1,23 | 0,031 | 0,996 |
| Q9DBZ5 | EIF3K | eukaryotic translation initiation factor 3 subunit K | Cytoplasm | translation regulator | 1,18 | 0,002 | 0,521 |
| Q8BK75 | ELP6 | elongator acetyltransferase complex subunit 6 | Other | other | 1,31 | 0,006 | 0,763 |
| Q69ZY2 | ENDOD1 | endonuclease domain containing 1 | Extracellular Space | enzyme | 1,21 | 0,044 | 0,997 |
| Q60902 | EPS15L1 | epidermal growth factor receptor pathway substrate 15 like 1 | Plasma Membrane | other | 1,11 | 0,032 | 0,996 |
| Q69ZP5 | ERGIC1 | endoplasmic reticulum-golgi intermediate compartment 1 | Cytoplasm | other | 1,10 | 0,047 | 0,997 |
| Q69ZD1 | EXOC4 | exocyst complex component 4 | Cytoplasm | transporter | 1,37 | 0,046 | 0,997 |
| P26040 | EZR | ezrin | Plasma Membrane | other | 1,10 | 0,038 | 0,997 |
| A0A0R4J094 | FAHD2B | fumarylacetoacetate hydrolase domain containing 2B | Cytoplasm | other | 1,70 | 0,047 | 0,997 |
| Q8BWU3 | FAM131A | family with sequence similarity 131 member A | Other | transporter | 0,00 | 0,020 | 0,973 |
| Q80VD1 | FAM98B | family with sequence similarity 98 member B | Nucleus | enzyme | 0,81 | 0,008 | 0,817 |
| A9E3L2 | FKBP1B | FKBP prolyl isomerase 1B | Cytoplasm | enzyme | 1,40 | 0,029 | 0,996 |
| Q3UBU9 | FKBP3 | FKBP prolyl isomerase 3 | Nucleus | enzyme | 1,13 | 0,020 | 0,973 |
| Q9D6K8 | FUNDC2 | FUN14 domain containing 2 | Cytoplasm | other | 1,24 | 0,033 | 0,996 |
| Q8BFQ8 | GATD1 | glutamine amidotransferase class 1 domain containing 1 | Cytoplasm | other | 1,12 | 0,044 | 0,997 |
| Q9D7M1 | GID8 | GID complex subunit 8 homolog | Nucleus | other | 1,18 | 0,027 | 0,996 |
| Q9JL35 | Gm15807/Hmgn5 | high-mobility group nucleosome binding domain 5 | Nucleus | other | 1,28 | 0,010 | 0,821 |
| B2M0S2 | Gm45927 | predicted gene, 45927 | Other | other | 1,30 | 0,031 | 0,996 |
| P08752 | GNAI2 | G protein subunit alpha i2 | Plasma Membrane | enzyme | 0,78 | 0,027 | 0,996 |
| E9QP99 | GOLGA3 | golgin A3 | Cytoplasm | transporter | 1,28 | 0,007 | 0,765 |
| Q542P2 | GPM6A | glycoprotein M6A | Plasma Membrane | ion channel | 0,67 | 0,002 | 0,523 |

| | | | | | | | |
|---|---|---|---|---|---|---|---|
| Q3UNH4 | GPRIN1 | G protein regulated inducer of neurite outgrowth 1 | Plasma Membrane | other | 1,15 | 0,002 | 0,521 |
| Q76LV0 | GPX4 | glutathione peroxidase 4 | Cytoplasm | enzyme | 0,91 | 0,011 | 0,842 |
| P43275 | H1f1 | H1.1 linker histone, cluster member | Nucleus | other | 1,25 | 0,026 | 0,996 |
| Q9QUP5 | HAPLN1 | hyaluronan and proteoglycan link protein 1 | Extracellular Space | other | 0,87 | 0,024 | 0,996 |
| Q8R1F6 | HID1 | HID1 domain containing | Plasma Membrane | other | 3,08 | 0,023 | 0,996 |
| O54879 | Hmgb3 | high mobility group box 3 | Nucleus | other | 1,94 | 0,018 | 0,973 |
| P18608 | HMGN1 | high mobility group nucleosome binding domain 1 | Nucleus | transcription regulator | 1,34 | 0,025 | 0,996 |
| Q3U6W4 | HMOX2 | heme oxygenase 2 | Cytoplasm | enzyme | 1,11 | 0,019 | 0,973 |
| O88569 | HNRNPA2B1 | heterogeneous nuclear ribonucleoprotein A2/B1 | Nucleus | other | 0,88 | 0,043 | 0,997 |
| Q91X72 | HPX | hemopexin | Extracellular Space | transporter | 0,67 | 0,024 | 0,996 |
| P17879 | Hspa1b | heat shock protein 1B | Cytoplasm | enzyme | 0,93 | 0,023 | 0,996 |
| P17156 | HSPA2 | heat shock protein family A (Hsp70) member 2 | Cytoplasm | other | 0,91 | 0,001 | 0,479 |
| Q4KL76 | HSPE1 | heat shock protein family E (Hsp10) member 1 | Cytoplasm | enzyme | 1,18 | 0,022 | 0,996 |
| P59644 | INPP5J | inositol polyphosphate-5-phosphatase J | Plasma Membrane | phosphatase | 1,41 | 0,009 | 0,821 |
| Q9JHU9 | ISYNA1 | inositol-3-phosphate synthase 1 | Cytoplasm | enzyme | 0,85 | 0,040 | 0,997 |
| Q9DBN4 | KIAA1191 | KIAA1191 | Cytoplasm | other | 1,15 | 0,025 | 0,996 |
| A0A2R8VKG2 | KIAA1549L | KIAA1549 like | Cytoplasm | other | 1,17 | 0,036 | 0,997 |
| Q61792 | LASP1 | LIM and SH3 protein 1 | Cytoplasm | transporter | 0,84 | 0,033 | 0,996 |
| Q3U944 | LMAN1 | lectin, mannose binding 1 | Cytoplasm | other | 1,13 | 0,013 | 0,891 |
| P97823 | LYPLA1 | lysophospholipase 1 | Cytoplasm | enzyme | 1,16 | 0,035 | 0,997 |
| Q9CXI5 | MANF | mesencephalic astrocyte derived neurotrophic factor | Extracellular Space | other | 0,67 | 0,005 | 0,697 |
| Q8R3Q6 | MIX23 | mitochondrial matrix import factor 23 | Cytoplasm | other | 0,86 | 0,046 | 0,997 |
| Q80YU5 | MOG | myelin oligodendrocyte glycoprotein | Extracellular Space | other | 0,86 | 0,041 | 0,997 |
| Q80X85 | MRPS7 | mitochondrial ribosomal protein S7 | Cytoplasm | other | 1,54 | 0,001 | 0,452 |
| Q61474 | MSI1 | musashi RNA binding protein 1 | Cytoplasm | other | 0,54 | 0,000 | 0,312 |
| Q9CZH7 | Mxra7 | matrix-remodelling associated 7 | Extracellular Space | other | 1,18 | 0,033 | 0,996 |
| B0R0C1 | MYT1 | myelin transcription factor 1 | Nucleus | transcription regulator | 0,71 | 0,019 | 0,973 |
| Q3UK27 | NAE1 | NEDD8 activating enzyme E1 subunit 1 | Cytoplasm | enzyme | 1,36 | 0,008 | 0,817 |
| Q9DB05 | NAPA | NSF attachment protein alpha | Cytoplasm | transporter | 0,89 | 0,035 | 0,997 |
| Q9ERS2 | NDUFA13 | NADH:ubiquinone oxidoreductase subunit A13 | Cytoplasm | enzyme | 1,13 | 0,027 | 0,996 |
| Q9CQ45 | NENF | neudesin neurotrophic factor | Extracellular Space | growth factor | 1,28 | 0,003 | 0,523 |
| A0A0N4SUH8 | NFU1 | NFU1 iron-sulfur cluster scaffold | Cytoplasm | other | 1,28 | 0,020 | 0,973 |
| P30415 | NKTR | natural killer cell triggering receptor | Plasma Membrane | enzyme | 0,79 | 0,037 | 0,997 |
| A0A286YDV7 | Nolc1 | nucleolar and coiled-body phosphoprotein 1 | Nucleus | other | 1,14 | 0,010 | 0,821 |
| D3Z4E2 | NRN1 | neuritin 1 | Cytoplasm | other | 0,78 | 0,003 | 0,625 |
| Q6PIP5 | NUDCD1 | NudC domain containing 1 | Nucleus | other | 0,87 | 0,039 | 0,997 |
| O08919 | NUMBL | NUMB like endocytic adaptor protein | Cytoplasm | other | 0,92 | 0,008 | 0,817 |
| Q7M761 | OVCH2 | ovochymase 2 | Extracellular Space | enzyme | 1,19 | 0,038 | 0,997 |
| Q4KMM3 | OXR1 | oxidation resistance 1 | Cytoplasm | enzyme | 1,10 | 0,021 | 0,995 |
| Q3TJL8 | PDIA6 | protein disulfide isomerase family A member 6 | Cytoplasm | enzyme | 1,16 | 0,033 | 0,996 |
| Q3TCN2 | PLBD2 | phospholipase B domain containing 2 | Extracellular Space | other | 1,12 | 0,028 | 0,996 |
| Q3UYM8 | PLP1 | proteolipid protein 1 | Plasma Membrane | other | 0,69 | 0,006 | 0,763 |
| Q3TTM6 | PMPCA | peptidase, mitochondrial processing subunit alpha | Cytoplasm | peptidase | 1,21 | 0,014 | 0,922 |
| Q6R891 | PPP1R9B | protein phosphatase 1 regulatory subunit 9B | Cytoplasm | enzyme | 1,08 | 0,037 | 0,997 |
| G3UWS4 | PPP2R1B | protein phosphatase 2 scaffold subunit Abeta | Plasma Membrane | phosphatase | 0,70 | 0,021 | 0,996 |
| Q8C6A3 | PREP | prolyl endopeptidase | Cytoplasm | peptidase | 1,11 | 0,004 | 0,625 |
| Q3TYK4 | PRKAR1A | protein kinase cAMP-dependent type I regulatory subunit alpha | Cytoplasm | kinase | 0,86 | 0,042 | 0,997 |
| Q6PAK3 | PRMT8 | protein arginine methyltransferase 8 | Nucleus | enzyme | 0,85 | 0,038 | 0,997 |
| E9PUL5 | PRRT2 | proline rich transmembrane protein 2 | Plasma Membrane | other | 1,27 | 0,004 | 0,643 |
| J3QPG5 | PSAP | prosaposin | Extracellular Space | other | 0,71 | 0,001 | 0,456 |
| Q60692 | PSMB6 | proteasome 20S subunit beta 6 | Nucleus | peptidase | 1,14 | 0,025 | 0,996 |
| A0A1S6GWH1 | PSMC5 | proteasome 26S subunit, ATPase 5 | Nucleus | transcription regulator | 0,90 | 0,000 | 0,312 |
| P97371 | PSME1 | proteasome activator subunit 1 | Cytoplasm | other | 1,21 | 0,001 | 0,452 |
| A0A068BFR3 | RAB11B | RAB11B, member RAS oncogene family | Cytoplasm | enzyme | 1,13 | 0,030 | 0,996 |
| A0A494B945 | RAB1B | RAB1B, member RAS oncogene family | Cytoplasm | other | 1,28 | 0,014 | 0,922 |
| Q99P58 | RAB27B | RAB27B, member RAS oncogene family | Cytoplasm | enzyme | 1,17 | 0,037 | 0,997 |
| A2A7Z6 | RAB3B | RAB3B, member RAS oncogene family | Cytoplasm | enzyme | 0,78 | 0,017 | 0,973 |
| Q91ZR1 | RAB4B | RAB4B, member RAS oncogene family | Plasma Membrane | enzyme | 1,10 | 0,027 | 0,996 |
| Q3KNA5 | RAB9B | RAB9B, member RAS oncogene family | Plasma Membrane | enzyme | 1,24 | 0,027 | 0,996 |
| Q3TLP8 | RAC1 | Rac family small GTPase 1 | Plasma Membrane | enzyme | 0,80 | 0,002 | 0,521 |
| A0A3B2WCL5 | RANBP3 | RAN binding protein 3 | Nucleus | other | 1,16 | 0,028 | 0,996 |
| Q8BG13 | RBM3 | RNA binding motif protein 3 | Cytoplasm | other | 1,29 | 0,010 | 0,821 |
| Q91YE7 | RBM5 | RNA binding motif protein 5 | Nucleus | other | 1,43 | 0,043 | 0,997 |
| Q3U111 | RDX | radixin | Cytoplasm | other | 1,12 | 0,032 | 0,996 |
| B9EI38 | REPS2 | RALBP1 associated Eps domain containing 2 | Cytoplasm | other | 1,66 | 0,047 | 0,997 |
| P48379 | RFX2 | regulatory factor X2 | Nucleus | transcription regulator | 0,51 | 0,002 | 0,521 |
| Q9JJC6 | RILPL1 | Rab interacting lysosomal protein like 1 | Cytoplasm | other | 1,18 | 0,030 | 0,996 |
| Q9QYK7 | RNF11 | ring finger protein 11 | Cytoplasm | enzyme | 0,79 | 0,006 | 0,763 |
| A0A1B0GSG5 | RNH1 | ribonuclease/angiogenin inhibitor 1 | Cytoplasm | other | 0,86 | 0,013 | 0,891 |
| P47964 | Rpl36 | ribosomal protein L36 | Nucleus | other | 0,74 | 0,001 | 0,479 |
| P99027 | RPLP2 | ribosomal protein lateral stalk subunit P2 | Cytoplasm | other | 1,14 | 0,008 | 0,817 |
| A2ATS6 | RTN4RL2 | reticulon 4 receptor like 2 | Plasma Membrane | other | 1,08 | 0,034 | 0,996 |
| P70122 | SBDS | SBDS ribosome maturation factor | Nucleus | other | 0,87 | 0,046 | 0,997 |
| Q3U276 | SDHAF1 | succinate dehydrogenase complex assembly factor 1 | Cytoplasm | other | 0,00 | 0,023 | 0,996 |
| Q8R238 | SDSL | serine dehydratase like | Cytoplasm | enzyme | 1,18 | 0,011 | 0,846 |
| P63300 | SELENOW | selenoprotein W | Cytoplasm | enzyme | 0,22 | 0,003 | 0,523 |
| Q6P6M7 | SEPSECS | Sep (O-phosphoserine) tRNA:Sec (selenocysteine) tRNA synthase | Cytoplasm | enzyme | 0,77 | 0,042 | 0,997 |
| Q3U213 | SERAC1 | serine active site containing 1 | Extracellular Space | other | 0,24 | 0,005 | 0,697 |
| O08797 | SERPINB9 | serpin family B member 9 | Cytoplasm | other | 1,19 | 0,042 | 0,997 |
| Q91WC0 | SETD3 | SET domain containing 3, actin N3(tau)-histidine methyltransferase | Nucleus | enzyme | 1,17 | 0,032 | 0,996 |
| Q8K2Q9 | SHTN1 | shootin 1 | Plasma Membrane | other | 1,18 | 0,030 | 0,996 |
| O35874 | SLC1A4 | solute carrier family 1 member 4 | Plasma Membrane | transporter | 1,17 | 0,023 | 0,996 |
| Q5U680 | SLC25A26 | solute carrier family 25 member 26 | Cytoplasm | transporter | 1,78 | 0,014 | 0,891 |
| Q14AR0 | SLIRP | SRA stem-loop interacting RNA binding protein | Cytoplasm | other | 1,28 | 0,045 | 0,997 |
| E9PYU8 | SMURF1 | SMAD specific E3 ubiquitin protein ligase 1 | Cytoplasm | enzyme | 0,25 | 0,025 | 0,996 |
| P08228 | SOD1 | superoxide dismutase 1 | Cytoplasm | enzyme | 0,73 | 0,000 | 0,312 |
| P49962 | SRP9 | signal recognition particle 9 | Cytoplasm | other | 0,77 | 0,040 | 0,997 |
| Q9EPQ7 | STARD5 | StAR related lipid transfer domain containing 5 | Cytoplasm | transporter | 1,54 | 0,048 | 0,997 |
| Q9CZ46 | STMN2 | stathmin 2 | Plasma Membrane | other | 1,18 | 0,032 | 0,996 |
| Q9D0I4 | STX17 | syntaxin 17 | Plasma Membrane | transporter | 1,13 | 0,043 | 0,997 |
| O09117 | SYPL1 | synaptophysin like 1 | Plasma Membrane | transporter | 0,70 | 0,002 | 0,521 |
| Q921I1 | TF | transferrin | Extracellular Space | transporter | 0,73 | 0,005 | 0,697 |
| P62075 | TIMM13 | translocase of inner mitochondrial membrane 13 | Cytoplasm | transporter | 1,24 | 0,042 | 0,997 |

| | | | | | | | |
|---|---|---|---|---|---|---|---|
| A2RS53 | TMED2 | transmembrane p24 trafficking protein 2 | Cytoplasm | transporter | 1,76 | 0,012 | 0,846 |
| Q9QZ06 | TOLLIP | toll interacting protein | Cytoplasm | other | 1,08 | 0,046 | 0,997 |
| A0A1S6GWJ9 | TSFM | Ts translation elongation factor, mitochondrial | Cytoplasm | translation regulator | 0,88 | 0,018 | 0,973 |
| Q80WR1 | TSPAN18 | tetraspanin 18 | Other | other | 2,12 | 0,010 | 0,821 |
| Q922J6 | TSPAN2 | tetraspanin 2 | Extracellular Space | other | 0,77 | 0,016 | 0,973 |
| Q9ERD7 | TUBB3 | tubulin beta 3 class III | Cytoplasm | other | 0,85 | 0,003 | 0,625 |
| P26369 | U2AF2 | U2 small nuclear RNA auxiliary factor 2 | Nucleus | other | 1,20 | 0,002 | 0,521 |
| Q0KL01 | UBXN2B | UBX domain protein 2B | Cytoplasm | other | 1,37 | 0,035 | 0,997 |
| Q99PL6 | UBXN6 | UBX domain protein 6 | Cytoplasm | other | 0,91 | 0,009 | 0,821 |
| Q9CR68 | UQCRFS1 | ubiquinol-cytochrome c reductase, Rieske iron-sulfur polypeptide 1 | Cytoplasm | enzyme | 1,12 | 0,017 | 0,973 |
| Q9D9M2 | USP12 | ubiquitin specific peptidase 12 | Cytoplasm | peptidase | 0,73 | 0,032 | 0,996 |
| E9PV45 | USP24 | ubiquitin specific peptidase 24 | Nucleus | peptidase | 0,00 | 0,042 | 0,997 |
| E9PYH0 | VCAN | versican | Extracellular Space | other | 0,86 | 0,026 | 0,996 |
| Q80T83 | VCPIP1 | valosin containing protein interacting protein 1 | Cytoplasm | peptidase | 1,28 | 0,050 | 0,997 |
| Q9CRC0 | VKORC1 | vitamin K epoxide reductase complex subunit 1 | Cytoplasm | enzyme | 0,55 | 0,006 | 0,763 |
| Q6TEK5 | VKORC1L1 | vitamin K epoxide reductase complex subunit 1 like 1 | Cytoplasm | enzyme | 0,73 | 0,032 | 0,996 |
| Q9R0D8 | WDR54 | WD repeat domain 54 | Other | other | 1,44 | 0,030 | 0,996 |
| Q5GH66 | XKR5 | XK related 5 | Plasma Membrane | other | 1,41 | 0,049 | 0,997 |
| Q9CQW1 | YKT6 | YKT6 v-SNARE homolog | Cytoplasm | enzyme | 1,18 | 0,035 | 0,997 |
| Q0VAV0 | Zfp947/Zfp995 | zinc finger protein 947 | Other | other | 1,31 | 0,035 | 0,997 |
| H3BLD9 | ZNF428 | zinc finger protein 428 | Other | other | 1,22 | 0,043 | 0,997 |
| Q9CQU5 | ZWINT | ZW10 interacting kinetochore protein | Nucleus | other | 1,19 | 0,033 | 0,996 |
