## Supplementary table 6 for "Mouse PAW: reverse-translating the FINGER multimodal lifestyle intervention enhances synaptic plasticity and cognition in adult wild type female mice"

SUPPLEMENTARY TABLE 6.
© 2000-2024 QIAGEN. All rights reserved.

| Ingenuity Canonical Pathways | -log10(p-value) | Ratio | Z-score | Molecules |
|---|---|---|---|---|
| Glutamate binding, activation of AMPA receptors and synaptic plasticity | 1,85×10^01 | 4,38×10^−01 | 3,742 | AP2A1,AP2A2,Ap2b1,AP2M1,AP2S1,CACNG2,CAMK2A,CAMK2D,CAMK2G,DLG4,GRIA1,GRIA2,GRIA3,PRKCG |
| Synaptogenesis Signaling Pathway | 1,79×10^01 | 9,52×10^−02 | 4,271 | ACTR3,ADCY9,AP2A1,AP2A2,Ap2b1,AP2M1,AP2S1,CAMK2A,CAMK2D,CAMK2G,CLASP2,CRK,DLG4,FARP1,GRIA1,GRIA2,GRIA3,GRIN1,( |
| SRP-dependent cotranslational protein targeting to membrane | 1,72×10^01 | 1,74×10^−01 | 3,578 | DDOST,RPL12,RPL13,RPL14,RPL15,RPL18A,RPL24,RPL26,RPL27,RPL3,RPL35A,RPL6,RPL7A,RPL8,RPS11,RPS16,RPS3,RPS5,RPS9,SSR1 |
| Eukaryotic Translation Termination | 1,63×10^01 | 1,91×10^−01 | 3,771 | RPL12,RPL13,RPL14,RPL15,RPL18A,RPL24,RPL26,RPL27,RPL3,RPL35A,RPL6,RPL7A,RPL8,RPS11,RPS16,RPS3,RPS5,RPS9 |
| Eukaryotic Translation Elongation | 1,63×10^01 | 1,89×10^−01 | 3,771 | RPL12,RPL13,RPL14,RPL15,RPL18A,RPL24,RPL26,RPL27,RPL3,RPL35A,RPL6,RPL7A,RPL8,RPS11,RPS16,RPS3,RPS5,RPS9 |
| Response of EIF2AK4 (GCN2) to amino acid deficiency | 1,56×10^01 | 1,75×10^−01 | 3,771 | RPL12,RPL13,RPL14,RPL15,RPL18A,RPL24,RPL26,RPL27,RPL3,RPL35A,RPL6,RPL7A,RPL8,RPS11,RPS16,RPS3,RPS5,RPS9 |
| Eukaryotic Translation Initiation | 1,55×10^01 | 1,56×10^−01 | 3,900 | EIF4A2,RPL12,RPL13,RPL14,RPL15,RPL18A,RPL24,RPL26,RPL27,RPL3,RPL35A,RPL6,RPL7A,RPL8,RPS11,RPS16,RPS3,RPS5,RPS9 |
| Selenoamino acid metabolism | 1,53×10^01 | 1,68×10^−01 | 3,771 | RPL12,RPL13,RPL14,RPL15,RPL18A,RPL24,RPL26,RPL27,RPL3,RPL35A,RPL6,RPL7A,RPL8,RPS11,RPS16,RPS3,RPS5,RPS9 |
| Nonsense-Mediated Decay (NMD) | 1,46×10^01 | 1,54×10^−01 | 3,771 | RPL12,RPL13,RPL14,RPL15,RPL18A,RPL24,RPL26,RPL27,RPL3,RPL35A,RPL6,RPL7A,RPL8,RPS11,RPS16,RPS3,RPS5,RPS9 |
| Glutaminergic Receptor Signaling Pathway (Enhanced) | 1,37×10^01 | 8×10^−02 | 2,746 | ADCY9,AKT1S1,CACNG2,CAMK2A,CAMK2D,CAMK2G,DGKH,DGKZ,GABBR1,GABRA1,GABRA5,GABRG2,GRIA1,GRIA2,GRIA3,GRIN1,GR |
| EIF2 Signaling | 1,33×10^01 | 9,57×10^−02 | 3,207 | EIF4A2,RALA,RPL12,RPL13,RPL14,RPL15,RPL18A,RPL24,RPL26,RPL27,RPL3,Rpl34 (includes others),RPL35A,Rpl36,RPL6,RPL7A,RPL8,R |
| EPH-Ephrin signaling | 1,25×10^01 | 1,6×10^−01 | 3,873 | ACTR3,AP2A1,AP2A2,Ap2b1,AP2M1,AP2S1,DNM1,GIT1,GRIN1,GRIN2B,KALRN,MYH10,MYH14,MYH9,ROCK2 |
| Calcium Signaling | 1,17×10^01 | 9,09×10^−02 | 3,873 | ATP2A2,ATP2B1,ATP2B2,CACNG2,CALR,CAMK2A,CAMK2D,CAMK2G,GRIA1,GRIA2,GRIA3,GRIN1,GRIN2B,ITPR1,MYH10,MYH14,MYH9,M |
| Major pathway of rRNA processing in the nucleolus and cytosol | 1,1×10^01 | 9,68×10^−02 | 3,771 | RPL12,RPL13,RPL14,RPL15,RPL18A,RPL24,RPL26,RPL27,RPL3,RPL35A,RPL6,RPL7A,RPL8,RPS11,RPS16,RPS3,RPS5,RPS9 |
| GABA Receptor Signaling | 1,03×10^01 | 1,14×10^−01 | NA | ADCY9,AP2A1,AP2A2,Ap2b1,AP2M1,AP2S1,CACNG2,DNM1,GABBR1,GABRA1,GABRA5,GABRG2,ITPR1,SLC32A1,SLC6A11 |
| Virus Entry via Endocytic Pathways | 9,9×10^00 | 1,19×10^−01 | 3,207 | AP1B1,AP1G1,AP1M1,AP2A1,AP2A2,Ap2b1,AP2M1,AP2S1,AP3B2,AP3D1,DNM1,PRKCE,PRKCG,RALA |
| Neuropathic Pain Signaling in Dorsal Horn Neurons | 9,68×10^00 | 1,29×10^−01 | 3,606 | CAMK2A,CAMK2D,CAMK2G,GRIA1,GRIA2,GRIA3,GRIN1,GRIN2B,GRM5,ITPR1,PRKACB,PRKCE,PRKCG |
| Neurovascular Coupling Signaling Pathway | 9,45×10^00 | 7,76×10^−02 | 2,357 | ADORA1,APP,CACNG2,GABBR1,GABRA1,GABRA5,GABRG2,GRIA1,GRIA2,GRIA3,GRIN1,GRIN2B,GRM5,ITPR1,PRKACB,ROCK2,RYR2,SLC1A2 |
| Synaptic Long Term Potentiation | 9,24×10^00 | 1,06×10^−01 | 3,207 | CAMK2A,CAMK2D,CAMK2G,GRIA1,GRIA2,GRIA3,GRIN1,GRIN2B,GRM5,ITPR1,PRKACB,PRKCE,PRKCG,RALA |
| DAG and IP3 signaling | 9,01×10^00 | 2,2×10^−01 | 2,828 | ADCY9,CAMK2A,CAMK2D,CAMK2G,ITPR1,NBEA,PRKACB,PRKCE,PRKCG |
| Opioid Signaling Pathway | 8,96×10^00 | 6,79×10^−02 | 3,153 | ADCY9,AP2A1,AP2A2,Ap2b1,AP2M1,AP2S1,CACNG2,CAMK2A,CAMK2D,CAMK2G,GRIN1,GRIN2B,ITPR1,PRKACB,PRKCE,PRKCG,RALA, |
| GABAergic Receptor Signaling Pathway (Enhanced) | 8,94×10^00 | 1,01×10^−01 | 0,000 | ADCY9,CACNG2,GABBR1,GABRA1,GABRA5,GABRG2,GRIN1,GRIN2B,ITPR1,PRKACB,SLC12A2,SLC12A5,SLC32A1,SLC6A11 |
| Dilated Cardiomyopathy Signaling Pathway | 8,51×10^00 | 9,33×10^−02 | 0,577 | ADCY9,ATP2A2,CACNG2,CAMK2A,CAMK2D,CAMK2G,ITPR1,MYH10,MYH14,MYH9,MYO18A,PRKACB,PRKCE,RYR2 |
| Clathrin-mediated Endocytosis Signaling | 8,41×10^00 | 7,69×10^−02 | NA | ACTR3,AMPH,AP1B1,AP1G1,AP1M1,AP2A1,AP2A2,Ap2b1,AP2M1,AP2S1,AP3B2,AP3D1,DNM1,HIP1R,SYNJ1,USP9X |
| Activation of NMDA receptors and postsynaptic events | 8,36×10^00 | 1,31×10^−01 | 2,236 | CAMK2A,CAMK2D,CAMK2G,DLG4,GIT1,GRIA1,GRIA2,GRIA3,GRIN1,GRIN2B,PRKACB |
| Glutamate Receptor Signaling | 8,28×10^00 | 1,52×10^−01 | 2,449 | DLG4,GRIA1,GRIA2,GRIA3,GRIN1,GRIN2B,GRM5,SLC17A7,SLC1A2,SLC38A1 |
| Neurotransmitter release cycle | 7,73×10^00 | 2×10^−01 | 2,121 | LIN7C,MAOA,PPFIA2,PPFIA3,SLC17A7,SLC1A2,SYN1,SYN3 |
| Fatty acyl-CoA biosynthesis | 7,67×10^00 | 2,59×10^−01 | 2,646 | ACACA,ACLY,ACSBG1,FASN,HACD2,HACD3,TECR |
| Circadian Rhythm Signaling | 7,64×10^00 | 6,34×10^−02 | NA | ADCY9,AKT1S1,CACNG2,CAMK2A,CAMK2D,CAMK2G,GRIA1,GRIA2,GRIA3,GRIN1,GRIN2B,ITPR1,PRKACB,PRKCE,PRKCG,RALA,RYR2 |
| Parkinson's Signaling Pathway | 7,53×10^00 | 5,86×10^−02 | 1,414 | ATP6V0A1,ATP6V1D,ATP6V1H,CACNG2,CAMK2A,CAMK2D,CAMK2G,GRIA1,GRIA2,GRIA3,GRIN1,GRIN2B,ITPR1,MAOA,MT-ND4,MT-ND5 |
| L1CAM interactions | 7,51×10^00 | 9,6×10^−02 | 2,309 | Ank2,AP2A1,AP2A2,AP2M1,AP2S1,DLG4,DNM1,NCAN,NFASC,SCN2A,SPTBN1,SPTBN2 |
| Assembly and cell surface presentation of NMDA receptors | 7,47×10^00 | 1,86×10^−01 | 2,121 | CAMK2A,CAMK2D,CAMK2G,DLG4,GRIN1,GRIN2B,LIN7C,NBEA |
| SNARE Signaling Pathway | 7,1×10^00 | 8,82×10^−02 | 1,732 | ADCY9,CAMK2A,CAMK2D,CAMK2G,MYH10,MYH14,MYH9,MYO18A,PRKACB,SYN1,SYN3,VTI1B |
| Mitochondrial Dysfunction | 6,79×10^00 | 5,23×10^−02 | −0,471 | ACO2,APP,ATP5PD,CACNG2,CAMK2A,CAMK2D,CAMK2G,HTT,ITPR1,MAOA,MFN2,MT-ND4,MT-ND5,NDUFA9,PDHX,PRKACB,UQCRQ,VDAC3 |
| Neutrophil degranulation | 6,58×10^00 | 4,41×10^−02 | 2,400 | ACLY,AP1M1,AP2A2,ATP6V0A1,ATP6V1D,CAND1,COPB1,CYFIP1,DDOST,DYNC1H1,IQGAP2,NFASC,PAFAH1B2,PFKL,PGRMC1,PSAP,P |
| Pancreatic Secretion Signaling Pathway | 6,54×10^00 | 6,05×10^−02 | 2,840 | ADCY9,AKT1S1,ARHGEF2,ATP2A2,ATP2B1,ATP2B2,ITPR1,PRKACB,PRKCE,PRKCG,ROCK2,RYR2,SLC12A2,SLC4A4,VTI1B |
| MHC class II antigen presentation | 6,51×10^00 | 8,73×10^−02 | 3,317 | AP1B1,AP1G1,AP1M1,AP2A1,AP2A2,AP2M1,AP2S1,DCTN1,DNM1,DYNC1H1,SPTBN2 |
| Transport of inorganic cations/anions and amino acids/oligopeptides | 6,36×10^00 | 9,62×10^−02 | 3,162 | SLC12A2,SLC12A5,SLC17A7,SLC1A2,SLC24A2,SLC32A1,SLC38A1,SLC4A10,SLC4A4,SLC7A8 |
| Reelin Signaling in Neurons | 6,11×10^00 | 7,97×10^−02 | 1,667 | ACTR3,APP,ARHGEF2,CAMK2A,CAMK2D,CAMK2G,CRK,GRIN1,GRIN2B,PAFAH1B2,PAFAH1B3 |
| Insulin Secretion Signaling Pathway | 5,97×10^00 | 5,45×10^−02 | 2,138 | ADCY9,CACNG2,CAMK2A,CAMK2D,CAMK2G,EIF4A2,ITPR1,PDHX,PRKACB,PRKCE,PRKCG,RYR2,SLC2A3,SSR1,VTI1B |
| Netrin Signaling | 5,97×10^00 | 6,94×10^−02 | 2,887 | ACTR3,CACNG2,CAMK2A,CAMK2D,CAMK2G,GRIA1,ITPR1,PRKACB,PRKCE,PRKCG,RYR2,VTI1B |
| Sleep NREM Signaling Pathway | 5,96×10^00 | 8,7×10^−02 | 1,897 | ADCY9,ADORA1,CAMK2A,CAMK2D,CAMK2G,GABRA1,GABRA5,GABRG2,PRKACB,RALA |
| nNOS Signaling in Neurons | 5,91×10^00 | 1,49×10^−01 | 2,000 | CAMK2A,DLG4,GRIN1,GRIN2B,PFKM,PRKCE,PRKCG |
| Actin Cytoskeleton Signaling | 5,87×10^00 | 5,74×10^−02 | 1,265 | ACTR3,CRK,CYFIP1,CYFIP2,GIT1,IQGAP2,MYH10,MYH14,MYH9,MYO18A,NCKAP1,RALA,ROCK2,TLN2 |
| Protein Sorting Signaling Pathway | 5,84×10^00 | 6,74×10^−02 | 2,887 | ADCY9,AP1B1,AP1G1,AP1M1,AP3B2,AP3D1,COPA,COPB1,COPG1,PLPP3,PRKACB,VPS35 |
| Endocannabinoid Neuronal Synapse Pathway | 5,78×10^00 | 7,38×10^−02 | 1,897 | ADCY9,CACNG2,FAAH,GRIA1,GRIA2,GRIA3,GRIN1,GRIN2B,GRM5,ITPR1,PRKACB |
| Ion channel transport | 5,71×10^00 | 6,56×10^−02 | 2,309 | ANO3,ATP2A2,ATP2B1,ATP2B2,ATP6V0A1,ATP6V1D,ATP6V1H,CAMK2A,CAMK2D,CAMK2G,NEDD4L,RYR2 |
| RAF/MAP kinase cascade | 5,51×10^00 | 5,34×10^−02 | 3,742 | CAMK2A,CAMK2D,CAMK2G,DLG4,GRIN1,GRIN2B,PEA15,PSMD1,PSMD6,RAPGEF2,RASGRP1,SPTBN1,SPTBN2,SYNGAP1 |
| Cardiac conduction | 5,47×10^00 | 7,69×10^−02 | 3,000 | ATP2A2,ATP2B1,ATP2B2,CAMK2A,CAMK2D,CAMK2G,ITPR1,KCND2,RYR2,SCN2A |
| Clathrin-mediated endocytosis | 5,47×10^00 | 7,69×10^−02 | 2,530 | ACTR3,AMPH,AP2A1,AP2A2,Ap2b1,AP2M1,AP2S1,DNM1,HIP1R,SYNJ1 |
| COPI-mediated anterograde transport | 5,47×10^00 | 8,82×10^−02 | 3,000 | Ank2,COPA,COPB1,COPG1,DCTN1,DYNC1H1,SPTBN1,SPTBN2,USO1 |
| Synaptic adhesion-like molecules | 5,42×10^00 | 2,38×10^−01 | 2,236 | DLG4,GRIA1,GRIA3,GRIN1,GRIN2B |
| Glucose metabolism | 5,33×10^00 | 1×10^−01 | 2,121 | HK1,PFKL,PFKM,PFKP,PRKACB,SLC25A11,SLC25A12,TPI1 |
| Oxytocin Signaling Pathway | 5,14×10^00 | 4,96×10^−02 | 2,138 | ATP2B1,ATP2B2,CACNG2,ITPR1,MFN2,MYH10,MYH14,MYH9,MYO18A,PRKACB,PRKCE,PRKCG,RALA,ROCK2 |
| Transcriptional Regulation by MECP2 | 5,04×10^00 | 1,11×10^−01 | −0,816 | CAMK2A,CAMK2D,CAMK2G,GRIA2,GRIN2B,HTT,SLC2A3 |
| Opioid Signalling | 4,95×10^00 | 8,89×10^−02 | 2,646 | ADCY9,CAMK2A,CAMK2D,CAMK2G,ITPR1,NBEA,PRKACB,PRKCG |
| Effects of PIP2 hydrolysis | 4,84×10^00 | 1,85×10^−01 | 2,236 | DGKH,DGKZ,ITPR1,PRKCE,RASGRP1 |
| Adrenergic Receptor Signaling Pathway (Enhanced) | 4,56×10^00 | 5,5×10^−02 | 2,714 | ADCY9,ATP6V0A1,ATP6V1D,ATP6V1H,CACNG2,ITPR1,MAOA,PRKACB,PRKCE,PRKCG,SLC1A2 |
| Huntington's Disease Signaling | 4,45×10^00 | 4,58×10^−02 | 2,236 | AP2A2,DCTN1,DLG4,DNM1,GRIN2B,GRM5,HTT,ITPR1,PRKCE,PRKCG,PSMD1,PSMD6,VTI1B |
| CREB Signaling in Neurons | 4,41×10^00 | 3,29×10^−02 | 4,025 | ADCY9,ADGRB1,ADORA1,BMPR2,CACNG2,CAMK2A,CAMK2D,CAMK2G,GABBR1,GRIA1,GRIA2,GRIA3,GRIN1,GRIN2B,GRM5,ITPR1,PRK |
| Signaling by NTRK1 (TRKA) | 4,32×10^00 | 8,64×10^−02 | 1,134 | AP2A1,AP2A2,AP2M1,AP2S1,CRK,DNM1,RALA |
| mTOR Signaling | 4,3×10^00 | 5,14×10^−02 | 0,447 | AKT1S1,DGKZ,EIF4A2,PRKCE,PRKCG,RALA,RPS11,RPS16,RPS3,RPS5,RPS9 |
| RHO GTPases activate CIT | 4,2×10^00 | 2,11×10^−01 | 2,000 | DLG4,MYH10,MYH14,MYH9 |
| RHO GTPases Activate ROCKs | 4,2×10^00 | 2,11×10^−01 | 2,000 | MYH10,MYH14,MYH9,ROCK2 |
| GNRH Signaling | 4,02×10^00 | 5,24×10^−02 | 2,333 | ADCY9,CACNG2,CAMK2A,CAMK2D,CAMK2G,ITPR1,PRKACB,PRKCE,PRKCG,RALA |
| Fcgamma receptor (FCGR) dependent phagocytosis | 3,97×10^00 | 5,73×10^−02 | 2,121 | ACTR3,CRK,CYFIP1,CYFIP2,ITPR1,MYH9,MYO5A,NCKAP1,PRKCE |
| Synaptic Long Term Depression | 3,89×10^00 | 5,05×10^−02 | 2,530 | CACNG2,GRIA1,GRIA2,GRIA3,GRM5,ITPR1,PRKCE,PRKCG,RALA,RYR2 |
| Orexin Signaling Pathway | 3,88×10^00 | 4,62×10^−02 | 2,111 | ADCY9,AKT1S1,ATP6V0A1,ATP6V1D,ATP6V1H,CACNG2,ITPR1,PI4KA,PRKACB,PRKCE,PRKCG |
| Calcium Transport I | 3,74×10^00 | 3×10^−01 | NA | ATP2A2,ATP2B1,ATP2B2 |
| Translocation of SLC2A4 (GLUT4) to the plasma membrane | 3,72×10^00 | 8,45×10^−02 | 1,633 | EXOC2,EXOC3,EXOC8,MYH9,MYO5A,RALA |
| Insulin processing | 3,71×10^00 | 1,6×10^−01 | 2,000 | EXOC2,EXOC3,EXOC8,MYO5A |
| Docosahexaenoic Acid (DHA) Signaling | 3,7×10^00 | 4,4×10^−02 | 2,111 | ADCY9,AKT1S1,APP,GRIN2B,ITPR1,PRKACB,PRKCE,PRKCG,SPTBN1,SPTBN2,SYNJ1 |
| Melatonin Signaling | 3,68×10^00 | 8,33×10^−02 | 0,000 | CAMK2A,CAMK2D,CAMK2G,PRKACB,PRKCE,PRKCG |
| Beta-catenin independent WNT signaling | 3,68×10^00 | 6,86×10^−02 | 2,646 | AP2A1,AP2A2,AP2M1,AP2S1,CAMK2A,ITPR1,PRKCG |
| Phagosome Formation | 3,61×10^00 | 2,87×10^−02 | 4,025 | ACTR3,ADGRB1,ADORA1,AP1B1,AP1G1,AP1M1,GABBR1,GRM5,ITPR1,MYH10,MYH14,MYH9,MYO18A,PI4KA,PRKCE,PRKCG,RALA,RAS |
| Glycolysis I | 3,57×10^00 | 1,48×10^−01 | 1,000 | PFKL,PFKM,PFKP,TPI1 |
| Plasma lipoprotein assembly, remodeling, and clearance | 3,56×10^00 | 7,89×10^−02 | 2,449 | AP2A1,AP2A2,AP2M1,AP2S1,NCEH1,PRKACB |
| Signaling by VEGF | 3,53×10^00 | 6,48×10^−02 | 1,633 | CRK,CYFIP1,CYFIP2,ITPR1,NCKAP1,PRKACB,ROCK2 |
| α-Adrenergic Signaling | 3,5×10^00 | 6,42×10^−02 | 1,342 | ADCY9,ITPR1,PRKACB,PRKCE,PRKCG,PYGB,RALA |
| RHO GTPase cycle | 3,5×10^00 | 3,33×10^−02 | 3,873 | ABR,ARHGAP26,ARHGEF2,CYFIP1,CYFIP2,DDX39B,FARP1,GIT1,IQGAP2,KALRN,NCKAP1,ROCK2,SPTBN1,SRGAP3,USP9X |
| Role of NFAT in Cardiac Hypertrophy | 3,46×10^00 | 4,46×10^−02 | 2,530 | ADCY9,CACNG2,CAMK2A,CAMK2D,CAMK2G,ITPR1,PRKACB,PRKCE,PRKCG,RALA |
| Thrombin Signaling | 3,45×10^00 | 4,44×10^−02 | 1,897 | ADCY9,ARHGEF2,CAMK2A,CAMK2D,CAMK2G,ITPR1,PRKCE,PRKCG,RALA,ROCK2 |
| Dopamine-DARPP32 Feedback in cAMP Signaling | 3,42×10^00 | 4,84×10^−02 | 2,646 | ADCY9,ATP2A2,CACNG2,GRIN1,GRIN2B,ITPR1,PRKACB,PRKCE,PRKCG |
| Neuroinflammation Signaling Pathway | 3,39×10^00 | 3,79×10^−02 | 1,633 | APP,BMPR2,GABBR1,GABRA1,GABRA5,GABRG2,GRIA1,GRIN1,GRIN2B,PRKCG,SLC1A2,SLC6A11 |
| Amyotrophic Lateral Sclerosis Signaling | 3,34×10^00 | 6,03×10^−02 | 1,633 | CACNG2,GRIA1,GRIA2,GRIA3,GRIN1,GRIN2B,SLC1A2 |
| Signaling by ROBO receptors | 3,34×10^00 | 6,03×10^−02 | 1,890 | CLASP2,MSI1,PRKACB,PSMD1,PSMD6,SLIT2,SRGAP3 |
| The citric acid (TCA) cycle and respiratory electron transport | 3,33×10^00 | 9,09×10^−02 | 1,342 | ACO2,LDHA,ME3,MPC2,PDHX |
| Acetylcholine Receptor Signaling Pathway | 3,29×10^00 | 4,64×10^−02 | 3,000 | ADCY9,CACNG2,CAMK2A,CAMK2D,CAMK2G,ITPR1,PRKACB,PRKCE,PRKCG |
| LGI-ADAM interactions | 3,27×10^00 | 2,14×10^−01 | NA | CACNG2,DLG4,LGI1 |
| Neurexins and neuroligins | 3,25×10^00 | 8,77×10^−02 | 1,342 | DLG4,GRIN1,GRIN2B,GRM5,LIN7C |
| Nitric Oxide Signaling in the Cardiovascular System | 3,25×10^00 | 5,83×10^−02 | 1,342 | ATP2A2,CACNG2,ITPR1,PRKACB,PRKCE,PRKCG,RYR2 |
| CLEAR Signaling Pathway | 3,22×10^00 | 3,86×10^−02 | 0,905 | AKT1S1,ATP6V0A1,ATP6V1D,ATP6V1H,BMPR2,ITPR1,PRKCE,PRKCG,PSAP,RALA,VPS35 |
| GABA receptor activation | 3,15×10^00 | 8,33×10^−02 | 2,236 | ADCY9,GABBR1,GABRA1,GABRA5,GABRG2 |
| Intra-Golgi and retrograde Golgi-to-ER traffic | 3,15×10^00 | 4,43×10^−02 | 1,667 | COPA,COPB1,COPG1,DCTN1,DYNC1H1,KIF1A,PAFAH1B2,PAFAH1B3,RAB3GAP1 |
| Neurotransmitter uptake and metabolism In glial cells | 3,09×10^00 | 5×10^−01 | NA | SLC1A2,SLC38A1 |
| RHO GTPases Activate WASPs and WAVEs | 3,08×10^00 | 1,11×10^−01 | 2,000 | ACTR3,CYFIP1,CYFIP2,NCKAP1 |
| Fcγ Receptor-mediated Phagocytosis in Macrophages and Monocytes | 3,06×10^00 | 6,38×10^−02 | 1,633 | ACTR3,CRK,MYO5A,PRKCE,PRKCG,TLN2 |
| HIF1α Signaling | 3,06×10^00 | 4,31×10^−02 | 2,333 | CAMK2A,CAMK2D,CAMK2G,HK1,LDHA,PRKCE,PRKCG,RALA,SLC2A3 |
| Cellular response to heat stress | 2,99×10^00 | 6,19×10^−02 | 0,816 | AKT1S1,CAMK2A,CAMK2D,CAMK2G,CCAR2,PTGES3 |
| Autism Signaling Pathway | 2,94×10^00 | 3,57×10^−02 | −0,905 | AKT1S1,CACNG2,CAMK2A,CAMK2D,CAMK2G,CYFIP1,GRIN1,GRIN2B,GRM5,PRKACB,RALA |
| RAC Signaling | 2,91×10^00 | 5,11×10^−02 | 1,890 | ACTR3,CYFIP1,CYFIP2,IQGAP2,NCKAP1,PI4KA,RALA |
| RHOGDI Signaling | 2,9×10^00 | 4,09×10^−02 | −2,236 | ACTR3,ARHGEF2,DGKZ,MYH10,MYH14,MYH9,MYO18A,PI4KA,ROCK2 |
| Protein Kinase A Signaling | 2,9×10^00 | 3,16×10^−02 | 2,496 | ADCY9,CAMK2A,CAMK2D,CAMK2G,ITPR1,MYH10,PRKACB,PRKCE,PRKCG,PTPN9,PYGB,ROCK2,RYR2 |
| Sensory processing of sound by inner hair cells of the cochlea | 2,88×10^00 | 7,25×10^−02 | 2,236 | ATP2B1,ATP2B2,MYH9,SPTBN1,SYN1 |
| Tight Junction Signaling | 2,86×10^00 | 4,44×10^−02 | NA | ARHGEF2,JAM3,MYH10,MYH14,MYH9,MYO18A,PRKACB,VTI1B |
| Serotonin Receptor Signaling | 2,84×10^00 | 2,99×10^−02 | 2,673 | ADCY9,APP,CACNG2,CAMK2A,CAMK2D,CAMK2G,KALRN,MAOA,PRKACB,PRKCE,PRKCG,RALA,ROCK2,RYR2 |
| trans-Golgi Network Vesicle Budding | 2,74×10^00 | 6,76×10^−02 | 1,342 | AP1B1,AP1G1,AP1M1,APP,HIP1R |
| RHO GTPases activate PAKs | 2,74×10^00 | 1,43×10^−01 | NA | MYH10,MYH14,MYH9 |
| Xenobiotic Metabolism PXR Signaling Pathway | 2,64×10^00 | 4,1×10^−02 | −1,414 | CAMK2A,CAMK2D,CAMK2G,GSTO1,MAOA,PRKACB,PRKCE,PRKCG |
| Factors Promoting Cardiogenesis in Vertebrates | 2,64×10^00 | 4,58×10^−02 | 2,646 | BMPR2,CAMK2A,CAMK2D,CAMK2G,PRKCE,PRKCG,ROCK2 |
| Chemokine Signaling | 2,57×10^00 | 6,17×10^−02 | 1,342 | CAMK2A,CAMK2D,CAMK2G,RALA,ROCK2 |
| GPCR-Mediated Nutrient Sensing in Enteroendocrine Cells | 2,56×10^00 | 5,08×10^−02 | 2,449 | ADCY9,CACNG2,ITPR1,PRKACB,PRKCE,PRKCG |
| Gustation Pathway | 2,54×10^00 | 3,94×10^−02 | 2,828 | ADCY9,CACNG2,GABRA1,GABRA5,GABRG2,ITPR1,PRKACB,SCN2A |
| UVC-Induced MAPK Signaling | 2,52×10^00 | 7,84×10^−02 | 1,000 | PRKCE,PRKCG,RALA,SMPD3 |
| Renin-Angiotensin Signaling | 2,5×10^00 | 4,96×10^−02 | 1,342 | ADCY9,ITPR1,PRKACB,PRKCE,PRKCG,RALA |
| Phagosome Maturation | 2,47×10^00 | 4,27×10^−02 | NA | ATP6V0A1,ATP6V1D,ATP6V1H,CALR,DYNC1H1,DYNLT3,VTI1B |
| Estrogen Receptor Signaling | 2,45×10^00 | 2,93×10^−02 | 1,414 | ADCY9,CACNG2,DDX5,DLG4,MT-ND4,MT-ND5,NDUFA9,PRKACB,PRKCE,PRKCG,RALA,ROCK2 |
| Glioma Signaling | 2,43×10^00 | 4,8×10^−02 | 1,633 | CAMK2A,CAMK2D,CAMK2G,PRKCE,PRKCG,RALA |
| CXCR4 Signaling | 2,41×10^00 | 4,17×10^−02 | 0,378 | ADCY9,CRK,ITPR1,PRKCE,PRKCG,RALA,ROCK2 |
| HSP90 chaperone cycle for steroid hormone receptors in the presence of ligand | 2,4×10^00 | 7,27×10^−02 | 0,000 | DCTN1,DNAJA2,DYNC1H1,PTGES3 |
| Electron transport, ATP synthesis, and heat production by uncoupling proteins | 2,38×10^00 | 4,69×10^−02 | 1,633 | ACAD9,ATP5PD,MT-ND4,MT-ND5,NDUFA9,UQCRQ |
| RHO GTPases activate PKNs | 2,37×10^00 | 1,07×10^−01 | NA | MYH10,MYH14,MYH9 |
| G Beta Gamma Signaling | 2,36×10^00 | 4,65×10^−02 | 1,342 | CACNG2,ITPR1,PRKACB,PRKCE,PRKCG,RALA |
| Response to elevated platelet cytosolic Ca2+ | 2,32×10^00 | 4,55×10^−02 | −0,816 | APP,CYRIB,MANF,PRKCG,PSAP,VTI1B |
| Iron uptake and transport | 2,31×10^00 | 6,9×10^−02 | 1,000 | ATP6V0A1,ATP6V1D,ATP6V1H,CAND1 |
| Metabolism of water-soluble vitamins and cofactors | 2,29×10^00 | 6,78×10^−02 | 1,000 | ACACA,FASN,GSTO1,SLC2A3 |
| Cardiac Hypertrophy Signaling (Enhanced) | 2,28×10^00 | 2,58×10^−02 | 3,051 | ADCY9,ATP2A2,BMPR2,CACNG2,CAMK2A,CAMK2D,CAMK2G,ITPR1,PRKACB,PRKCE,PRKCG,RALA,ROCK2,RYR2 |
| Gap Junction Signaling | 2,26×10^00 | 3,08×10^−02 | −1,265 | ADCY9,GRIA1,GRIA2,GRIA3,ITPR1,PRKACB,PRKCE,PRKCG,RALA,VDAC3 |
| Eicosanoid Signaling | 2,21×10^00 | 3,21×10^−02 | 2,333 | ADCY9,AP1B1,AP1G1,AP1M1,ATP2A2,PRKACB,PRKCE,PRKCG,RALA |
| Regulation of eIF4 and p70S6K Signaling | 2,2×10^00 | 3,8×10^−02 | NA | EIF4A2,RALA,RPS11,RPS16,RPS3,RPS5,RPS9 |
| Semaphorin interactions | 2,16×10^00 | 6,25×10^−02 | 2,000 | MYH10,MYH14,MYH9,ROCK2 |
| Sirtuin Signaling Pathway | 2,1×10^00 | 3,09×10^−02 | −1,633 | ACLY,APP,LDHA,MT-ND4,MT-ND5,NDUFA9,PFKM,SLC25A4,VDAC3 |
| PPARα/RXRα Activation | 2,08×10^00 | 3,61×10^−02 | 1,633 | ADCY9,AP2A2,BMPR2,CAND1,FASN,PRKACB,RALA |
| Xenobiotic Metabolism Signaling | 2,07×10^00 | 3,06×10^−02 | NA | CAMK2A,CAMK2D,CAMK2G,GSTO1,MAOA,PRKCE,PRKCG,PTGES3,RALA |
| Remodeling of Epithelial Adherens Junctions | 2,07×10^00 | 5,88×10^−02 | NA | ACTR3,DNM1,EXOC2,RALA |
| WNT/SHH Axonal Guidance Signaling Pathway | 2,07×10^00 | 4,03×10^−02 | 0,816 | ADCY9,CAMK2A,CAMK2D,CAMK2G,ITPR1,PRKACB |
| Cargo recognition for clathrin-mediated endocytosis | 2,06×10^00 | 4,67×10^−02 | 2,236 | AP2A1,AP2A2,Ap2b1,AP2M1,AP2S1 |
| GM-CSF Signaling | 2,03×10^00 | 5,71×10^−02 | 1,000 | CAMK2A,CAMK2D,CAMK2G,RALA |
| Corticotropin Releasing Hormone Signaling | 2,03×10^00 | 3,95×10^−02 | 1,342 | ADCY9,CACNG2,ITPR1,PRKACB,PRKCE,PRKCG |
| G-Protein Coupled Receptor Signaling | 2,02×10^00 | 2,28×10^−02 | 2,500 | ADCY9,ADGRB1,ADORA1,CAMK2A,CAMK2D,CAMK2G,CRK,GABBR1,GRM5,PRKACB,PRKCE,PRKCG,RALA,RASGRP1,ROCK2,SYNGAP1 |
| Type II Diabetes Mellitus Signaling | 2,01×10^00 | 3,92×10^−02 | NA | ACSBG1,CACNG2,ITPR1,PRKCE,PRKCG,SMPD3 |
| Advanced glycosylation endproduct receptor signaling | 2×10^00 | 1,54×10^−01 | NA | APP,DDOST |

| | | | | |
|---|---|---|---|---|
| Ephrin Receptor Signaling | 1,99×10^00 | 3,47×10^−02 | 1,134 | ACTR3,CRK,GRIN1,GRIN2B,KALRN,RALA,ROCK2 |
| Necroptosis Signaling Pathway | 1,99×10^00 | 3,87×10^−02 | 1,633 | CAMK2A,CAMK2D,CAMK2G,SLC25A3,SLC25A4,VDAC3 |
| S100 Family Signaling Pathway | 1,98×10^00 | 2,2×10^−02 | 3,153 | ADGRB1,ADORA1,APP,ATP2A2,CACNG2,CAMK2A,CAMK2D,CAMK2G,GABBR1,GRM5,ITPR1,MYH9,PRKACB,PRKCE,PRKCG,ROCK2,RYR2 |
| Oxidative Phosphorylation | 1,98×10^00 | 4,46×10^−02 | 1,342 | ATP5PD,MT-ND4,MT-ND5,NDUFA9,UQCRQ |
| eNOS Signaling | 1,97×10^00 | 3,85×10^−02 | 2,000 | ADCY9,AQP4,ITPR1,PRKACB,PRKCE,PRKCG |
| Epithelial Adherens Junction Signaling | 1,95×10^00 | 3,8×10^−02 | 0,816 | ACTR3,BMPR2,CRK,MYH10,RALA,ROCK2 |
| Acetyl-CoA Biosynthesis III (from Citrate) | 1,93×10^00 | 1×10^00 | NA | ACLY |
| Mechanisms of Viral Exit from Host Cells | 1,91×10^00 | 7,32×10^−02 | NA | PDCD6IP,PRKCE,PRKCG |
| Neuregulin Signaling | 1,9×10^00 | 4,27×10^−02 | 1,000 | CRK,DLG4,PRKCE,PRKCG,RALA |
| Cholecystokinin/Gastrin-mediated Signaling | 1,87×10^00 | 4,2×10^−02 | 1,342 | ITPR1,PRKCE,PRKCG,RALA,ROCK2 |
| Signaling by Insulin receptor | 1,85×10^00 | 6,98×10^−02 | NA | ATP6V0A1,ATP6V1D,ATP6V1H |
| Vasopressin regulates renal water homeostasis via Aquaporins | 1,85×10^00 | 6,98×10^−02 | NA | ADCY9,AQP4,PRKACB |
| G alpha (12/13) signalling events | 1,83×10^00 | 5×10^−02 | 2,000 | ABR,ARHGEF2,KALRN,ROCK2 |
| Rap1 signalling | 1,83×10^00 | 1,25×10^−01 | NA | PRKACB,RASGRP1 |
| Androgen Signaling | 1,82×10^00 | 3,55×10^−02 | 2,236 | CACNG2,CALR,ITPR1,PRKACB,PRKCE,PRKCG |
| Coronavirus Replication Pathway | 1,8×10^00 | 6,67×10^−02 | NA | COPA,COPB1,COPG1 |
| RAN Signaling | 1,78×10^00 | 1,18×10^−01 | NA | IPO5,TNPO1 |
| ROBO SLIT Signaling Pathway | 1,76×10^00 | 3,94×10^−02 | 0,447 | ACTR3,APP,GIT1,PRKACB,SLIT2 |
| 14-3-3-mediated Signaling | 1,76×10^00 | 3,94×10^−02 | 1,000 | AKT1S1,PDCD6IP,PRKCE,PRKCG,RALA |
| Axonal Guidance Signaling | 1,73×10^00 | 2,35×10^−02 | NA | ACTR3,CRK,GIT1,KALRN,MAG,PRKACB,PRKCE,PRKCG,RALA,ROCK2,SLIT2,SRGAP3 |
| Platelet homeostasis | 1,72×10^00 | 4,65×10^−02 | 2,000 | ATP2A2,ATP2B1,ATP2B2,ITPR1 |
| XBP1(S) activates chaperone genes | 1,72×10^00 | 6,25×10^−02 | NA | DCTN1,HYOU1,SSR1 |
| G alpha (z) signalling events | 1,72×10^00 | 6,25×10^−02 | NA | ADCY9,PRKCE,PRKCG |
| nNOS Signaling in Skeletal Muscle Cells | 1,72×10^00 | 6,25×10^−02 | NA | CACNG2,ITPR1,RYR2 |
| fMLP Signaling in Neutrophils | 1,71×10^00 | 3,82×10^−02 | 1,342 | ACTR3,ITPR1,PRKCE,PRKCG,RALA |
| PFKFB4 Signaling Pathway | 1,7×10^00 | 6,12×10^−02 | NA | HK1,PFKM,PRKACB |
| GABA synthesis, release, reuptake and degradation | 1,68×10^00 | 1,05×10^−01 | NA | SLC32A1,SLC6A11 |
| P2Y Purinergic Receptor Signaling Pathway | 1,68×10^00 | 3,76×10^−02 | 1,342 | ADCY9,PRKACB,PRKCE,PRKCG,RALA |
| RAB GEFs exchange GTP for GDP on RABs | 1,68×10^00 | 4,49×10^−02 | 2,000 | CCZ1/CCZ1B,MADD,RAB3GAP1,SBF1 |
| CCR3 Signaling in Eosinophils | 1,67×10^00 | 3,73×10^−02 | NA | ITPR1,PRKCE,PRKCG,RALA,ROCK2 |
| Amyloid Processing | 1,65×10^00 | 5,88×10^−02 | NA | APP,PRKACB,PRKCE |
| cAMP-mediated signaling | 1,65×10^00 | 2,97×10^−02 | 2,646 | ADCY9,ADORA1,CAMK2A,CAMK2D,CAMK2G,GABBR1,PRKACB |
| Crosstalk between Dendritic Cells and Natural Killer Cells | 1,64×10^00 | 4,4×10^−02 | NA | CAMK2A,CAMK2D,CAMK2G,TLN2 |
| Receptor-type tyrosine-protein phosphatases | 1,64×10^00 | 1×10^−01 | NA | PPFIA2,PPFIA3 |
| Fatty Acid Biosynthesis Initiation II | 1,63×10^00 | 5×10^−01 | NA | FASN |
| MAPK6/MAPK4 signaling | 1,63×10^00 | 4,35×10^−02 | NA | KALRN,PRKACB,PSMD1,PSMD6 |
| Iron homeostasis signaling pathway | 1,62×10^00 | 3,62×10^−02 | NA | ACO2,ATP6V0A1,ATP6V1D,ATP6V1H,BMPR2 |
| White Adipose Tissue Browning Pathway | 1,62×10^00 | 3,62×10^−02 | 2,000 | ADCY9,CACNG2,ITPR1,LDHA,PRKACB |
| AMPK Signaling | 1,6×10^00 | 2,89×10^−02 | 0,816 | ACACA,AKT1S1,FASN,PFKL,PFKM,PFKP,PRKACB |
| Gαi Signaling | 1,6×10^00 | 3,57×10^−02 | −0,447 | ADCY9,ADORA1,GABBR1,PRKACB,RALA |
| Insulin Receptor Signaling | 1,6×10^00 | 3,57×10^−02 | −1,342 | ACLY,CRK,PRKACB,RALA,SYNJ1 |
| Sensory processing of sound by outer hair cells of the cochlea | 1,57×10^00 | 5,45×10^−02 | NA | ATP2B2,MYH9,SPTBN1 |
| Leukocyte Extravasation Signaling | 1,57×10^00 | 3,11×10^−02 | 1,342 | CRK,JAM3,PRKCE,PRKCG,RASGRP1,ROCK2 |
| Melanocyte Development and Pigmentation Signaling | 1,54×10^00 | 4,08×10^−02 | 0,000 | ADCY9,CRK,PRKACB,RALA |
| FAT10 Signaling Pathway | 1,53×10^00 | 5,26×10^−02 | NA | HTT,PSMD1,PSMD6 |
| Apelin Cardiomyocyte Signaling Pathway | 1,53×10^00 | 4,04×10^−02 | 2,000 | ATP2A2,ITPR1,PRKCE,PRKCG |
| Vitamin-C Transport | 1,53×10^00 | 8,7×10^−02 | NA | GSTO1,SLC2A3 |
| Endocannabinoid Cancer Inhibition Pathway | 1,52×10^00 | 3,4×10^−02 | −0,447 | ADCY9,AKT1S1,PRKACB,ROCK2,SMPD3 |
| p75 NTR receptor-mediated signalling | 1,51×10^00 | 4×10^−02 | 1,000 | ABR,ARHGEF2,KALRN,MAG |
| Oxytocin in Brain Signaling Pathway | 1,51×10^00 | 3,02×10^−02 | 1,633 | CACNG2,ITPR1,PRKCE,PRKCG,RALA,SLC12A5 |
| Signaling by ERBB4 | 1,49×10^00 | 5,08×10^−02 | NA | DLG4,GABRA1,GABRG2 |
| Semaphorin Neuronal Repulsive Signaling Pathway | 1,49×10^00 | 3,33×10^−02 | −0,447 | FARP1,NCAN,PRKACB,ROCK2,VCAN |
| Integration of energy metabolism | 1,47×10^00 | 3,88×10^−02 | NA | ACLY,ADCY9,ITPR1,PRKACB |
| Cilium Assembly | 1,46×10^00 | 2,94×10^−02 | 2,449 | DCTN1,DYNC1H1,EXOC2,EXOC3,EXOC8,TNPO1 |
| Coronavirus Pathogenesis Pathway | 1,46×10^00 | 2,94×10^−02 | −1,633 | RPS11,RPS16,RPS3,RPS5,RPS9,TNPO1 |
| Cellular Effects of Sildenafil (Viagra) | 1,46×10^00 | 2,03×10^−02 | 0,000 | ADCY9,ADGRB1,ADORA1,ATP2A2,BMPR2,CACNG2,GABBR1,GRM5,MYH10,MYH14,MYH9,MYO18A,PRKACB,PRKCE |
| MTF1 activates gene expression | 1,46×10^00 | 3,33×10^−01 | NA | CSRP1 |
| Pyrophosphate hydrolysis | 1,46×10^00 | 3,33×10^−01 | NA | LHPP |
| Palmitate Biosynthesis I (Animals) | 1,46×10^00 | 3,33×10^−01 | NA | FASN |
| Ascorbate Recycling (Cytosolic) | 1,46×10^00 | 3,33×10^−01 | NA | GSTO1 |
| Anandamide Degradation | 1,46×10^00 | 3,33×10^−01 | NA | FAAH |
| Biotin-carboxyl Carrier Protein Assembly | 1,46×10^00 | 3,33×10^−01 | NA | ACACA |
| NCAM signaling for neurite out-growth | 1,45×10^00 | 4,92×10^−02 | NA | NCAN,SPTBN1,SPTBN2 |
| 3-phosphoinositide Biosynthesis | 1,42×10^00 | 2,87×10^−02 | 2,236 | HACD2,PI4KA,PLPPR4,PPFIA2,PPFIA3,SYNJ1 |
| Thrombopoietin Signaling | 1,42×10^00 | 4,76×10^−02 | NA | PRKCE,PRKCG,RALA |
| Mitochondrial protein import | 1,4×10^00 | 4,69×10^−02 | NA | ACO2,SLC25A12,SLC25A4 |
| Integrin signaling | 1,4×10^00 | 7,41×10^−02 | NA | CRK,RASGRP1 |
| Integrin Signaling | 1,4×10^00 | 2,83×10^−02 | −0,447 | ACTR3,ARHGAP26,CRK,GIT1,RALA,TLN2 |
| Myelination Signaling Pathway | 1,39×10^00 | 2,45×10^−02 | −0,707 | AKT1S1,BMPR2,FASN,MAG,PRKACB,RALA,RAPGEF2,ROCK2 |
| ERK/MAPK Signaling | 1,37×10^00 | 2,79×10^−02 | 0,816 | CRK,PRKACB,PRKCE,PRKCG,RALA,TLN2 |
| Hedgehog 'off' state | 1,35×10^00 | 3,54×10^−02 | 2,000 | ADCY9,PRKACB,PSMD1,PSMD6 |
| Role of MAPK Signaling in Promoting the Pathogenesis of Influenza | 1,34×10^00 | 3,51×10^−02 | 0,000 | ATP6V0A1,ATP6V1D,ATP6V1H,RALA |
| Threonine catabolism | 1,33×10^00 | 2,5×10^−01 | NA | RIDA |
| Melatonin Degradation II | 1,33×10^00 | 2,5×10^−01 | NA | MAOA |
| ERBB4 Signaling | 1,33×10^00 | 4,41×10^−02 | NA | PRKCE,PRKCG,RALA |
| Molecular Mechanisms of Cancer | 1,32×10^00 | 1,87×10^−02 | 3,000 | ADCY9,ADGRB1,ADORA1,ARHGEF2,BMPR2,CAMK2A,CAMK2D,CAMK2G,CRK,GABBR1,GRM5,PRKACB,PRKCE,PRKCG,RALA,RASGRP1 |
| TP53 Regulates Transcription of DNA Repair Genes | 1,3×10^00 | 4,29×10^−02 | NA | AP1B1,AP1G1,AP1M1 |
| Breast Cancer Regulation by Stathmin1 | 1,3×10^00 | 2,02×10^−02 | 0,577 | ADGRB1,ADORA1,ARHGEF2,CAMK2A,CAMK2D,CAMK2G,GABBR1,GRM5,PRKACB,PRKCE,PRKCG,RALA |
| Fc Epsilon RI Signaling | 1,29×10^00 | 3,39×10^−02 | 0,000 | PRKCE,PRKCG,RALA,SYNJ1 |
| Gαq Signaling | 1,29×10^00 | 2,94×10^−02 | 2,236 | GRM5,ITPR1,PRKCE,PRKCG,ROCK2 |
| Stearate Biosynthesis I (Animals) | 1,29×10^00 | 4,23×10^−02 | NA | ACSBG1,FASN,TECR |
| Aldosterone Signaling in Epithelial Cells | 1,27×10^00 | 2,91×10^−02 | 2,236 | ITPR1,PI4KA,PRKCE,PRKCG,SLC12A2 |
| NGF Signaling | 1,27×10^00 | 3,33×10^−02 | NA | CRK,RALA,ROCK2,SMPD3 |
| RIPK1-mediated regulated necrosis | 1,26×10^00 | 6,25×10^−02 | NA | OGT,PDCD6IP |
| rRNA processing | 1,26×10^00 | 6,25×10^−02 | NA | MT-ND4,MT-ND5 |
| Granzyme A Signaling | 1,24×10^00 | 4,05×10^−02 | NA | MT-ND4,MT-ND5,NDUFA9 |
| Arsenate Detoxification I (Glutaredoxin) | 1,24×10^00 | 2×10^−01 | NA | GSTO1 |
| Pentose Phosphate Pathway (Oxidative Branch) | 1,24×10^00 | 2×10^−01 | NA | PGD |
| CMP-N-acetylneuraminate Biosynthesis I (Eukaryotes) | 1,24×10^00 | 2×10^−01 | NA | CMAS |
| Caveolar-mediated Endocytosis Signaling | 1,23×10^00 | 4×10^−02 | NA | COPA,COPB1,COPG1 |
| GPCR-Mediated Integration of Enteroendocrine Signaling Exemplified by an L Cell | 1,23×10^00 | 4×10^−02 | NA | ADCY9,ITPR1,PRKACB |
| RHOA Signaling | 1,23×10^00 | 3,23×10^−02 | 2,000 | ACTR3,PI4KA,RAPGEF2,ROCK2 |
| Gap junction trafficking and regulation | 1,22×10^00 | 5,88×10^−02 | NA | AP2M1,DNM1 |
| Macropinocytosis Signaling | 1,22×10^00 | 3,95×10^−02 | NA | PRKCE,PRKCG,RALA |
| Gαs Signaling | 1,21×10^00 | 3,17×10^−02 | 2,000 | ADCY9,PRKACB,RAPGEF2,RYR2 |
| Cardiac β-adrenergic Signaling | 1,2×10^00 | 2,78×10^−02 | NA | ADCY9,ATP2A2,CACNG2,PRKACB,RYR2 |
| NRF2-mediated Oxidative Stress Response | 1,2×10^00 | 2,53×10^−02 | 1,000 | DNAJA2,GSTO1,HACD3,PRKCE,PRKCG,RALA |
| Superpathway of Inositol Phosphate Compounds | 1,2×10^00 | 2,53×10^−02 | 2,236 | HACD2,PI4KA,PLPPR4,PPFIA2,PPFIA3,SYNJ1 |
| Endocannabinoid Developing Neuron Pathway | 1,2×10^00 | 3,15×10^−02 | NA | ADCY9,AKT1S1,PRKACB,RALA |
| NF-κB Activation by Viruses | 1,19×10^00 | 3,85×10^−02 | NA | PRKCE,PRKCG,RALA |
| D-myo-inositol (1,4,5,6)-Tetrakisphosphate Biosynthesis | 1,18×10^00 | 2,73×10^−02 | 2,000 | HACD2,PLPPR4,PPFIA2,PPFIA3,SYNJ1 |
| D-myo-inositol (3,4,5,6)-tetrakisphosphate Biosynthesis | 1,18×10^00 | 2,73×10^−02 | 2,000 | HACD2,PLPPR4,PPFIA2,PPFIA3,SYNJ1 |
| IL-3 Signaling | 1,18×10^00 | 3,8×10^−02 | NA | PRKCE,PRKCG,RALA |
| Creatine metabolism | 1,16×10^00 | 1,67×10^−01 | NA | SLC6A11 |
| Pyruvate Fermentation to Lactate | 1,16×10^00 | 1,67×10^−01 | NA | LDHA |
| Glycerol Degradation I | 1,16×10^00 | 1,67×10^−01 | NA | Gk |
| Dopamine Receptor Signaling | 1,16×10^00 | 3,75×10^−02 | NA | ADCY9,MAOA,PRKACB |
| Gene and protein expression by JAK-STAT signaling after IL-12 stimulation | 1,15×10^00 | 5,41×10^−02 | NA | GSTO1,RALA |
| Cachexia Signaling Pathway | 1,15×10^00 | 2,17×10^−02 | 2,121 | ADCY9,BMPR2,MFN2,PRKACB,PRKCE,PRKCG,PSMD1,PSMD6 |
| Neddylation | 1,14×10^00 | 2,44×10^−02 | 2,449 | CAND1,CUL5,DDB1,NAE1,PSMD1,PSMD6 |
| PI Metabolism | 1,14×10^00 | 3,66×10^−02 | NA | PI4KA,SBF1,SYNJ1 |
| VEGF Family Ligand-Receptor Interactions | 1,11×10^00 | 3,57×10^−02 | NA | PRKCE,PRKCG,RALA |
| tRNA Charging | 1,11×10^00 | 5,13×10^−02 | NA | EPRS1,IARS2 |
| DHCR24 Signaling Pathway | 1,1×10^00 | 2,92×10^−02 | 1,000 | APP,PRKCE,PRKCG,RALA |
| LPS-stimulated MAPK Signaling | 1,1×10^00 | 3,53×10^−02 | NA | PRKCE,PRKCG,RALA |
| Pentose phosphate pathway | 1,1×10^00 | 1,43×10^−01 | NA | PGD |
| Aryl hydrocarbon receptor signalling | 1,1×10^00 | 1,43×10^−01 | NA | PTGES3 |
| Trehalose Degradation II (Trehalase) | 1,1×10^00 | 1,43×10^−01 | NA | HK1 |
| Endothelin-1 Signaling | 1,1×10^00 | 2,58×10^−02 | 0,447 | ADCY9,ITPR1,PRKCE,PRKCG,RALA |
| 3-phosphoinositide Degradation | 1,1×10^00 | 2,58×10^−02 | 2,000 | HACD2,PLPPR4,PPFIA2,PPFIA3,SYNJ1 |
| Class C/3 (Metabotropic glutamate/pheromone receptors) | 1,09×10^00 | 5×10^−02 | NA | GABBR1,GRM5 |
| Ephrin A Signaling | 1,08×10^00 | 2,88×10^−02 | 0,000 | APP,GIT1,PRKACB,ROCK2 |
| PDGF Signaling | 1,08×10^00 | 3,45×10^−02 | NA | CRK,RALA,SYNJ1 |
| Hematoma Resolution Signaling Pathway | 1,07×10^00 | 2,33×10^−02 | 0,816 | GRIN1,GRIN2B,MT-ND4,MT-ND5,NDUFA9,VPS35 |
| Apelin Endothelial Signaling Pathway | 1,07×10^00 | 2,84×10^−02 | 0,000 | ADCY9,PRKCE,PRKCG,RALA |
| D-myo-inositol-5-phosphate Metabolism | 1,06×10^00 | 2,51×10^−02 | 2,000 | HACD2,PLPPR4,PPFIA2,PPFIA3,SYNJ1 |
| Xenobiotic Metabolism General Signaling Pathway | 1,05×10^00 | 2,8×10^−02 | 0,000 | GSTO1,PRKCE,PRKCG,RALA |
| Miro GTPase Cycle | 1,04×10^00 | 1,25×10^−01 | NA | MFN2 |
| NFE2L2 regulates pentose phosphate pathway genes | 1,04×10^00 | 1,25×10^−01 | NA | PGD |
| Sphingomyelin Metabolism | 1,04×10^00 | 1,25×10^−01 | NA | SMPD3 |
| BMP signaling pathway | 1,03×10^00 | 3,3×10^−02 | NA | BMPR2,PRKACB,RALA |
| C-type lectin receptors (CLRs) | 1,03×10^00 | 2,76×10^−02 | 2,000 | ITPR1,PRKACB,PSMD1,PSMD6 |
| ABRA Signaling Pathway | 1,02×10^00 | 3,26×10^−02 | NA | ACTR3,MYH9,ROCK2 |
| Actin Nucleation by ARP-WASP Complex | 1,01×10^00 | 3,23×10^−02 | NA | ACTR3,RALA,ROCK2 |
| ERBB Signaling | 1,01×10^00 | 3,23×10^−02 | NA | PRKCE,PRKCG,RALA |
| GPER1 signaling | 1,01×10^00 | 4,44×10^−02 | NA | ADCY9,PRKACB |
| Interferon gamma signaling | 1×10^00 | 3,19×10^−02 | NA | CAMK2A,CAMK2D,CAMK2G |
| Role of Macrophages, Fibroblasts and Endothelial Cells in Rheumatoid Arthritis | 9,99×10^−01 | 2,1×10^−02 | 1,890 | CAMK2A,CAMK2D,CAMK2G,PRKCE,PRKCG,RALA,ROCK2 |
| Sucrose Degradation V (Mammalian) | 9,95×10^−01 | 1,11×10^−01 | NA | TPI1 |
| Prolactin Signaling | 9,91×10^−01 | 3,16×10^−02 | NA | PRKCE,PRKCG,RALA |
| Neutrophil Extracellular Trap Signaling Pathway | 9,91×10^−01 | 2×10^−02 | 2,828 | ITPR1,MT-ND4,MT-ND5,NDUFA9,PRKCE,PRKCG,SLC25A4,VDAC3 |
| Agranulocyte Adhesion and Diapedesis | 9,86×10^−01 | 2,38×10^−02 | NA | JAM3,MYH10,MYH14,MYH9,MYO18A |
| Neurotransmitter clearance | 9,52×10^−01 | 1×10^−01 | NA | MAOA |
| ATF6 (ATF6-alpha) activates chaperone genes | 9,52×10^−01 | 1×10^−01 | NA | CALR |
| Prostanoid Biosynthesis | 9,52×10^−01 | 1×10^−01 | NA | PTGES3 |

| | | | | |
|---|---|---|---|---|
| Transcriptional activity of SMAD2/SMAD3:SMAD4 heterotrimer | 9,45×10^-01 | 4,08×10^-02 | NA | NEDD4L,USP9X |
| ABC-family proteins mediated transport | 9,32×10^-01 | 2,97×10^-02 | NA | ABCB7,PSMD1,PSMD6 |
| Signaling by ERBB2 | 9,3×10^-01 | 4×10^-02 | NA | CUL5,PRKCE |
| Netrin-1 signaling | 9,3×10^-01 | 4×10^-02 | NA | AGAP2,SLIT2 |
| Neurotransmitter receptors and postsynaptic signal transmission | 9,13×10^-01 | 9,09×10^-02 | NA | NBEA |
| Aspartate and asparagine metabolism | 9,13×10^-01 | 9,09×10^-02 | NA | SLC25A12 |
| Pentose Phosphate Pathway | 9,13×10^-01 | 9,09×10^-02 | NA | PGD |
| Glycogen Degradation II | 9,13×10^-01 | 9,09×10^-02 | NA | PYGB |
| GDP-glucose Biosynthesis | 9,13×10^-01 | 9,09×10^-02 | NA | HK1 |
| UVB-Induced MAPK Signaling | 9,02×10^-01 | 3,85×10^-02 | NA | PRKCE,PRKCG |
| Regulation of Apoptosis | 8,89×10^-01 | 3,77×10^-02 | NA | PSMD1,PSMD6 |
| NAFLD Signaling Pathway | 8,88×10^-01 | 2,21×10^-02 | 0,447 | ACACA,AKT1S1,ATP2A2,FASN,PRKCE |
| WNK Renal Signaling Pathway | 8,86×10^-01 | 2,83×10^-02 | NA | PRKCE,PRKCG,SLC12A5 |
| Extracellular matrix organization | 8,86×10^-01 | 2,83×10^-02 | NA | APP,NCAN,VCAN |
| Paxillin Signaling | 8,77×10^-01 | 2,8×10^-02 | NA | CRK,RALA,TLN2 |
| Presynaptic depolarization and calcium channel opening | 8,77×10^-01 | 8,33×10^-02 | NA | CACNG2 |
| Glucose and Glucose-1-phosphate Degradation | 8,77×10^-01 | 8,33×10^-02 | NA | HK1 |
| RAR Activation | 8,66×10^-01 | 1,86×10^-02 | 2,121 | ADCY9,BMPR2,CAMK2A,CAMK2D,CAMK2G,GRIA1,PRKACB,RPL7A |
| Interleukin-3, Interleukin-5 and GM-CSF signaling | 8,63×10^-01 | 3,64×10^-02 | NA | CRK,SYNJ1 |
| Amino acids regulate mTORC1 | 8,63×10^-01 | 3,64×10^-02 | NA | ATP6V1D,ATP6V1H |
| Triacylglycerol Biosynthesis | 8,5×10^-01 | 3,57×10^-02 | NA | PLPP3,PLPPR4 |
| Signaling by the B Cell Receptor (BCR) | 8,49×10^-01 | 2,35×10^-02 | NA | ITPR1,PSMD1,PSMD6,RASGRP1 |
| Cytosolic iron-sulfur cluster assembly | 8,45×10^-01 | 7,69×10^-02 | NA | ABCB7 |
| Passive transport by Aquaporins | 8,45×10^-01 | 7,69×10^-02 | NA | AQP4 |
| Mitochondrial calcium ion transport | 8,45×10^-01 | 7,69×10^-02 | NA | VDAC3 |
| Glycogen Degradation III | 8,45×10^-01 | 7,69×10^-02 | NA | PYGB |
| UDP-N-acetyl-D-galactosamine Biosynthesis II | 8,45×10^-01 | 7,69×10^-02 | NA | HK1 |
| TNF signaling | 8,38×10^-01 | 3,51×10^-02 | NA | MADD,SMPD3 |
| Mitotic Metaphase and Anaphase | 8,37×10^-01 | 2,13×10^-02 | 2,236 | CLASP2,DYNC1H1,PSMD1,PSMD6,TNPO1 |
| Carnitine metabolism | 8,15×10^-01 | 7,14×10^-02 | NA | ACACA |
| Phenylalanine Degradation IV (Mammalian, via Side Chain) | 8,15×10^-01 | 7,14×10^-02 | NA | MAOA |
| Metabolism of polyamines | 8,14×10^-01 | 3,39×10^-02 | NA | PSMD1,PSMD6 |
| CDK5 Signaling | 8,11×10^-01 | 2,61×10^-02 | NA | ADCY9,PRKACB,RALA |
| Erythropoietin Signaling Pathway | 8,1×10^-01 | 2,27×10^-02 | 1,000 | ITPR1,PRKCE,PRKCG,RALA |
| NIK-->noncanonical NF-kB signaling | 8,03×10^-01 | 3,33×10^-02 | NA | PSMD1,PSMD6 |
| Metabolism of nitric oxide: NOS3 activation and regulation | 7,88×10^-01 | 6,67×10^-02 | NA | DDAH1 |
| Synthesis of Prostaglandins (PG) and Thromboxanes (TX) | 7,88×10^-01 | 6,67×10^-02 | NA | PTGES3 |
| Fatty Acid Activation | 7,88×10^-01 | 6,67×10^-02 | NA | ACSBG1 |
| UFMylation Signaling Pathway | 7,8×10^-01 | 3,23×10^-02 | NA | EIF4A2,RPL26 |
| D-myo-inositol (1,4,5)-trisphosphate Degradation | 7,62×10^-01 | 6,25×10^-02 | NA | SYNJ1 |
| Hedgehog ligand biogenesis | 7,49×10^-01 | 3,08×10^-02 | NA | PSMD1,PSMD6 |
| Induction of Apoptosis by HIV1 | 7,49×10^-01 | 3,08×10^-02 | NA | SLC25A3,SLC25A4 |
| MicroRNA Biogenesis Signaling Pathway | 7,45×10^-01 | 2,14×10^-02 | −1,000 | DDX5,DNAJA2,PTGES3,RALA |
| 1D-myo-inositol Hexakisphosphate Biosynthesis II (Mammalian) | 7,39×10^-01 | 5,88×10^-02 | NA | SYNJ1 |
| D-myo-inositol (1,3,4)-trisphosphate Biosynthesis | 7,39×10^-01 | 5,88×10^-02 | NA | SYNJ1 |
| Sperm Motility | 7,27×10^-01 | 1,95×10^-02 | 2,000 | ITPR1,PRKACB,PRKCE,PRKCG,SLC12A2 |
| GP6 Signaling Pathway | 7,23×10^-01 | 2,36×10^-02 | NA | ITPR1,PRKCE,PRKCG |
| G alpha (i) signalling events | 7,22×10^-01 | 1,94×10^-02 | 0,447 | ADCY9,ADORA1,APP,GABBR1,PSAP |
| TNFR2 non-canonical NF-kB pathway | 7,19×10^-01 | 2,94×10^-02 | NA | PSMD1,PSMD6 |
| TR/RXR Activation | 7,16×10^-01 | 2,34×10^-02 | NA | ACACA,FASN,PFKP |
| Interleukin-1 family signaling | 7,1×10^-01 | 2,33×10^-02 | NA | PSMD1,PSMD6,PTPN9 |
| Cardiac Hypertrophy Signaling | 7,08×10^-01 | 1,92×10^-02 | 1,000 | ADCY9,CACNG2,PRKACB,RALA,ROCK2 |
| Hepatic Fibrosis / Hepatic Stellate Cell Activation | 7,07×10^-01 | 2,06×10^-02 | NA | MYH10,MYH14,MYH9,MYO18A |
| Deubiquitination | 6,99×10^-01 | 1,9×10^-02 | 2,236 | OGT,PSMD1,PSMD6,USP9X,VDAC3 |
| Ferroptosis Signaling Pathway | 6,96×10^-01 | 2,29×10^-02 | NA | ACACA,RALA,SLC38A1 |
| Metabolism of cofactors | 6,95×10^-01 | 5,26×10^-02 | NA | COQ9 |
| γ-linolenate Biosynthesis II (Animals) | 6,95×10^-01 | 5,26×10^-02 | NA | ACSBG1 |
| Mitochondrial L-carnitine Shuttle Pathway | 6,95×10^-01 | 5,26×10^-02 | NA | ACSBG1 |
| Growth Hormone Signaling | 6,9×10^-01 | 2,82×10^-02 | NA | PRKCE,PRKCG |
| HGF Signaling | 6,9×10^-01 | 2,27×10^-02 | NA | PRKCE,PRKCG,RALA |
| Signaling by Rho Family GTPases | 6,82×10^-01 | 1,87×10^-02 | 2,236 | ACTR3,ARHGEF2,CYFIP1,PI4KA,ROCK2 |
| COPII-mediated vesicle transport | 6,81×10^-01 | 2,78×10^-02 | NA | GRIA1,USO1 |
| Ephrin B Signaling | 6,81×10^-01 | 2,78×10^-02 | NA | KALRN,ROCK2 |
| Pulmonary Healing Signaling Pathway | 6,81×10^-01 | 2,01×10^-02 | 1,000 | BMPR2,PRKCE,PRKCG,RALA |
| Mitotic G2-G2/M phases | 6,81×10^-01 | 2,01×10^-02 | 2,000 | DCTN1,DYNC1H1,PSMD1,PSMD6 |
| Adrenomedullin signaling pathway | 6,81×10^-01 | 2,01×10^-02 | 1,000 | ADCY9,ITPR1,PRKACB,RALA |
| Class I peroxisomal membrane protein import | 6,75×10^-01 | 5×10^-02 | NA | GDAP1 |
| Regulation of RUNX2 expression and activity | 6,72×10^-01 | 2,74×10^-02 | NA | PSMD1,PSMD6 |
| Amyloid fiber formation | 6,72×10^-01 | 2,74×10^-02 | NA | APP,USP9X |
| Human Embryonic Stem Cell Pluripotency | 6,71×10^-01 | 1,99×10^-02 | 1,000 | BMPR2,PRKCE,PRKCG,RALA |
| ILK Signaling | 6,71×10^-01 | 1,99×10^-02 | NA | MYH10,MYH14,MYH9,MYO18A |
| Lipid Antigen Presentation by CD1 | 6,58×10^-01 | 1,69×10^-02 | 1,134 | AP2A1,AP2A2,Ap2b1,AP2M1,AP2S1,CALR,PSAP |
| Endoplasmic Reticulum Stress Pathway | 6,57×10^-01 | 4,76×10^-02 | NA | CALR |
| Protein Ubiquitination Pathway | 6,56×10^-01 | 1,83×10^-02 | 2,000 | NEDD4L,PSMD1,PSMD6,USO1,USP9X |
| Cellular response to hypoxia | 6,55×10^-01 | 2,67×10^-02 | NA | PSMD1,PSMD6 |
| Hepatic Fibrosis Signaling Pathway | 6,52×10^-01 | 1,68×10^-02 | 0,816 | BMPR2,CACNG2,PRKACB,PRKCE,PRKCG,RALA,ROCK2 |
| Angiopoietin Signaling | 6,47×10^-01 | 2,63×10^-02 | NA | CRK,RALA |
| Leptin Signaling in Obesity | 6,47×10^-01 | 2,63×10^-02 | NA | ADCY9,PRKACB |
| GDNF Family Ligand-Receptor Interactions | 6,47×10^-01 | 2,63×10^-02 | NA | ITPR1,RALA |
| RHO GTPases Activate Formins | 6,47×10^-01 | 2,16×10^-02 | NA | CLASP2,DYNC1H1,SCAI |
| Fc epsilon receptor (FCERI) signaling | 6,46×10^-01 | 1,94×10^-02 | NA | ITPR1,PSMD1,PSMD6,RASGRP1 |
| CGAS-STING Signaling Pathway | 6,41×10^-01 | 2,14×10^-02 | NA | ATP6V0A1,ATP6V1D,ATP6V1H |
| Cellular hexose transport | 6,39×10^-01 | 4,55×10^-02 | NA | SLC2A3 |
| Insertion of tail-anchored proteins into the endoplasmic reticulum membrane | 6,39×10^-01 | 4,55×10^-02 | NA | APP |
| Superpathway of D-myo-inositol (1,4,5)-trisphosphate Metabolism | 6,39×10^-01 | 4,55×10^-02 | NA | SYNJ1 |
| Pregnenolone Biosynthesis | 6,39×10^-01 | 4,55×10^-02 | NA | CYP46A1 |
| Putrescine Degradation III | 6,39×10^-01 | 4,55×10^-02 | NA | MAOA |
| VDR/RXR Activation | 6,3×10^-01 | 2,56×10^-02 | NA | PRKCE,PRKCG |
| IL-8 Signaling | 6,27×10^-01 | 1,9×10^-02 | 1,000 | PRKCE,PRKCG,RALA,ROCK2 |
| Renal Cell Carcinoma Signaling | 6,22×10^-01 | 2,53×10^-02 | NA | CRK,RALA |
| RAF-independent MAPK1/3 activation | 6,22×10^-01 | 4,35×10^-02 | NA | PEA15 |
| Nephrin family interactions | 6,22×10^-01 | 4,35×10^-02 | NA | SPTBN1 |
| TCA Cycle II (Eukaryotic) | 6,22×10^-01 | 4,35×10^-02 | NA | ACO2 |
| Histidine Degradation VI | 6,22×10^-01 | 4,35×10^-02 | NA | CYP46A1 |
| Glycosaminoglycan metabolism | 6,14×10^-01 | 2,5×10^-02 | NA | NCAN,VCAN |
| Glycogen metabolism | 6,06×10^-01 | 4,17×10^-02 | NA | PYGB |
| Transport of bile salts and organic acids, metal ions and amine compounds | 5,92×10^-01 | 2,41×10^-02 | NA | SLC30A1,SLC6A11 |
| Signaling by NOTCH4 | 5,92×10^-01 | 2,41×10^-02 | NA | PSMD1,PSMD6 |
| Signaling by NTRK2 (TRKB) | 5,91×10^-01 | 4×10^-02 | NA | GRIN2B |
| PEDF Signaling | 5,84×10^-01 | 2,38×10^-02 | NA | RALA,ROCK2 |
| Nuclear Cytoskeleton Signaling Pathway | 5,83×10^-01 | 1,82×10^-02 | 2,000 | DCTN1,DYNC1H1,KIF1A,PLEC |
| PTEN Signaling | 5,8×10^-01 | 1,99×10^-02 | NA | BMPR2,RALA,SYNJ1 |
| BAG2 Signaling Pathway | 5,77×10^-01 | 2,35×10^-02 | NA | PSMD1,PSMD6 |
| Signaling by BMP | 5,76×10^-01 | 3,85×10^-02 | NA | BMPR2 |
| WNT ligand biogenesis and trafficking | 5,76×10^-01 | 3,85×10^-02 | NA | VPS35 |
| Ubiquinol-10 Biosynthesis (Eukaryotic) | 5,76×10^-01 | 3,85×10^-02 | NA | CYP46A1 |
| D-myo-inositol (1,4,5)-Trisphosphate Biosynthesis | 5,76×10^-01 | 3,85×10^-02 | NA | PI4KA |
| Hepatic Cholestasis | 5,7×10^-01 | 1,79×10^-02 | 2,000 | ADCY9,PRKACB,PRKCE,PRKCG |
| Hedgehog 'on' state | 5,7×10^-01 | 2,33×10^-02 | NA | PSMD1,PSMD6 |
| FGF Signaling | 5,7×10^-01 | 2,33×10^-02 | NA | CRK,ITPR1 |
| Sphingolipid metabolism | 5,63×10^-01 | 2,3×10^-02 | NA | PSAP,SMPD3 |
| Xenobiotic Metabolism AHR Signaling Pathway | 5,63×10^-01 | 2,3×10^-02 | NA | GSTO1,PTGES3 |
| Cardiomyocyte Differentiation via BMP Receptors | 5,62×10^-01 | 3,7×10^-02 | NA | BMPR2 |
| Tryptophan Degradation X (Mammalian, via Tryptamine) | 5,62×10^-01 | 3,7×10^-02 | NA | MAOA |
| Gluconeogenesis I | 5,62×10^-01 | 3,7×10^-02 | NA | ME3 |
| Regulation of mitotic cell cycle | 5,56×10^-01 | 2,27×10^-02 | NA | PSMD1,PSMD6 |
| HER-2 Signaling in Breast Cancer | 5,54×10^-01 | 1,76×10^-02 | 0,000 | AKT1S1,PRKCE,PRKCG,RALA |
| MTOR signalling | 5,49×10^-01 | 3,57×10^-02 | NA | AKT1S1 |
| Glyoxylate metabolism and glycine degradation | 5,49×10^-01 | 3,57×10^-02 | NA | PDHX |
| CDP-diacylglycerol Biosynthesis I | 5,49×10^-01 | 3,57×10^-02 | NA | CDS2 |
| Regulation of Cellular Mechanics by Calpain Protease | 5,43×10^-01 | 2,22×10^-02 | NA | RALA,TLN2 |
| Unfolded protein response | 5,43×10^-01 | 2,22×10^-02 | NA | CALR,DNAJA2 |
| CTLA4 Signaling in Cytotoxic T Lymphocytes | 5,37×10^-01 | 1,48×10^-02 | 3,000 | AP1B1,AP1G1,AP1M1,AP2A1,AP2A2,Ap2b1,AP2M1,AP2S1,RALA |
| KEAP1-NFE2L2 pathway | 5,36×10^-01 | 2,2×10^-02 | NA | PSMD1,PSMD6 |
| Factors involved in megakaryocyte development and platelet production | 5,36×10^-01 | 2,2×10^-02 | NA | MFN2,PRKACB |
| Ceramide Signaling | 5,36×10^-01 | 2,2×10^-02 | NA | RALA,SMPD3 |
| Apelin Adipocyte Signaling Pathway | 5,36×10^-01 | 2,2×10^-02 | NA | ADCY9,PRKACB |
| EGR2 and SOX10-mediated initiation of Schwann cell myelination | 5,36×10^-01 | 3,45×10^-02 | NA | MAG |
| Degradation of beta-catenin by the destruction complex | 5,3×10^-01 | 2,17×10^-02 | NA | PSMD1,PSMD6 |
| Activation of kainate receptors upon glutamate binding | 5,24×10^-01 | 3,33×10^-02 | NA | DLG4 |
| Mitophagy | 5,24×10^-01 | 3,33×10^-02 | NA | MFN2 |
| Protein ubiquitination | 5,24×10^-01 | 3,33×10^-02 | NA | USP9X |
| Signaling by CSF3 (G-CSF) | 5,24×10^-01 | 3,33×10^-02 | NA | CUL5 |
| Phosphatidylglycerol Biosynthesis II (Non-plastidic) | 5,24×10^-01 | 3,33×10^-02 | NA | CDS2 |
| Inhibition of ARE-Mediated mRNA Degradation Pathway | 5,21×10^-01 | 1,84×10^-02 | NA | PRKACB,PSMD1,PSMD6 |
| Sertoli Cell-Germ Cell Junction Signaling Pathway (Enhanced) | 5,18×10^-01 | 1,69×10^-02 | 1,000 | ACTR3,JAM3,PRKACB,RALA |
| Non-Small Cell Lung Cancer Signaling | 5,17×10^-01 | 2,13×10^-02 | NA | ITPR1,RALA |
| Transcriptional Regulatory Network in Embryonic Stem Cells | 5,16×10^-01 | 1,83×10^-02 | NA | BMPR2,GRIN1,RALA |
| Toll-like Receptor Cascades | 5,12×10^-01 | 3,23×10^-02 | NA | DNM1 |
| Bile acid and bile salt metabolism | 5,12×10^-01 | 3,23×10^-02 | NA | CYP46A1 |
| Asparagine N-linked glycosylation | 5,12×10^-01 | 3,23×10^-02 | NA | CALR |
| RHO GTPases activate IQGAPs | 5,12×10^-01 | 3,23×10^-02 | NA | IQGAP2 |
| Cristae formation | 5,12×10^-01 | 3,23×10^-02 | NA | ATP5PD |
| Sonic Hedgehog Signaling | 5,12×10^-01 | 3,23×10^-02 | NA | PRKACB |
| Transcriptional regulation by RUNX3 | 5,05×10^-01 | 2,08×10^-02 | NA | PSMD1,PSMD6 |

| | | | | |
|---|---|---|---|---|
| IL-1 Signaling | 5,05×10^−01 | 2,08×10^−02 | NA | ADCY9,PRKACB |
| Toll Like Receptor 3 (TLR3) Cascade | 5,01×10^−01 | 3,12×10^−02 | NA | SARM1 |
| Dopamine Degradation | 5,01×10^−01 | 3,12×10^−02 | NA | MAOA |
| TGF-β Signaling | 4,99×10^−01 | 2,06×10^−02 | NA | BMPR2,RALA |
| p53 Signaling | 4,93×10^−01 | 2,04×10^−02 | NA | ADGRB1,TIGAR |
| UVA-Induced MAPK Signaling | 4,93×10^−01 | 2,04×10^−02 | NA | RALA,SMPD3 |
| Sertoli Cell-Sertoli Cell Junction Signaling | 4,92×10^−01 | 1,65×10^−02 | 0,000 | JAM3,RALA,SPTBN1,SPTBN2 |
| Glycerophospholipid biosynthesis | 4,87×10^−01 | 2,02×10^−02 | NA | CDS2,GPCPD1 |
| VEGF Signaling | 4,87×10^−01 | 2,02×10^−02 | NA | RALA,ROCK2 |
| S Phase | 4,81×10^−01 | 2×10^−02 | NA | PSMD1,PSMD6 |
| Cargo concentration in the ER | 4,79×10^−01 | 2,94×10^−02 | NA | GRIA1 |
| Triglyceride metabolism | 4,79×10^−01 | 2,94×10^−02 | NA | PRKACB |
| Fatty Acid β-oxidation I | 4,79×10^−01 | 2,94×10^−02 | NA | ACSBG1 |
| G alpha (q) signalling events | 4,72×10^−01 | 1,72×10^−02 | NA | APP,GRM5,KALRN |
| Phase I - Functionalization of compounds | 4,7×10^−01 | 1,96×10^−02 | NA | CYP46A1,MAOA |
| Transcriptional Regulation by NPAS4 | 4,69×10^−01 | 2,86×10^−02 | NA | IQSEC3 |
| Potassium Channels | 4,65×10^−01 | 1,94×10^−02 | NA | GABBR1,KCND2 |
| DNA Replication Pre-Initiation | 4,59×10^−01 | 1,92×10^−02 | NA | PSMD1,PSMD6 |
| Mouse Embryonic Stem Cell Pluripotency | 4,59×10^−01 | 1,92×10^−02 | NA | BMPR2,RALA |
| Apoptosis Signaling | 4,59×10^−01 | 1,92×10^−02 | NA | PRKCE,RALA |
| IGF-1 Signaling | 4,54×10^−01 | 1,9×10^−02 | NA | PRKACB,RALA |
| MyD88-independent TLR4 cascade | 4,49×10^−01 | 2,7×10^−02 | NA | SARM1 |
| Noradrenaline and Adrenaline Degradation | 4,49×10^−01 | 2,7×10^−02 | NA | MAOA |
| Post-translational protein phosphorylation | 4,44×10^−01 | 1,87×10^−02 | NA | APP,VCAN |
| Glutathione-mediated Detoxification | 4,4×10^−01 | 2,63×10^−02 | NA | GSTO1 |
| Notch Signaling | 4,4×10^−01 | 2,63×10^−02 | NA | MAG |
| Telomerase Signaling | 4,39×10^−01 | 1,85×10^−02 | NA | PTGES3,RALA |
| Antigen Presentation Pathway | 4,31×10^−01 | 2,56×10^−02 | NA | CALR |
| Binding and Uptake of Ligands by Scavenger Receptors | 4,19×10^−01 | 1,79×10^−02 | NA | CALR,HYOU1 |
| Activation of gene expression by SREBF (SREBP) | 4,14×10^−01 | 2,44×10^−02 | NA | FASN |
| Transport of vitamins, nucleosides, and related molecules | 4,14×10^−01 | 2,44×10^−02 | NA | SLC25A4 |
| RET signaling | 4,14×10^−01 | 2,44×10^−02 | NA | PRKACB |
| Antioxidant Action of Vitamin C | 4,1×10^−01 | 1,75×10^−02 | NA | GSTO1,SLC2A3 |
| Formation of WDR5-containing histone-modifying complexes | 4,06×10^−01 | 2,38×10^−02 | NA | OGT |
| Regulation of Actin-based Motility by Rho | 4,05×10^−01 | 1,74×10^−02 | NA | ACTR3,PI4KA |
| IL-13 Signaling Pathway | 4×10^−01 | 1,72×10^−02 | NA | MAOA,ROCK2 |
| Cell Cycle Checkpoints | 3,99×10^−01 | 1,47×10^−02 | 2,000 | CLASP2,DYNC1H1,PSMD1,PSMD6 |
| Aggrephagy | 3,98×10^−01 | 2,33×10^−02 | NA | DYNC1H1 |
| Oncostatin M Signaling | 3,98×10^−01 | 2,33×10^−02 | NA | RALA |
| Xenobiotic Metabolism CAR Signaling Pathway | 3,97×10^−01 | 1,55×10^−02 | NA | GSTO1,PRKCE,PRKCG |
| PAK Signaling | 3,96×10^−01 | 1,71×10^−02 | NA | GIT1,RALA |
| ESR-mediated signaling | 3,92×10^−01 | 1,69×10^−02 | NA | DDX5,PTGES3 |
| SUMOylation of transcription cofactors | 3,91×10^−01 | 2,27×10^−02 | NA | DDX5 |
| TAK1-dependent IKK and NF-kappa-B activation | 3,91×10^−01 | 2,27×10^−02 | NA | APP |
| Regulation of mRNA stability by proteins that bind AU-rich elements | 3,91×10^−01 | 2,27×10^−02 | NA | TNPO1 |
| Signaling by Retinoic Acid | 3,91×10^−01 | 2,27×10^−02 | NA | PDHX |
| TBC/RABGAPs | 3,91×10^−01 | 2,27×10^−02 | NA | TBC1D24 |
| Synthesis of DNA | 3,87×10^−01 | 1,68×10^−02 | NA | PSMD1,PSMD6 |
| Arachidonic acid metabolism | 3,83×10^−01 | 2,22×10^−02 | NA | FAAH |
| Sphingosine-1-phosphate Signaling | 3,83×10^−01 | 1,67×10^−02 | NA | ADCY9,SMPD3 |
| Cytosolic sensors of pathogen-associated DNA | 3,76×10^−01 | 2,17×10^−02 | NA | LRRFIP1 |
| Interleukin-2 family signaling | 3,76×10^−01 | 2,17×10^−02 | NA | SYNJ1 |
| Apelin Pancreas Signaling Pathway | 3,76×10^−01 | 2,17×10^−02 | NA | PRKACB |
| Neuroprotective Role of THOP1 in Alzheimer's Disease | 3,74×10^−01 | 1,64×10^−02 | NA | APP,PRKACB |
| ID1 Signaling Pathway | 3,74×10^−01 | 1,49×10^−02 | NA | APP,BMPR2,RALA |
| LXR/RXR Activation | 3,7×10^−01 | 1,63×10^−02 | NA | ACACA,FASN |
| Sensory perception of taste | 3,69×10^−01 | 2,13×10^−02 | NA | SCN2A |
| Mitotic Prometaphase | 3,67×10^−01 | 1,48×10^−02 | NA | CLASP2,DCTN1,DYNC1H1 |
| Regulation of Insulin-like Growth Factor (IGF) transport and uptake by IGFBPs | 3,66×10^−01 | 1,61×10^−02 | NA | APP,VCAN |
| Role of NANOG in Mammalian Embryonic Stem Cell Pluripotency | 3,66×10^−01 | 1,61×10^−02 | NA | BMPR2,RALA |
| DNA Damage Bypass | 3,62×10^−01 | 2,08×10^−02 | NA | DDB1 |
| TCR signaling | 3,58×10^−01 | 1,59×10^−02 | NA | PSMD1,PSMD6 |
| NR1H2 and NR1H3-mediated signaling | 3,49×10^−01 | 2×10^−02 | NA | FASN |
| Melanoma Signaling | 3,49×10^−01 | 2×10^−02 | NA | RALA |
| MYC Mediated Apoptosis Signaling | 3,49×10^−01 | 2×10^−02 | NA | PRKACB |
| FAT10 Cancer Signaling Pathway | 3,49×10^−01 | 2×10^−02 | NA | BMPR2 |
| Class I MHC mediated antigen processing and presentation | 3,43×10^−01 | 1,33×10^−02 | 1,342 | CALR,CUL5,NEDD4L,PSMD1,PSMD6 |
| Cell surface interactions at the vascular wall | 3,43×10^−01 | 1,42×10^−02 | NA | JAM3,MAG,SLC7A8 |
| TNFR1 Signaling | 3,43×10^−01 | 1,96×10^−02 | NA | MADD |
| Mitotic G1 phase and G1/S transition | 3,39×10^−01 | 1,53×10^−02 | NA | PSMD1,PSMD6 |
| Apoptotic execution phase | 3,37×10^−01 | 1,92×10^−02 | NA | PLEC |
| Gα12/13 Signaling | 3,32×10^−01 | 1,5×10^−02 | NA | RALA,ROCK2 |
| Signaling by TGF-beta Receptor Complex | 3,31×10^−01 | 1,89×10^−02 | NA | NEDD4L |
| Spliceosomal Cycle | 3,31×10^−01 | 1,89×10^−02 | NA | DDX39B |
| Senescence Pathway | 3,28×10^−01 | 1,34×10^−02 | NA | BMPR2,CACNG2,PDHX,RALA |
| Autophagy | 3,26×10^−01 | 1,38×10^−02 | NA | AKT1S1,PRKACB,RALA |
| Phototransduction Pathway | 3,25×10^−01 | 1,85×10^−02 | NA | PRKACB |
| STAT3 Pathway | 3,24×10^−01 | 1,48×10^−02 | NA | BMPR2,RALA |
| Metabolism of non-coding RNA | 3,19×10^−01 | 1,82×10^−02 | NA | WDR77 |
| Deadenylation-dependent mRNA decay | 3,19×10^−01 | 1,82×10^−02 | NA | EIF4A2 |
| Signaling by PTK6 | 3,19×10^−01 | 1,82×10^−02 | NA | CRK |
| NLR signaling pathways | 3,14×10^−01 | 1,79×10^−02 | NA | APP |
| Folate Signaling Pathway | 3,14×10^−01 | 1,79×10^−02 | NA | OGT |
| EGF Signaling | 3,14×10^−01 | 1,79×10^−02 | NA | ITPR1 |
| CNTF Signaling | 3,08×10^−01 | 1,75×10^−02 | NA | RALA |
| Signaling by PDGF | 3,03×10^−01 | 1,72×10^−02 | NA | CRK |
| Cancer Drug Resistance by Drug Efflux | 3,03×10^−01 | 1,72×10^−02 | NA | RALA |
| Polyamine Regulation in Colon Cancer | 2,98×10^−01 | 1,69×10^−02 | NA | AKT1S1 |
| Endometrial Cancer Signaling | 2,93×10^−01 | 1,67×10^−02 | NA | RALA |
| Triacylglycerol Degradation | 2,93×10^−01 | 1,67×10^−02 | NA | FAAH |
| PCP (Planar Cell Polarity) Pathway | 2,93×10^−01 | 1,67×10^−02 | NA | ROCK2 |
| Class A/1 (Rhodopsin-like receptors) | 2,88×10^−01 | 1,26×10^−02 | 0,000 | ADORA1,APP,PLPPR4,PSAP |
| Assembly of collagen fibrils and other multimeric structures | 2,88×10^−01 | 1,64×10^−02 | NA | PLEC |
| Kinesins | 2,88×10^−01 | 1,64×10^−02 | NA | KIF1A |
| Semaphorin Signaling in Neurons | 2,88×10^−01 | 1,64×10^−02 | NA | ROCK2 |
| IL-2 Signaling | 2,83×10^−01 | 1,61×10^−02 | NA | RALA |
| Peroxisomal protein import | 2,79×10^−01 | 1,59×10^−02 | NA | USP9X |
| PTEN Regulation | 2,76×10^−01 | 1,33×10^−02 | NA | PSMD1,PSMD6 |
| NAD Signaling Pathway | 2,73×10^−01 | 1,32×10^−02 | NA | LDHA,RYR2 |
| Transcriptional regulation by RUNX1 | 2,7×10^−01 | 1,32×10^−02 | NA | PSMD1,PSMD6 |
| PXR/RXR Activation | 2,69×10^−01 | 1,54×10^−02 | NA | PRKACB |
| ERB2-ERBB3 Signaling | 2,69×10^−01 | 1,54×10^−02 | NA | RALA |
| Pyridoxal 5'-phosphate Salvage Pathway | 2,69×10^−01 | 1,54×10^−02 | NA | PRKCE |
| Role of PI3K/AKT Signaling in the Pathogenesis of Influenza | 2,65×10^−01 | 1,52×10^−02 | NA | CRK |
| WNT/Ca+ pathway | 2,65×10^−01 | 1,52×10^−02 | NA | CAMK2A |
| Relaxin Signaling | 2,62×10^−01 | 1,29×10^−02 | NA | ADCY9,PRKACB |
| Phase II - Conjugation of compounds | 2,61×10^−01 | 1,49×10^−02 | NA | CNDP2 |
| Role of Pattern Recognition Receptors in Recognition of Bacteria and Viruses | 2,59×10^−01 | 1,28×10^−02 | NA | PRKCE,PRKCG |
| Ovarian Cancer Signaling | 2,54×10^−01 | 1,27×10^−02 | NA | PRKACB,RALA |
| Agrin Interactions at Neuromuscular Junction | 2,52×10^−01 | 1,45×10^−02 | NA | RALA |
| Role of JAK1 and JAK3 in γc Cytokine Signaling | 2,52×10^−01 | 1,45×10^−02 | NA | RALA |
| Superpathway of Melatonin Degradation | 2,52×10^−01 | 1,45×10^−02 | NA | MAOA |
| SPINK1 General Cancer Pathway | 2,52×10^−01 | 1,45×10^−02 | NA | RALA |
| Aryl Hydrocarbon Receptor Signaling | 2,51×10^−01 | 1,26×10^−02 | NA | GSTO1,PTGES3 |
| Microautophagy Signaling Pathway | 2,48×10^−01 | 1,25×10^−02 | NA | PSMD1,PSMD6 |
| HEY1 Signaling Pathway | 2,46×10^−01 | 1,24×10^−02 | NA | ADORA1,BMPR2 |
| ISG15 antiviral mechanism | 2,45×10^−01 | 1,41×10^−02 | NA | EIF4A2 |
| Glioma Invasiveness Signaling | 2,37×10^−01 | 1,37×10^−02 | NA | RALA |
| Serotonin Degradation | 2,37×10^−01 | 1,37×10^−02 | NA | MAOA |
| ERK5 Signaling | 2,33×10^−01 | 1,35×10^−02 | NA | RALA |
| Pre-NOTCH Expression and Processing | 2,26×10^−01 | 1,32×10^−02 | NA | ATP2A2 |
| Hypoxia Signaling in the Cardiovascular System | 2,26×10^−01 | 1,32×10^−02 | NA | LDHA |
| Antiproliferative Role of Somatostatin Receptor 2 | 2,22×10^−01 | 1,3×10^−02 | NA | RALA |
| Signaling by NOTCH1 | 2,19×10^−01 | 1,28×10^−02 | NA | NBEA |
| Neurotrophin/TRK Signaling | 2,19×10^−01 | 1,28×10^−02 | NA | RALA |
| DDX58/IFIH1-mediated induction of interferon-alpha/beta | 2,16×10^−01 | 1,27×10^−02 | NA | APP |
| Signaling by MET | 2,16×10^−01 | 1,27×10^−02 | NA | CRK |
| Maturity Onset Diabetes of Young (MODY) Signaling | 2,16×10^−01 | 1,27×10^−02 | NA | CACNG2 |
| Thyroid Cancer Signaling | 2,16×10^−01 | 1,27×10^−02 | NA | RALA |
| FLT3 Signaling in Hematopoietic Progenitor Cells | 2,06×10^−01 | 1,22×10^−02 | NA | RALA |
| Estrogen-Dependent Breast Cancer Signaling | 2,06×10^−01 | 1,22×10^−02 | NA | RALA |
| JAK/STAT Signaling | 2,06×10^−01 | 1,22×10^−02 | NA | RALA |
| Role of MAPK Signaling in the Pathogenesis of Influenza | 2×10^−01 | 1,19×10^−02 | NA | RALA |
| Pulmonary Fibrosis Idiopathic Signaling Pathway | −0×10^00 | 9,2×10^−03 | NA | BMPR2,RALA,ROCK2 |
| Pyroptosis Signaling Pathway | −0×10^00 | 1,06×10^−02 | NA | PRKACB |
| Wound Healing Signaling Pathway | −0×10^00 | 7,94×10^−03 | NA | BMPR2,RALA |
| Immunogenic Cell Death Signaling Pathway | −0×10^00 | 1,11×10^−02 | NA | CALR |
| Macrophage Classical Activation Signaling Pathway | −0×10^00 | 1,06×10^−02 | NA | ACO2,PDHX |
| Multiple Sclerosis Signaling Pathway | −0×10^00 | 9,01×10^−03 | NA | GRIN1,GRIN2B |
| Pathogen Induced Cytokine Storm Signaling Pathway | −0×10^00 | 5,39×10^−03 | NA | RYR2,SLC2A3 |
| Role of Chondrocytes in Rheumatoid Arthritis Signaling Pathway | −0×10^00 | 7,09×10^−03 | NA | AKT1S1 |
| Role of Osteoblasts in Rheumatoid Arthritis Signaling Pathway | −0×10^00 | 4,1×10^−03 | NA | BMPR2 |
| Role of Osteoclasts in Rheumatoid Arthritis Signaling Pathway | −0×10^00 | 3,25×10^−03 | NA | RALA |
| Ribonucleotide Reductase Signaling Pathway | −0×10^00 | 5,85×10^−03 | NA | AKT1S1 |
| Glucocorticoid Receptor Signaling | −0×10^00 | 1,03×10^−02 | NA | MT-ND4,MT-ND5,NDUFA9,PRKACB,PTGES3,RALA |

| | | | | |
|---|---|---|---|---|
| Natural Killer Cell Signaling | −0×10^00 | 5,05×10^−03 | NA | RALA |
| Macrophage Alternative Activation Signaling Pathway | −0×10^00 | 1,05×10^−02 | NA | ACLY,AKT1S1 |
| Chaperone Mediated Autophagy Signaling Pathway | −0×10^00 | 9,45×10^−03 | 0,816 | APP,ATP6V0A1,ATP6V1D,ATP6V1H,HTT,PEA15 |
| IL-33 Signaling Pathway | −0×10^00 | 5,41×10^−03 | NA | PRKACB |
| Activin Inhibin Signaling Pathway | −0×10^00 | 4,74×10^−03 | NA | ROCK2 |
| LPS/IL-1 Mediated Inhibition of RXR Function | −0×10^00 | 1,17×10^−02 | NA | ACSBG1,GSTO1,MAOA |
| PIP3 activates AKT signaling | −0×10^00 | 7,14×10^−03 | NA | AKT1S1 |
| TCF dependent signaling in response to WNT | −0×10^00 | 1,01×10^−02 | NA | PSMD1,PSMD6 |
| RNA Polymerase II Transcription | −0×10^00 | 6,71×10^−03 | NA | DDX39B |
| Integrin cell surface interactions | −0×10^00 | 1,18×10^−02 | NA | JAM3 |
| Oxidative Stress Induced Senescence | −0×10^00 | 1,16×10^−02 | NA | MINK1 |
| Protein folding | −0×10^00 | 1,02×10^−02 | NA | PFDN5 |
| G alpha (s) signalling events | −0×10^00 | 6,9×10^−03 | NA | ADCY9 |
| Cell junction organization | −0×10^00 | 1,09×10^−02 | NA | PLEC |
| O-linked glycosylation | −0×10^00 | 9,01×10^−03 | NA | THSD7A |
| TP53 Regulates Metabolic Genes | −0×10^00 | 1,14×10^−02 | NA | TIGAR |
| Nucleotide Excision Repair | −0×10^00 | 1,03×10^−02 | NA | DDB1 |
| Interleukin-4 and Interleukin-13 signaling | −0×10^00 | 9,09×10^−03 | NA | MAOA |
| Mitotic Prophase | −0×10^00 | 9,71×10^−03 | NA | USO1 |
| Processing of Capped Intron-Containing Pre-mRNA | −0×10^00 | 6,97×10^−03 | NA | DDX39B,DDX5 |
| Acute Phase Response Signaling | −0×10^00 | 5,41×10^−03 | NA | RALA |
| FXR/RXR Activation | −0×10^00 | 1,07×10^−02 | NA | FASN,GSTO1 |
| BBSome Signaling Pathway | −0×10^00 | 1,02×10^−02 | 2,236 | ADGRB1,ADORA1,GABBR1,GRM5,PRKACB |
| Sleep REM Signaling Pathway | −0×10^00 | 9,01×10^−03 | NA | PRKACB |
| Cohesin Chromatin Regulation Pathway | −0×10^00 | 8,1×10^−03 | NA | DDX5,PCDH1 |
| Histone Modification Signaling Pathway | −0×10^00 | 3,39×10^−03 | NA | HDGFL2 |
| IL-12 Signaling and Production in Macrophages | −0×10^00 | 8,77×10^−03 | NA | PRKCE,PRKCG |
| Role of NFAT in Regulation of the Immune Response | −0×10^00 | 3,87×10^−03 | NA | IPO5,ITPR1,RALA,TNPO1 |
| FcγRIIB Signaling in B Lymphocytes | −0×10^00 | 5,65×10^−03 | NA | CACNG2,ITPR1,RALA |
| CCR5 Signaling in Macrophages | −0×10^00 | 8,02×10^−03 | NA | CACNG2,ITPR1,PRKCE,PRKCG |
| Calcium-induced T Lymphocyte Apoptosis | −0×10^00 | 8,7×10^−03 | 2,000 | ATP2A2,ITPR1,PRKCE,PRKCG |
| IL-17 Signaling | −0×10^00 | 5,35×10^−03 | NA | RALA |
| CD28 Signaling in T Helper Cells | −0×10^00 | 3,87×10^−03 | NA | ACTR3,ITPR1 |
| IL-15 Signaling | −0×10^00 | 3,8×10^−03 | NA | AKT1S1,RALA |
| ICOS-ICOSL Signaling in T Helper Cells | −0×10^00 | 7,89×10^−03 | NA | CAMK2A,CAMK2D,CAMK2G,ITPR1 |
| HMGB1 Signaling | −0×10^00 | 5,99×10^−03 | NA | RALA |
| Germ Cell-Sertoli Cell Junction Signaling | −0×10^00 | 5,88×10^−03 | NA | RALA |
| Prostate Cancer Signaling | −0×10^00 | 8,77×10^−03 | NA | RALA |
| Acute Myeloid Leukemia Signaling | −0×10^00 | 1,1×10^−02 | NA | RALA |
| Bladder Cancer Signaling | −0×10^00 | 8,62×10^−03 | NA | RALA |
| Chronic Myeloid Leukemia Signaling | −0×10^00 | 1,08×10^−02 | NA | AKT1S1,CRK,RALA |
| Production of Nitric Oxide and Reactive Oxygen Species in Macrophages | −0×10^00 | 1,05×10^−02 | NA | PRKCE,PRKCG |
| p70S6K Signaling | −0×10^00 | 5,18×10^−03 | NA | PRKCE,PRKCG,RALA |
| Colorectal Cancer Metastasis Signaling | −0×10^00 | 1,11×10^−02 | NA | ADCY9,PRKACB,RALA |
| Pancreatic Adenocarcinoma Signaling | −0×10^00 | 7,94×10^−03 | NA | RALA |
| Systemic Lupus Erythematosus Signaling | −0×10^00 | 9,38×10^−04 | NA | RALA |
| CDC42 Signaling | −0×10^00 | 1,04×10^−02 | NA | ACTR3,EXOC2,EXOC3,EXOC8,IQGAP2,RALA |
| FAK Signaling | −0×10^00 | 8,7×10^−03 | 1,667 | ACTR3,ADGRB1,ADORA1,BMPR2,CRK,GABBR1,GIT1,GRM5,RALA |
| Hereditary Breast Cancer Signaling | −0×10^00 | 7,25×10^−03 | NA | RALA |
| Role of Osteoblasts, Osteoclasts and Chondrocytes in Rheumatoid Arthritis | −0×10^00 | 4,39×10^−03 | NA | BMPR2 |
| Phospholipase C Signaling | −0×10^00 | 5,36×10^−03 | 1,000 | ADCY9,ARHGEF2,ITPR1,PRKCE,PRKCG,RALA |
| Glioblastoma Multiforme Signaling | −0×10^00 | 1,17×10^−02 | NA | ITPR1,RALA |
| Regulation of IL-2 Expression in Activated and Anergic T Lymphocytes | −0×10^00 | 2,16×10^−03 | NA | RALA |
| NUR77 Signaling in T Lymphocytes | −0×10^00 | 3,9×10^−03 | NA | PRKCE,PRKCG |
| PKCθ Signaling in T Lymphocytes | −0×10^00 | 1,08×10^−02 | 1,000 | CACNG2,CAMK2A,CAMK2D,CAMK2G,ITPR1,RALA |
| PI3K Signaling in B Lymphocytes | −0×10^00 | 8,49×10^−03 | 1,342 | CAMK2A,CAMK2D,CAMK2G,ITPR1,RALA |
| Role of Tissue Factor in Cancer | −0×10^00 | 9,66×10^−03 | NA | BMPR2,RALA |
| Salvage Pathways of Pyrimidine Ribonucleotides | −0×10^00 | 1,03×10^−02 | NA | PRKCE |
| Granulocyte Adhesion and Diapedesis | −0×10^00 | 5,29×10^−03 | NA | JAM3 |
| Regulation of the Epithelial-Mesenchymal Transition Pathway | −0×10^00 | 5,13×10^−03 | NA | RALA |
| TEC Kinase Signaling | −0×10^00 | 3,46×10^−03 | NA | PRKCE,PRKCG |
| Adipogenesis pathway | −0×10^00 | 7,19×10^−03 | NA | BMPR2 |
| HIPPO signaling | −0×10^00 | 1,15×10^−02 | NA | DLG4 |
| SAPK/JNK Signaling | −0×10^00 | 5,98×10^−03 | NA | CRK,MINK1,RALA |
| PI3K/AKT Signaling | −0×10^00 | 1×10^−02 | NA | RALA,SYNJ1 |
| IL-4 Signaling | −0×10^00 | 5,2×10^−03 | NA | AKT1S1,MAOA,RALA |
| B Cell Receptor Signaling | −0×10^00 | 7,89×10^−03 | 0,447 | CAMK2A,CAMK2D,CAMK2G,RALA,SYNJ1 |
| WNT/β-catenin Signaling | −0×10^00 | 5,75×10^−03 | NA | BMPR2 |
| PPAR Signaling | −0×10^00 | 9,35×10^−03 | NA | RALA |
| NF-κB Signaling | −0×10^00 | 5,25×10^−03 | NA | BMPR2,PRKACB,RALA |
| T Cell Receptor Signaling | −0×10^00 | 4,87×10^−03 | NA | AKT1S1,RALA,RASGRP1 |
| IL-6 Signaling | −0×10^00 | 7,75×10^−03 | NA | RALA |
| PD-1, PD-L1 cancer immunotherapy pathway | −0×10^00 | 9,35×10^−03 | NA | RASGRP1 |
| Th1 and Th2 Activation Pathway | −0×10^00 | 5,81×10^−03 | NA | BMPR2 |
| Th2 Pathway | −0×10^00 | 7,3×10^−03 | NA | BMPR2 |
| Osteoarthritis Pathway | −0×10^00 | 4,24×10^−03 | NA | BMPR2 |
| NER (Nucleotide Excision Repair, Enhanced Pathway) | −0×10^00 | 1,1×10^−02 | NA | DDB1 |
| T Cell Exhaustion Signaling Pathway | −0×10^00 | 3,53×10^−03 | NA | BMPR2,RALA |
| Systemic Lupus Erythematosus in T Cell Signaling Pathway | −0×10^00 | 6,22×10^−03 | 1,000 | AKT1S1,ITPR1,RALA,ROCK2 |
| Systemic Lupus Erythematosus in B Cell Signaling Pathway | −0×10^00 | 6,92×10^−03 | 0,447 | PRKCE,PRKCG,RALA,RASGRP1,SYNJ1 |
| HOTAIR Regulatory Pathway | −0×10^00 | 6,13×10^−03 | NA | ROCK2 |
| Regulation of the Epithelial Mesenchymal Transition by Growth Factors Pathway | −0×10^00 | 5,21×10^−03 | NA | RALA |
| Tumor Microenvironment Pathway | −0×10^00 | 1,12×10^−02 | NA | RALA,SLC2A3 |
| MSP-RON Signaling in Cancer Cells Pathway | −0×10^00 | 7,14×10^−03 | NA | RALA |
| MSP-RON Signaling in Macrophages Pathway | −0×10^00 | 8,4×10^−03 | NA | RALA |
