## Supplementary material for "Mouse PAW: reverse-translating the FINGER multimodal lifestyle intervention enhances synaptic plasticity and cognition in adult wild type female mice"

### Results

#### Activity in the IntelliCage

We initially assessed the exploratory activity during the first hour and the subsequent 24 h after the mice were placed in the ICs. The activity was similar between the three groups as determined by the number of visits they made to the corners both during the very first hour (one-way ANOVA, $F_{(2, 33)}$ = 0.08, $p$ = 0.925, **Supplementary Fig. 1A**) and the following 24 h in the ICs (two-way mixed models ANOVA, $F_{(8, 132)}$ = 1.2, $p$ = 0.319 for the two-way interaction group*time, **Supplementary Fig. 1B**). To assess the average pattern of daily activity in the IC we compared the median visit number for each mouse during the light daytime and dark nighttime separately, both during baseline and intervention. We observed that the corner visit activity was not only dependent on the daytime but also on the group and experimental phase (three-way mixed model ANOVA, $F_{(2, 32)}$ = 4.2, $p$ = 0.024 for the three-way interaction, **Supplementary Fig. 1C**). Based on the subsequent post hoc analyses, there were no significant group differences in neither day or night activity during baseline (one-way ANOVA, $F = 0.9$, $p = 0.433$ and $F = 0.4$, $p = 0.678$, respectively). Yet, during the intervention period, there were significant group differences in the daily patterns (one-way ANOVA, Day $F = 11.6$, $p = 0.0002$, Night $F = 4.5$, $p = 0.019$). During the daytime, the PAW group (24.4±7.7 visits) visited corners more actively than the Control (14.6±4.4) and Pharma (15.7±2.8) groups (two-tailed Student's *t*-test with Bonferroni correction, $p = 0.004$ and $p = 0.008$, respectively). On average, the PAW group (62.3±17.0) also visited the corners more than the Control group (41.4±15.1) during nighttime ($p = 0.013$). The nighttime difference between PAW and Pharma (50.5±19.1) was not significant ($p = 0.396$). Furthermore, the Control mice slightly yet significantly reduced their activity during the daytime between baseline (17.5±5.2) and intervention (paired two-tailed Student's *t*-test with Bonferroni correction, $p = 0.012$) as well as their nighttime activity (baseline 66.3±30.0, $p = 0.004$). The PAW group, on the other hand, had a significant increase in daytime visits from baseline (18.3±6.7) to intervention ($p = 0.0003$). PAW group's nighttime activity remained similar from baseline (66.5±19.1) to intervention ($p = 1.00$). Like the Controls, the Pharma group had a reduction in daytime visits from baseline (20.5±4.3, $p = 0.001$), yet their nighttime activity did not significantly change ($p = 0.636$).

Despite the PAW group visiting the corners more often than the two other groups during the intervention period, on average they spent the least time in the corners (two-way mixed models ANOVA, $F_{(2, 33)} = 16.7$, $p < 0.0001$ for the two-way interaction group*time, **Supplementary Fig. 1D**). While the PAW group spent on average 8.2±1.5 s in the corners during the intervention, the Control group spent 14.6±4.7 s and the Pharma group 21.9±6.7 s (post hoc two-tailed Student's *t*-test with Bonferroni correction, $p = 0.002$ and $p < 0.0001$, respectively). Moreover, the difference between the Control and Pharma groups was significant ($p = 0.017$). Compared to the baseline, the PAW mice significantly reduced their time spent in the corners during the intervention phase (baseline 10.4±2.6 s, paired two-tailed Student's *t*-test with Bonferroni correction, $p = 0.033$) whereas the Pharma group increased the duration (baseline 13.0±4.0 s, $p = 0.0003$). There was no significant difference between baseline and intervention for the Control group (baseline 10.2±2.5 s, $p = 0.072$). Neither were there any significant group differences in visit durations at baseline. The overall daily activity patterns of all three groups during the study are visualised in **Supplementary Figure 1E**. It is important to mention that when cognitive training programs were running the PAW group was more active than the two other groups as they needed to actively search for water.
